# Longitudinal characterization of circulating neutrophils uncovers distinct phenotypes associated with disease severity in hospitalized COVID-19 patients

**DOI:** 10.1101/2021.10.04.463121

**Authors:** Thomas J. LaSalle, Anna L. K. Gonye, Samuel S. Freeman, Paulina Kaplonek, Irena Gushterova, Kyle R. Kays, Kasidet Manakongtreecheep, Jessica Tantivit, Maricarmen Rojas-Lopez, Brian C. Russo, Nihaarika Sharma, Molly F. Thomas, Kendall M. Lavin-Parsons, Brendan M. Lilly, Brenna N. Mckaig, Nicole C. Charland, Hargun K. Khanna, Carl L. Lodenstein, Justin D. Margolin, Emily M. Blaum, Paola B. Lirofonis, Abraham Sonny, Roby P. Bhattacharyya, Blair Alden Parry, Marcia B. Goldberg, Galit Alter, Michael R. Filbin, Alexandra Chloe Villani, Nir Hacohen, Moshe Sade-Feldman

## Abstract

Multiple studies have identified an association between neutrophils and COVID-19 disease severity; however, the mechanistic basis of this association remains incompletely understood. Here we collected 781 longitudinal blood samples from 306 hospitalized COVID-19^+^ patients, 78 COVID-19^−^ acute respiratory distress syndrome patients, and 8 healthy controls, and performed bulk RNA-sequencing of enriched neutrophils, plasma proteomics, cfDNA measurements and high throughput antibody profiling assays to investigate the relationship between neutrophil states and disease severity or death. We identified dynamic switches between six distinct neutrophil subtypes using non-negative matrix factorization (NMF) clustering. At days 3 and 7 post-hospitalization, patients with severe disease had an enrichment of a granulocytic myeloid derived suppressor cell-like state gene expression signature, while non-severe patients with resolved disease were enriched for a progenitor-like immature neutrophil state signature. Severe disease was associated with gene sets related to neutrophil degranulation, neutrophil extracellular trap (NET) signatures, distinct metabolic signatures, and enhanced neutrophil activation and generation of reactive oxygen species (ROS). We found that the majority of patients had a transient interferon-stimulated gene signature upon presentation to the emergency department (ED) defined here as Day 0, regardless of disease severity, which persisted only in patients who subsequently died. Humoral responses were identified as potential drivers of neutrophil effector functions, as enhanced antibody-dependent neutrophil phagocytosis and reduced NETosis was associated with elevated SARS-CoV-2-specific IgG1-to-IgA1 ratios in plasma of severe patients who survived. *In vitro* experiments confirmed that while patient-derived IgG antibodies mostly drove neutrophil phagocytosis and ROS production in healthy donor neutrophils, patient-derived IgA antibodies induced a predominant NETosis response. Overall, our study demonstrates neutrophil dysregulation in severe COVID-19 and a potential role for IgA-dominant responses in driving neutrophil effector functions in severe disease and mortality.

## INTRODUCTION

Since December 2019, severe acute respiratory syndrome coronavirus 2 (SARS-CoV-2) has caused over 228 million cases of coronavirus disease (COVID-19) and over 4.6 million deaths globally^1,2^. While our understanding of the molecular and cellular effects of COVID-19 continues to grow, it remains an urgent concern; the disease causes a variety of clinical presentations and results in a myriad of symptoms in non-hospitalized and hospitalized patients. Thus far, many studies of SARS-CoV-2 infections have highlighted the response of the host immune system during the course of COVID-19 cases. These investigations have widely shown that severe COVID-19 patients tend to present with broad immune dysfunction, specifically, decreased lymphocyte counts, abundant inflammation and heightened cytokine levels, delayed B cell activation and antibody production, and impaired interferon-mediated antiviral responses^3–9^. Additionally, current literature describes bulk proteomic and transcriptomic signatures of neutrophil hyperactivation states in severe COVID-19 patients and suggests a unique role of neutrophils in the pathogenesis of the disease^7,10–13^. An overabundance of neutrophil precursors coupled with dysfunctional mature neutrophils in the blood of severe COVID-19 patients compared to mild patients suggests that a dysregulated myeloid cell compartment is indicative of severe disease^11^. Finally, multiple studies have proposed that emergency myelopoiesis, the activation of hematopoietic progenitors in the bone marrow which leads to an abundance of suppressive immature neutrophils, is a prominent feature of severe COVID-19 associated with poor prognosis^11,14,15^.

The observed dysregulation of the humoral immune response in severe COVID-19 patients has implications beyond immediate viral neutralization, as pathogen recognition by neutrophils is partly driven by opsonic receptors such as Fc receptors, and thus neutrophil responses may also be affected^16^. As of yet, the effects of dysregulated humoral responses on neutrophil effector functions is not understood. Current literature suggests that recognition of antigen-antibody complexes by neutrophils may be important in eliciting various effector responses. The removal of pathogens by neutrophils can occur directly in a process termed antibody-dependent neutrophil phagocytosis (ADNP) in which neutrophils recognize pathogen-antibody immune complexes^17,18^, or through NETosis, a specialized cell death program in which neutrophils release neutrophil extracellular traps (NETs) consisting of chromatin modified with anti-microbial proteins^19,20^. In the context of various pathologies such as viral infection, cancer, and heparin-induced thrombocytopenia, NETosis has been shown to be largely driven by antibody-Fc receptor interactions which can be further regulated by antibody isotype and glycosylation profile^21–23^. In regard to the role of NETosis in COVID-19 specifically, limited studies indicate the importance of NETs in COVID-19-associated myocardial infarctions and immunothrombosis^24,25^. Furthermore, protein and transcriptional signatures of neutrophil activation and degranulation in plasma and whole blood of hospitalized COVID-19 patients were predictive of increased mortality^14^.

While most studies on COVID-19 immunity utilize samples lacking polymorphonuclear cells (i.e. peripheral blood mononuclear cells (PBMCs)), there is a need to perform in-depth analyses of the role of neutrophil dynamics in differential COVID-19 progression in large cohorts in order to better understand the role of these cell types within the wider host immune response to SARS-CoV-2. Here, we present a longitudinal study of enriched blood neutrophils from a large cohort of hospitalized COVID-19 patients that combines unbiased, bulk transcriptomic analysis of enriched blood neutrophils with plasma proteomics, cfDNA measurements, and high-throughput antibody profiling in order to understand neutrophil response dynamics during the immune response to SARS-CoV-2 infection.

## RESULTS

### Longitudinal Profiling of Neutrophils from COVID-19 patients

Between March and May 2020, we enrolled 384 patients who presented to the Massachusetts General Hospital Emergency Department (ED) with suspected cases of COVID-19 based on clinical presentation of acute respiratory distress. Subsequently, 306 tested positive for COVID-19. We stratified patient disease acuity into five categories based on the World Health Organization (WHO) COVID-19 ordinal outcome scale as previously described^26^: A1, death within 28 days; A2, intubation, mechanical ventilation, and survival to 28 days; A3, hospitalized and requiring supplemental oxygen; A4, hospitalized without requiring supplemental oxygen; A5, discharged directly from the ED without requiring admission within 28 days. Additionally, A1-A2 were classified as severe, and A3-A5 as non-severe. Primary outcomes for each patient (Acuity_Max_, Severity_Max_) were defined as the most severe disease level with 28 days of enrollment (on the ordinal WHO COVID-19 outcome scale or binary severity scale, respectively; Supplementary Table S1). For each patient, we took blood draws on Days 0 (n=374) upon admission to the ED (likely corresponding to Day 7-8 post infection), 3 (n=212), and 7 (n=143) of all who remained hospitalized during this time, as well as an additional event-driven draw if there was a significant change in the patient’s clinical status (e.g., intubation, removal from ventilator). We collected an additional 8 blood draws from healthy individuals. Using negative selection to enrich for neutrophils, we obtained a total of 781 neutrophil-enriched samples from 388 individuals (Figure S1A). Additionally, we analyzed 1472 unique plasma proteins measured by Proximity Extension Assay using the Olink platform (a dataset that we previously published)^26^, quantified cell-free DNA, and performed high-throughput antibody profiling (partial cohort recently published)^27–29^ (Figure 1A, Method Details). After quality control analysis of the RNA-sequencing data^30^, we retained 698 samples from 370 unique patients for analysis (Figure S1B-H).

**Figure 1.**
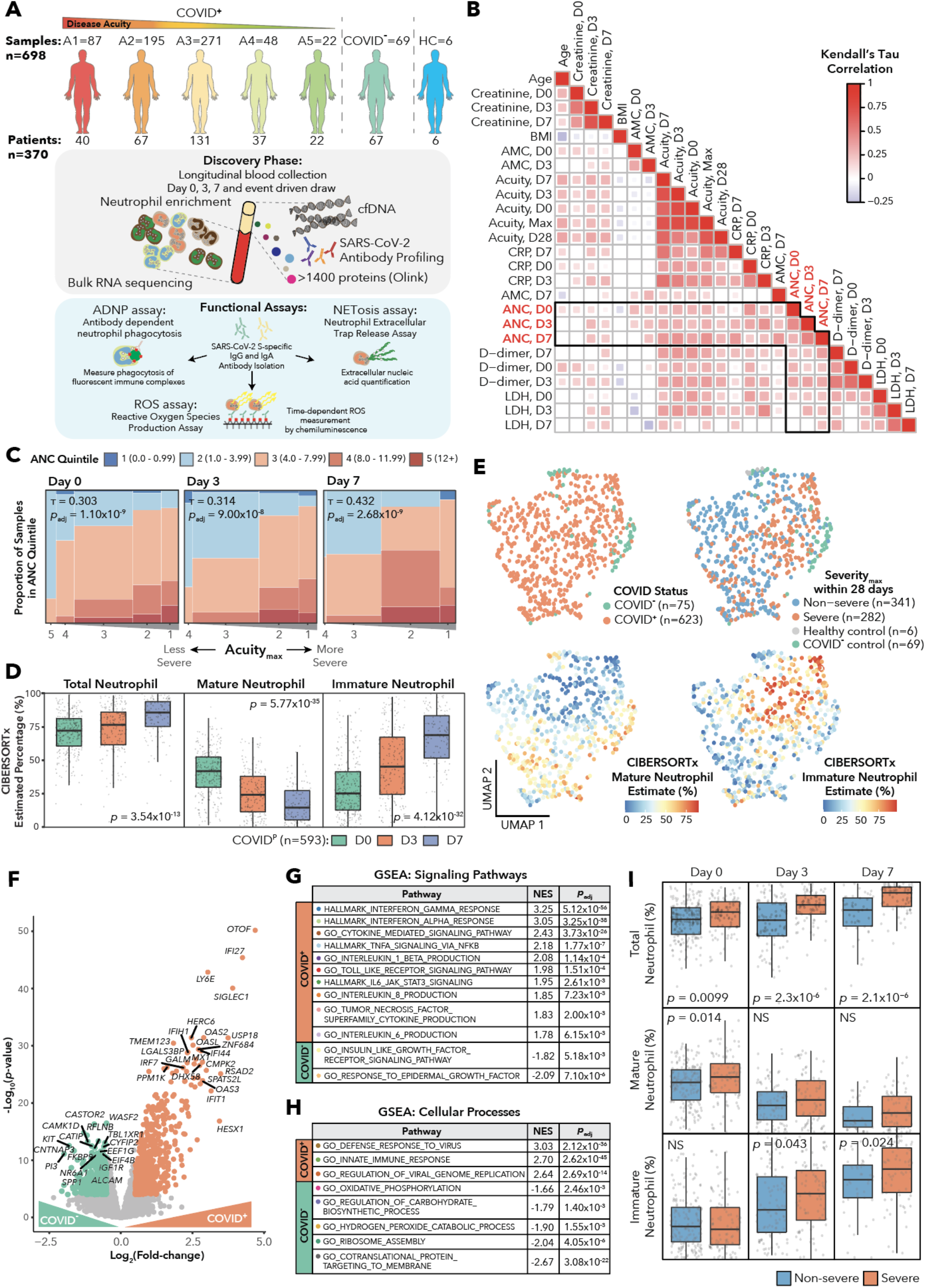
SARS-CoV-2 Infection Induces Transcriptionally Distinct Neutrophil Profiles from COVID-19-negative Respiratory Disease Patients. **(A)** Schematic of cohort and study methodology. **(B)** Kendall’s tau correlation heatmap showing clinical variables that have significant correlations with absolute neutrophil counts on Days 0, 3, or 7 with FDR q < 0.05 in COVID-19-positive patients. **(C)** Mosaic plots displaying the ordinal correlation measured by Kendall’s tau between absolute neutrophil count (ANC) quintile and Acuity_Max_ within 28 days for COVID-19-positive patients. **(D)** Boxplots of Total Neutrophil Percentage, Mature Neutrophil Percentage, and Immature Neutrophil Percentage as estimated by CIBERSORTx across Days 0, 3, and 7 for COVID-19-positive patients. Indicated p values are for the Kruskal-Wallis test. **(E)** UMAP (Uniform Manifold Approximation and Projection) plots of all neutrophil-enriched bulk RNA-seq samples that passed quality control, color-coded by (left to right) COVID-19 status, Severity_Max_ within 28 days, CIBERSORTx Estimated Mature Neutrophil Percentage, and CIBERSORTx Estimated Immature Neutrophil Percentage. **(F)** Volcano plot showing differentially expressed genes between COVID-19-positive and COVID-19-negative patients hospitalized with respiratory disease on Day 0. Color-coded circles indicate genes with log_2_(fold-change) > 0.5 and p < 10^−4^. **(G)**-**(H)** Gene set enrichment analysis for (G) signaling pathways and (H) cellular processes from MSigDB for the differential expression result between COVID-19-positive and COVID-19-negative samples on Day 0 from (F). Tables indicate pathway, normalized enrichment score (NES), and adjusted p-values. **(I)** Box plots comparing CIBERSORTx Estimated Total Neutrophil Percentage, Mature Neutrophil Percentage, and Immature Neutrophil Percentage between severe and non-severe patients on Days 0, 3, and 7. P values are determined using the Wilcoxon rank-sum test. Y-axis runs from 0 to 100 percent.

### COVID-19 induces a strong interferon response signature in neutrophils followed by an expansion of immature neutrophils

Based on previous studies that identified associations between circulating neutrophil counts and COVID-19 disease severity^31–36^, we analyzed ordinal correlations between clinically-obtained absolute neutrophil count (ANC) quintile and clinical parameters associated with disease severity. We observed positive correlations between ANC and creatinine, LDH, CRP, and D-dimer, consistent with the known role of neutrophils in inflammation and thrombosis^11,37^ (Figure 1B, Supplementary Table S1). There were particularly strong correlations between ANC and both CRP and D-dimer at Days 0 and 3. Additionally, we found robust ordinal correlations between ANC and acuity (and accordingly, intubation) that increased from Day 0 upon presentation to the ED (corresponding to approximately day 7-8 of infection) to Day 7 (Figure 1C).

Since 100% neutrophil purity could not be guaranteed following enrichment for bulk RNA-seq, we next sought to determine the cell type composition of our samples using CIBERSORTx^38^. We used the COVID-19 Bonn Cohort 2 fresh whole blood single-cell dataset as a single-cell reference for the deconvolution of our bulk data^11^ (Supplementary Table S1). We generated cell-type-specific signatures and used these to estimate fractions of mature neutrophils, immature neutrophils, monocytes, T and NK cells (together denoted T/NK), B cells, and plasmablasts for each sample (Figure S2). We defined total neutrophils as the sum of the mature and immature neutrophil fractions. We found that lower estimated total neutrophil content was associated with lower ANC (Fisher’s test, *p* = 1.2×10^−17^) obtained clinically, and lower estimated T/NK fraction was associated with lower absolute lymphocyte count (ALC, Fisher’s test, *p* = 0.0129), indicating that the total neutrophil fraction reflected the abundance of neutrophils in patients’ blood (Figure S3). Overall, CIBERSORTx estimated that a mean of ∼75% of cells per sample are attributed to neutrophils (Fig S2D-E). Among COVID-19 patients, the estimated total neutrophil fraction increased from Day 0 to Day 7 (Figure 1D), driven largely by the expansion of immature neutrophils, which increased over time, while the mature neutrophil fraction decreased. Indeed, immature neutrophils have been reported to expand in time with COVID-19 infection^14^. In agreement with the correlations between ANC counts and clinical variables, we observed an association between intubation status and total neutrophil fraction at Day 3 (Figure S4). This trend was more significant at Day 7 (Figure S4), with the fraction of immature but not mature neutrophils significantly associated with intubation status on Day 7.

Uniform manifold approximation and projection (UMAP) visualization of bulk RNA-seq samples revealed distinct groupings based on disease status and immature/mature neutrophil fraction (Figure 1E and Figure S5A-C) with COVID-19-negative samples largely grouped together. Among the COVID-19 samples, the landscape was defined by a gradient of mature to immature neutrophil percentages that overlapped with mild and severe disease, respectively.

Next, to analyze differentially expressed genes and programs that were induced during COVID-19 infection with respiratory distress, we performed differential expression analysis and gene set enrichment analysis (GSEA) between COVID-19-positive and similarly symptomatic COVID-19-negative respiratory disease patients on Day 0, excluding healthy controls (Figure 1F-H, Supplementary Table S1). In order to correct for contamination from non-neutrophil cell types, we added CIBERSORTx cell type fractions as covariates to the differential expression model. We used the CIBERSORTx fractions for total neutrophils, monocytes, T/NK cells, plasmablasts, and an additional immunoglobulin score (Figure S6; Method Details). GSEA revealed strong anti-viral signatures enriched in COVID-19-positive samples, such as response to IFNγ and IFN-α, TLR signaling, and cytokine production (TNFα, IL1β, IL8) (Figure 1G-H). Relative to COVID-19-positive, the COVID-19-negative samples showed higher levels of response to epidermal growth factor (EGF).

Finally, to identify neutrophil expression correlates of COVID-19 disease severity, we began by comparing CIBERSORTx cell type fractions across severe and non-severe patients (Figure 1I). The total neutrophil fraction (sum of mature and immature fractions) was significantly elevated in severe patients across all time points, consistent with our earlier observation of elevated ANC in severe disease. The elevation of total neutrophil fraction in severe COVID-19 patients on Day 0 was driven by a higher proportion of mature neutrophils, while on Days 3 and 7 the difference was attributed to a higher proportion of immature neutrophils (Figure 1I). Furthermore, in agreement with studies demonstrating lymphopenia in severe COVID-19 patients^39^, we found higher fractions of T/NK cells in non-severe patients across all time points (Figure S5D).

### Unbiased NMF clustering defines neutrophil states in the context of SARS-CoV-2 infection

We next sought to identify bulk neutrophil gene expression subtypes and their associations with disease outcomes using Bayesian non-negative matrix factorization (NMF) clustering^40^ (Method Details). We clustered all samples that had CIBERSORTx total neutrophil fraction above 50% (n = 635, 91% of samples) to reduce artifacts of cell type contamination, and identified six robust bulk neutrophil subtypes (Figure 2A, Figure S7, Supplementary Table S2). We denote samples with less than 50% neutrophil content as Neu-Lo and note that this group had significantly lower C-reactive protein (CRP) on Day 0 and were significantly less likely to come from patients requiring intubation (Figure S8).

**Figure 2.**
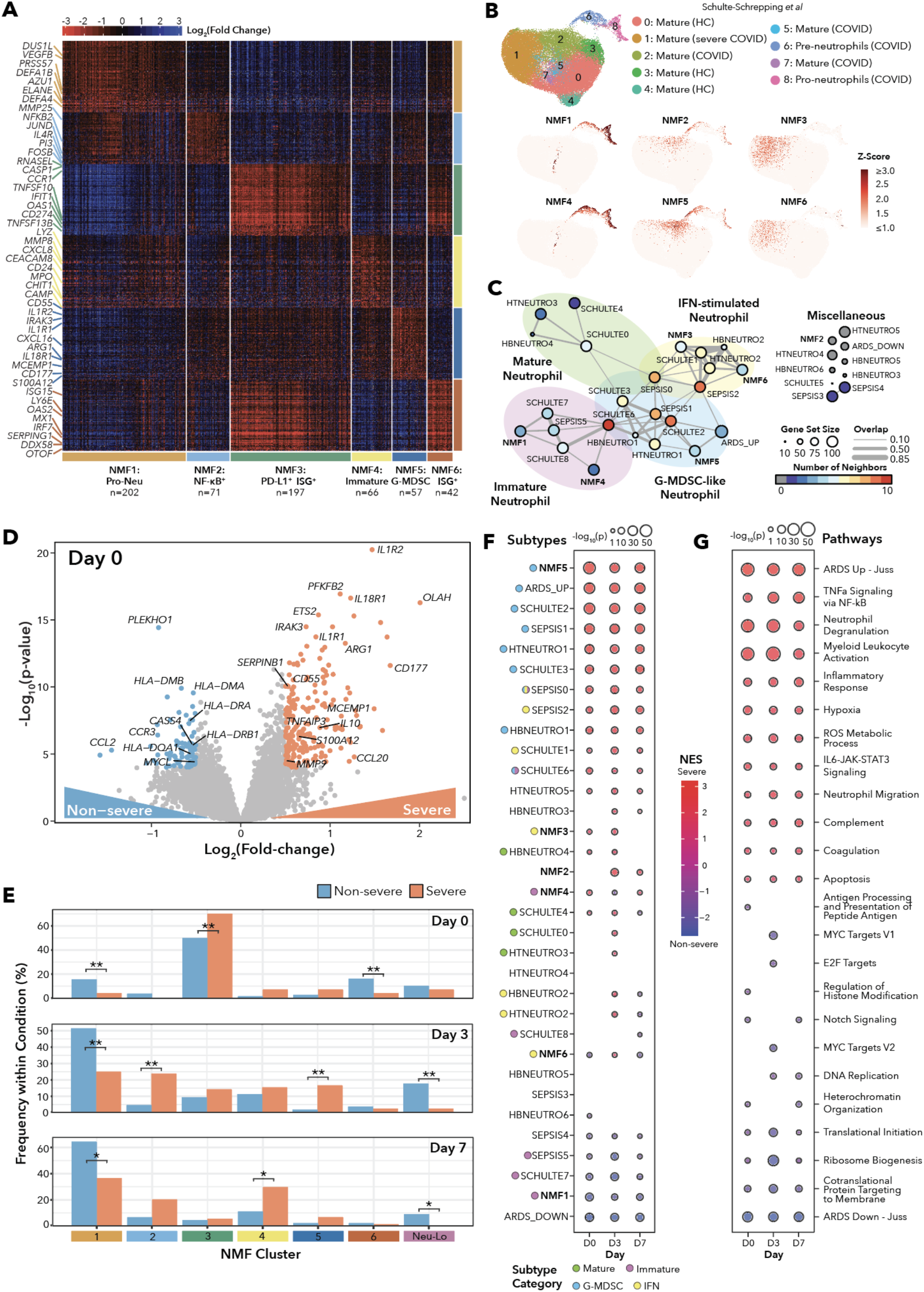
Severe Outcomes Are Associated with Transitions Between Neutrophil States in COVID-19 Patients. **(A)** Unbiased NMF clustering of neutrophil-enriched bulk RNA-seq data for samples with CIBERSORTx Estimated Total Neutrophil Percentage greater than 50%. The six mRNA expression subtypes are denoted as NMF1 (Pro-Neu), NMF2 (NF-kB^+^), NMF3 (PD-L1^+^ISG^+^), NMF4 (Immature Activated), NMF5 (G-MDSC), and NMF6 (ISG^+^). **(B)** UMAPs of single-cell RNA-seq data of fresh whole-blood neutrophils from COVID-19 positive patients and healthy controls from Schulte-Schrepping *et al*. Cohort 2. UMAPs are color-coded by Louvain clustering (top) and NMF cluster metagene scores (bottom) from each NMF cluster’s marker genes in (B). HC; healthy control. **(C)** Network diagram displaying relationships between NMF subtype marker gene lists and previously-defined neutrophil signature gene lists from RNA-seq data. Edges are drawn between nodes with Jaccard index > 0.05 between gene sets. Node radius is proportional to the number of genes in each gene set, edge width indicates the overlap coefficient between gene sets, and node color indicates number of other signatures with shared genes. Gene sets include fresh whole blood single-cell neutrophils from Schulte-Schrepping *et al*. Cohort 2, single-cell neutrophils from tumors and blood of lung cancer patients from Zilionis *et al*., single-cell neutrophils from sepsis patients from Reyes *et al*., and bulk neutrophils from COVID-19-negative ARDS patients and healthy controls from Juss *et al*. Gene sets have been grouped according to their connections in the network into IFN-stimulated, Mature, G-MDSC-like, and Immature Neutrophils. Gene sets with Jaccard index < 0.05 with all other gene sets are categorized as Miscellaneous. **(D)** Volcano plot showing differentially expressed genes between COVID-19-positive severe and non-severe patients on Day 0. Color-coded points indicate genes with log_2_(fold-change) > 0.5 and p < 10^−4^. **(E)** Bar plots showing the proportion of COVID-19-positive samples with membership in each NMF cluster. Bar heights are normalized to the number of COVID-19-positive samples within a particular severity for each day. Single asterisk indicates FDR q < 0.05, double asterisk indicates FDR q < 0.01 according to Fisher’s Exact Test within each day, with multiple hypothesis correction across all days. On Day 0, 161 out of 283 samples were classified as NMF3, and a higher proportion of severe patients were in this category (*p_adj_* = 6.35 x 10^−3^), while more non-severe samples were classified as NMF1 (*p_adj_* = 9.99 x 10^−3^) or NMF6 (*p_adj_* = 9.99 x 10^−3^). On Day 3, the majority of non-severe samples (74 out of 107) were classified as NMF1 or Neu-Lo. There was enrichment for severe samples in NMF2 (*p_adj_* = 2.25 x 10^−3^) and NMF5 (*p_adj_* = 2.25 x 10^−3^). Finally, on Day 7, the majority of non-severe samples (29 out of 45) remained in NMF1, and there was enrichment for non-severe samples in NMF1 (*p_adj_* = 0.0114) and Neu-Lo (*p_adj_* = 0.0438). NMF4 was enriched for severe samples (*p_adj_* = 0.0495). **(F)-(G)** Gene set enrichment analysis for the differentially expressed genes between COVID-19-positive severe and non-severe patients on Days 0, 3, and 7. Bubble size is scaled to −log_10_(p-value) and color corresponds to normalized enrichment score (NES). Neutrophil subtype gene set enrichment is shown in (F), while pathway enrichment is shown in (G).

Two subtypes (NMF3 and NMF6) were characterized by high expression of interferon-stimulated genes (ISGs). NMF3 included inflammatory caspases (*CASP1*, *CASP4*, *CASP5*), Fc receptors (*FCGR3A*, *FCGR3B*, *FCGR1CP)*, and complement receptor *C3AR1* expression. Clinically, samples from NMF3 had significantly higher CRP on Day 0 and had a higher fraction of samples from patients requiring intubation as compared to all other clusters (Figure S8). On the other hand, NMF6 had high levels of several granzymes (*GZMB, GZMH, GZMM*) and a non-overlapping set of ISGs. Consistent with these granzyme marker genes, the estimated proportion of T/NK cells was highest in NMF6 compared to the other subtypes, potentially indicating higher levels of non-neutrophil contamination than other subtypes (Figure S7F).

NMF1 and NMF4 were composed of predominantly immature neutrophils. NMF1 was enriched for ribosomal genes, components of neutrophil granules (*ELANE*, *AZU1*, *DEFA1B*, *DEFA4*), and protective antioxidant potential (*VEGFB*), suggestive of a neutrophil-progenitor-like state, while NMF4 had a more activated (*CEACAM8*/CD66b, *CD24*), chemoattractive (*CXCL8*), and potentially tissue-damaging (*MPO*, *CHIT1*, *MMP8*, *LYZ*) profile. On Day 7, the immature neutrophil clusters showed significant differences in D-dimer; NMF1 had lower levels while NMF4 was skewed towards higher D-dimer, potentially implicating NMF4 in thrombosis (Figure S8). To probe the differences between the types of immature neutrophils, we performed differential expression analysis and GSEA on samples classified as NMF1 versus NMF4 (Figure S9A-D, Supplementary Table S2). NMF4 samples showed a strong enrichment of neutrophil degranulation, as well as an enrichment of metabolic pathways responsible for generation of reactive oxygen species (ROS) needed for degranulation including the tricarboxylic acid (TCA) cycle, fatty acid metabolism, glycolysis, and peroxisome organization. In contrast, NMF1 samples showed an enrichment for the electron transport chain pathway and oxidative phosphorylation for the production of ATP, suggesting that though these pro-neutrophils are activated, they may be generating energy stores for further differentiation rather than accumulating energy to perform effector functions^41^.

Finally, NMF2 and NMF5 shared transcriptional similarities with myeloid-derived suppressor cells (MDSCs). NMF2 displayed strong NF-kB activation (*NFKB2*, *BCL3*) and *MMP25* expression, while NMF5 showed a robust granulocytic MDSC gene expression signature (*ARG1, CD177, MCEMP1, S100A12*), interferon receptor expression (*IFNGR1*), and IL1B signaling (*IL1R1, IL1R2, IL1RAP*). On Day 3, NMF2 and NMF5 had significantly higher fractions of samples from patients who required intubation during their hospitalization, and in addition, NMF2 had higher CRP and LDH in plasma (Figure S8).

Our NMF-derived clusters were similar to those previously identified by single-cell RNA sequencing (scRNA-seq) in a smaller cohort^11^, and notably, none of the neutrophil NMF gene signatures mapped to single-cell clusters which were composed of healthy controls (Figure 2B, Figure S9E). Additionally, to better contextualize the neutrophil state transcriptional signatures identified from our NMF clustering, we built a network displaying the relationships between our NMF signatures and previously-defined neutrophil gene signatures (Figure 2C, Figure S9F). To define these gene sets, we utilized the NMF cluster marker genes alongside the following signatures derived from neutrophil transcriptomics data in various disease contexts including COVID-19^11^, cancer^42^, sepsis^43^, and non-COVID-19 acute respiratory distress syndrome (ARDS)^44^. By examining the overlap of the NMF marker genes with these previously-defined neutrophil transcriptional signatures, we found that multiple signatures across studies shared several genes, suggesting that these NMF signatures may represent universal neutrophil subtypes.

### Transcriptionally-distinct neutrophil states are associated with COVID-19 disease severity

To identify neutrophil states, genes, and pathways associated with COVID-19 severity, we performed differential gene expression analysis between severe and non-severe COVID-19 patients for each time point, correcting for the confounding effects of cell type composition from CIBERSORTx (Figure 2D, Figure S10A, Supplementary Table S2). To interpret these results and understand the evolution of neutrophil subtypes, we first explored how neutrophil NMF cluster membership varied across disease severity and over time in COVID-19 samples (Figure 2E, Figure S10B-C), and second, we performed GSEA using the neutrophil gene signatures (Figure 2F). While on Day 0 the majority of samples were classified as NMF3 (PD-L1^+^ ISG^+^), patients who progressed in disease severity (Severity_Max_ = severe) were significantly enriched for this antiviral response cluster (as found in the Schulte-Schrepping data when the cutoff for early/late disease is moved from day 10 to day 11 (Figure S10D-E)). On Day 3, the severe samples were more evenly distributed across NMF clusters 1 to 5, with NMF2 (NF-kB^+^) and NMF5 (G-MDSC) significantly enriched for severe samples. In agreement with our NMF clustering results, the signature most strongly enriched in severe patients by GSEA was NMF5 (G-MDSC) across all three time points (Figure 2F). Finally, on Day 7, NMF4 (Immature Activated) was enriched for severe samples.

Unlike the severe samples which were enriched for activated and inflammatory neutrophils, non-severe samples were enriched for NMF1 (Pro-Neu) (all days) and NMF6 (ISG+) (Day 0). The GSEA results also indicated NMF1 (Pro-Neu) as the most enriched consistently neutrophil signature in non-severe samples. In addition, non-severe samples had higher frequencies of Neu-Lo (Days 3 and 7), indicating a resolution of neutrophil activation and reduction of neutrophils in circulation.

We next performed gene- and pathway-level analyses using GSEA (Figure 2G, Supplementary Table S2). Across all three days, the pathways most highly enriched in severe patients included neutrophil degranulation, hypoxia, TNFa signaling via NF-kB, ROS metabolic processes, and neutrophil migration (Figure 2G). Many of the top genes enriched in severe patients across Days 0, 3, and 7 are involved in IL1β signaling (*IL1R1*, *IL1R2*) and neutrophil degranulation (*ARG1*, *CD177*, *MCEMP1*). MHC II genes (*HLA-DMA*, *HLA-DMB*, *HLA-DRA*, *HLA-DRB1*,*HLA-DQA1*) were strongly associated with non-severe disease, as was observed for monocytes in other COVID-19 studies^43,45^. Of note, the two gene sets “ARDS Up - Juss” and “ARDS Down - Juss” were consistently significantly enriched pathways in severe and non-severe patients, respectively, serving as positive control gene sets and suggesting that COVID-19 ARDS and non-COVID-19 ARDS affect neutrophils in similar ways (Figure S11). We also utilized our time course data to search for genes and pathways with diverging expression patterns across time between severe and non-severe patients. Using the likelihood ratio test (LRT) with DESeq2 (Method Details), we tested for an interaction between Day and Severity_Max_ (Figure S12). On the gene level, *SERPINB2*, a gene involved in Th1/Th2 modulation during lentiviral infections, increased with time in severe patients but slightly decreased in non-severe patients (LRT, *q* = 3.3×10^−5^) ^46^. *ZBTB16*, a glucocorticoid response negative feedback gene, was expressed at a higher level on Day 0 in severe patients but the expression decreased at later time points compared to non-severe patients (LRT, *q* = 3.8×10^−8^). On the pathway level, granulocyte chemotaxis remained high in severe patients, but the pathway metagene score decreased with time in non-severe patients (GSEA, *q* = 3.3×10^−3^). Furthermore, the TNFa signaling via NF-kB metagene score increased with time in severe patients but stayed constant in non-severe patients (GSEA, *q* = 3.7x10^−15^). These pathway results are in agreement with the neutrophil subtype analysis, highlighting the role of neutrophil activation in severe disease.

### Neutrophil states are among the most powerful predictors of COVID-19 disease severity as early as Day 0 of hospitalization

Based on the strong enrichments of neutrophil subtype gene signatures in the GSEA for disease severity, we hypothesized that neutrophil subtype scores could add predictive power to models of disease severity upon patient presentation to the ED. To test this hypothesis, we began by assigning each sample a metagene score for the six neutrophil NMF subtypes and the ARDS Up and Down gene sets. We observed that the NMF5 (G-MDSC) signature stratified the patients by acuity category on Day 0 (Figure 3A). Next, we built a series of three nested logistic regression models for predicting Severity_Max_ following SARS-CoV-2 infection (Figure 3B, Figure S13A-C). The first (Model 1) included only patient characteristics, the second (Model 2) added the clinical laboratory values, and the third (Model 3) incorporated the neutrophil gene set scores (Method Details, Supplementary Table S3). Adding the neutrophil subtypes in Model 3 resulted in a marked improvement (AUC: 0.9601, 95% CI: 0.9383 – 0.9819, LRT Model 3 vs. Model 2 *p* = 7.93×10^−6^), demonstrating that neutrophil subtypes may add information to predictive models of COVID-19 severity that is not captured by clinical or laboratory values.

**Figure 3.**
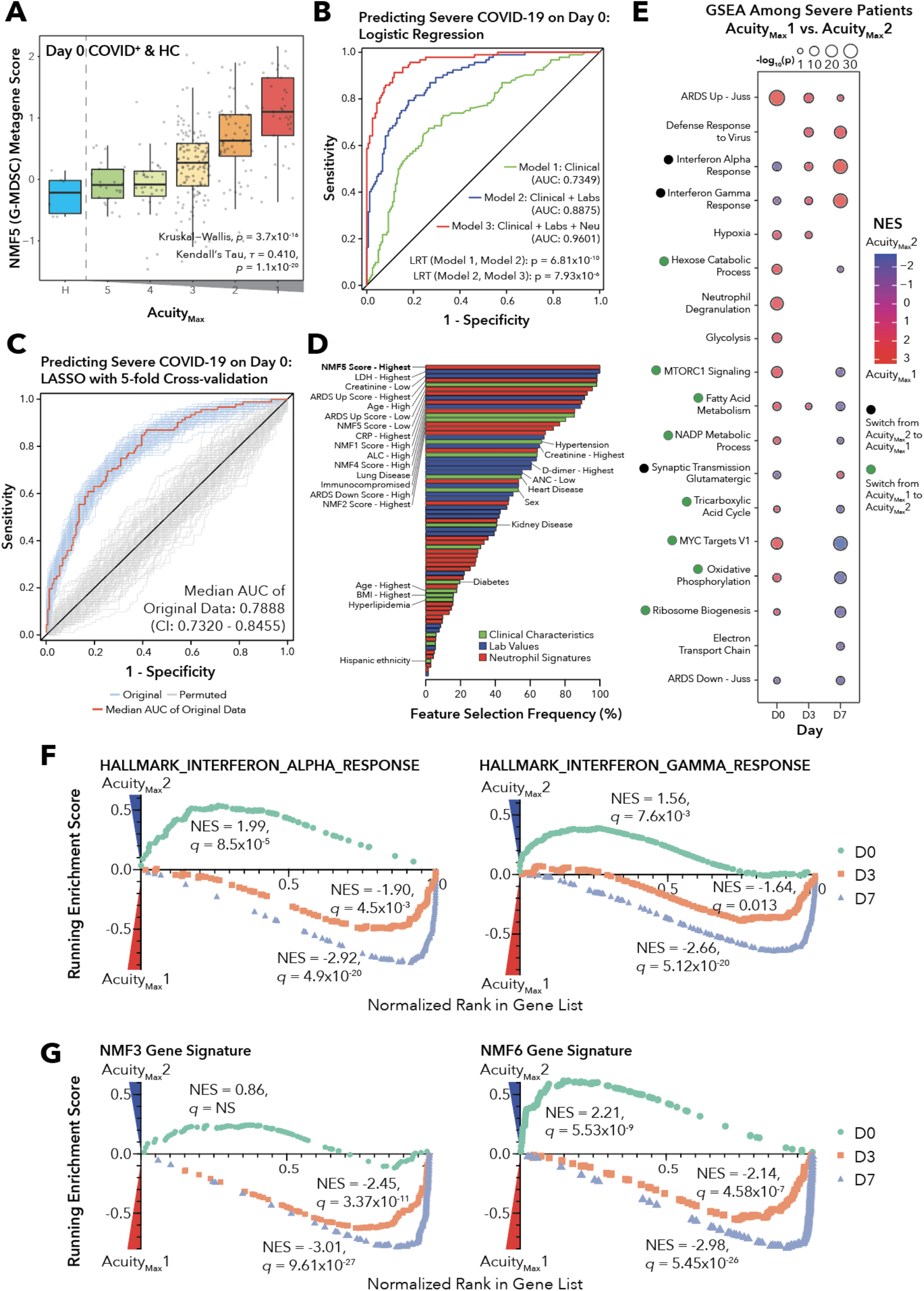
Neutrophil Metabolism and Dysregulated Interferon Signaling Are Associated with Disease Severity and Acuity Following SARS-CoV-2 Infection. **(A)** Box plots displaying the NMF5 (G-MDSC) metagene score for COVID-19-positive samples on Day 0 grouped by Acuity_Max_ and healthy controls. P values and correlations determined with the Kruskal-Wallis test and Kendall’s Tau test. **(B)** Receiver operating characteristic (ROC) curve for predictive performance of logistic regression models predicting COVID-19 disease severity on Day 0. Model 1 includes only clinical characteristics: age, sex, ethnicity, heart disease, diabetes, hypertension, hyperlipidemia, lung disease, kidney disease, immunocompromised status, BMI (AUC: 0.7349, 95% CI: 0.6708 – 0.7991). Model 2 adds the following clinical laboratory values: ANC, ALC, Creatinine, CRP, D-dimer, LDH (AUC: 0.8875, 95% CI: 0.8478 – 0.9273). Model 3 incorporates the following neutrophil gene signature scores, broken into quintiles: NMF1, NMF2, NMF3, NMF4, NMF5, NMF6, ARDS Up - Juss, ARDS Down - Juss (AUC: 0.9601, 95% CI: 0.9383 – 0.9819). Significance of improvement of model determined with the likelihood ratio test. **(C)** ROC curve of predictive performance of a least absolute shrinkage and selection operator (LASSO) model of COVID-19 disease severity on Day 0. Prediction was performed with repeated 5-fold cross-validation with 100 repeats for both the original data and permutated labels of severity. Shown in red is the ROC curve for the cross-validation repeat with the median AUC across all repeats. **(D)** Bar plot displaying the selection frequency for each factor in the LASSO regression model. Bars are color coded by variable type, corresponding to the three models shown in (D). Lab values and gene signatures variables are broken into quintiles with levels: 1 = lowest, 2 = low, 3 = mid, 4 = high, 5 = highest. **(E)** Gene set enrichment analysis for differentially expressed genes between COVID-19-positive patients with Acuity_Max_ of 1 (death) or Acuity_Max_ of 2 (intubation, survival) across Days 0, 3, and 7. Black dots indicate a switch in enrichment from Acuity_Max_2 to Acuity_Max_1, and green dots indicate a switch in enrichment from Acuity_Max_1 to Acuity_Max_2. **(F)** GSEA enrichment plots for the HALLMARK_RESPONSE_TO_INTERFERON_ALPHA (left) and HALLMARK_RESPONSE_TO_INTERFERON_GAMMA (right) gene sets with genes ranked on differential expression between COVID-19-positive patients with Acuity_Max_ 1 and Acuity_Max_ 2 across Days 0, 3, and 7. **(G)** GSEA enrichment plots for the NMF3 gene signature (left) and the NMF6 gene signature (right) with genes ranked on differential expression between COVID-19-positive patients with Acuity_Max_ 1 and Acuity_Max_ 2 across Days 0, 3, and 7.

Given the improvement of Model 3 over Model 2, we next sought to determine which subset of features from Models 1-3 were most important for predicting disease severity and whether these included neutrophil gene signatures. To this end, we performed feature selection using a least absolute shrinkage and selection operator (LASSO) logistic regression model of COVID-19 disease severity on Day 0 (Figure 3C, Method Details). Across all 100 five-fold repeats of cross-validation, the two features which were included by the model every time were the highest NMF5:G-MDSC score quintile and the highest LDH quintile, indicating that the LASSO logistic regression model selected these two features as strong predictors of severe disease for COVID-19 patients on Day 0 of hospitalization (Figure 3D, Supplementary Table S3).

### Longitudinal analyses reveal diverging pathway dynamics between survivors and non-survivors among severe patients

To test whether any neutrophil genes or pathways could be used to predict survival of the most severe patients upon intubation, we performed differential gene expression analysis (Figure S13D-F, Supplementary Table S3) and GSEA (Figure 3E) between samples of Acuity_Max_1 (death within 28 days) and Acuity_Max_2 (intubated but survived). On Day 0, the most significantly enriched pathways in patients who died were the Juss et al. non-COVID-19 ARDS neutrophil signature (*p_adj_* = 7.7×10^−26^, NES = −3.12) and neutrophil degranulation (*p_adj_* = 1.1×10^−15^, NES = −2.13). Interestingly, we observed that several metabolic signaling pathways switched from being enriched at Day 0 in patients who died to being enriched in patients who survived at Day 7. On Day 0, the interferon alpha response and interferon gamma response pathways were enriched in patients who survived, but on Days 3 and 7, the signature became more enriched in patients who died, with stronger enrichment on Day 7 (Figure 3F). Recent work has shown that interferon signaling is delayed or dysregulated in patients infected with SARS-CoV-2, and a late burst of interferon activity and immune cell activation once the lungs are already highly infiltrated has been suggested to contribute to fatal outcomes^5^. In accordance with the interferon response signatures, we also observed that the enrichment of the NMF3 (PD-L1+ISG+) and NMF6 (ISG+) signatures switched from patients who survived on Day 0 to patients who died on Days 3 and 7 (Figure 3G, Figure S13G). Interestingly, the metabolic pathways followed the opposite trend; on Day 0, many of the key metabolic pathways distinguishing NMF1 (Pro-Neu) and NMF4 (Immature Activated) such as fatty acid metabolism, NADP metabolic process, tricarboxylic acid cycle, and MTORC1 signaling were enriched in patients who died within 28 days, but on Day 7, these pathways were enriched in patients who survived intubation. Though NMF cluster membership was not associated with death or survival among severe patients (Figure S13H), the GSEA results suggest that the metabolic differences underlying the NMF clustering are associated with survival.

### NETosis is implicated in severe COVID-19 pathology through transcriptomics, proteomics, and circulating cell-free DNA

Several studies have reported associations between SARS-CoV-2 infection and NETs^47–49^. NETs are extracellular webs of chromatin that contain highly reactive oxidative enzymes such as myeloperoxidase which have the ability to capture and neutralize microbes including viruses^21^, but in excess can cause tissue damage and microvascular thrombosis^50^. To look for a transcriptomic signature of virally-induced NETosis and its associations with disease outcomes, we defined a NETosis metagene score composed of *PADI4*, *MPO*, *ELANE*, *TNF*, *CXCL8*, *GSDMD*, and *TLR3* genes (Figure S14A-E), encoding protein components of NETs, signaling inducers and the viral double-stranded (ds)DNA-sensing receptor *TLR3* (Figure S14A). Our score correlated strongly with a previously-defined NETosis gene signature (Figure S14B)^51^. The analysis revealed a significant enrichment of the NETosis metagene score on Days 3 and 7 in severe patients compared to non-severe patients (Figure 4A). Additionally, across NMF clusters, we found higher NETosis scores in the immature clusters NMF1 (Pro-Neu) and NMF4 (Immature Activated) (Figure 4A).

**Figure 4.**
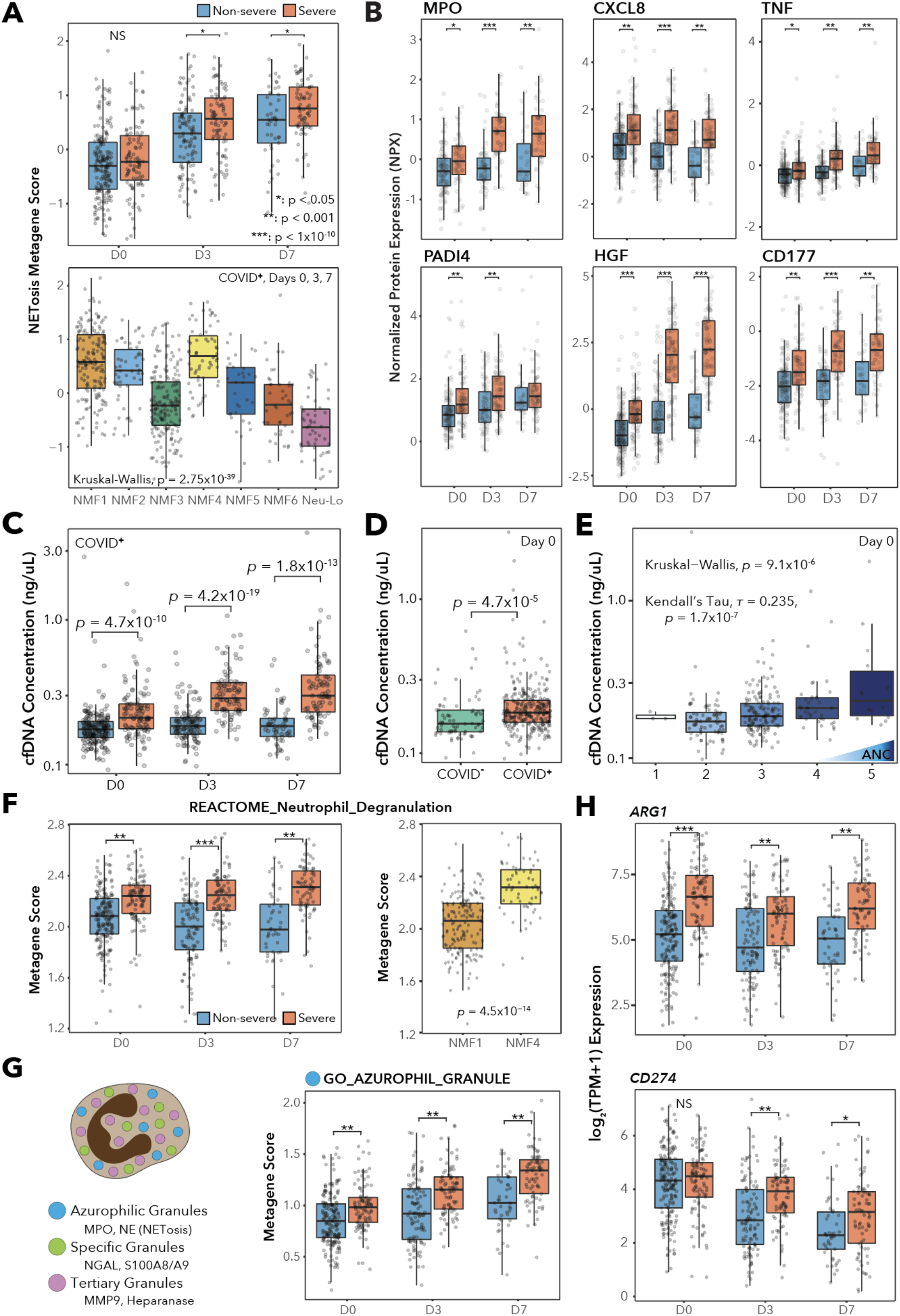
Transcriptomics, Proteomics, and cfDNA Analyses Identify Neutrophil Effector Function Signatures Associated with Severe COVID-19 Outcomes. **(A)** Box plots of NETosis metagene score (*PADI4*, *MPO*, *ELANE*, *TNF*, *CXCL8*, *GSDMD*, *TLR3*) over time split by Severity_Max_ (top), and across NMF clusters (bottom) for COVID-19-positive samples taken on Days 0, 3, and 7. P values indicate Wilcoxon rank-sum test and Kruskal-Wallis test, respectively. **(B)** Box plots of Olink plasma proteomics data Normalized Protein Expression (NPX) values over time for select proteins implicated in NETosis, split by Severity_Max_. Asterisks indicate significance according to the Wilcoxon rank-sum test corresponding to the legend in (A). Samples which do not meet Olink’s quality control criteria for a given protein are not shown. **(C)-(E)** Box plots of cell-free DNA concentration (ng/uL) measured by the Qubit assay, arranged by (C) Day and Severity_Max_, (D) COVID-19 status, and (E) absolute neutrophil count. P values indicate the (C)-(E) Wilcoxon rank-sum test and (E only) Kendall’s tau correlation test. **(F)** Box plots of the pathway metagene score for REACTOME_NEUTROPHIL_DEGRANULATION, arranged by Day and Severity_Max_ (left) and NMF1 vs. NMF4 membership (right) for COVID-19-positive samples on Days 0, 3, and 7. Significance is determined with the Wilcoxon rank-sum test. **(G)** Cartoon depicting the three types of neutrophil granules and their characteristics, accompanied by the pathway metagene score for GO_AZUROPHIL_GRANULE arranged by day and Severity_Max_. P value indicates the Wilcoxon rank-sum test. **(H)** Box plots of log_2_(TPM+1) expression versus time and Severity_Max_ of *ARG1* and *CD274* (PD-L1), two genes implicated in T-cell suppression. P values are for the Wilcoxon rank-sum test.

Though the transcriptional analysis of NETosis revealed an association between an expression signature of NETosis and disease severity, many factors promoting NETosis, such as histone modification, are post-transcriptional^52^ and would not be captured by RNA-seq. Therefore, we next searched for protein markers of NETosis in the matched plasma sample proteomic dataset^26^. Several known protein markers of NETosis were found to be significantly associated with severe disease across all time points and were measured at different levels across neutrophil subtypes, including MPO, CXCL8, TNF, PADI4, HGF, and CD177 (Figure 4B, Figure S14F). In addition to protein markers, we measured levels of circulating cell-free DNA (cfDNA) in the plasma using the Qubit double-stranded DNA high sensitivity Assay. Though the assay measures cfDNA concentration regardless of the source, recent cfDNA methylation studies in COVID-19 have traced one of the major sources of cfDNA to neutrophils^37^. Cell-free DNA concentration was significantly associated with COVID-19 status and disease severity across all time points, and was also correlated with ANC; however, we did not observe significant differences between Acuity_Max_1 and Acuity_Max_2 patients (Figure 4C-E, Figure S15A-C). Furthermore, cell-free DNA was elevated in the plasma from neutrophil samples classified as NMF4 (Immature Activated) versus NMF1 (Pro-Neu), suggesting that NMF4 neutrophils may release greater amounts of NETs (Figure S15B).

### Neutrophil degranulation and neutrophil-mediated T cell suppression are associated with disease severity and are distinguishing features of neutrophil NMF subtypes

Uncontrolled neutrophil degranulation can cause significant tissue damage and is commonly implicated in pathologic inflammatory conditions such as sepsis, acute lung injury, and rheumatoid arthritis^53^. Our GSEA of both severe versus non-severe disease and intubated survivors versus non-survivors revealed associations between neutrophil degranulation enrichment and worse outcomes of COVID-19 (Figure 2G, Figure 3E). Therefore, we defined a neutrophil degranulation metagene score based on the REACTOME_Neutrophil_Degranulation gene set. As expected, the metagene was highly enriched in severe patients versus non-severe patients across all time points, though it was only enriched in Acuity_Max_1 patients over Acuity_Max_2 patients on Day 0 (Figure 4F, Figure S16A). Of note, the neutrophil degranulation metagene score was highly enriched in the NMF4 (Immature Activated) subtype over the NMF1 (Pro-Neu) subtype, identifying a key distinguishing feature of the two immature neutrophil subtypes and supporting the possibility that NMF4 neutrophils may be more capable of executing effector functions (Figure S16B). Additionally, metagene scores for each of the three different types of neutrophil granules (azurophilic, specific, tertiary) based on Gene Ontology gene sets were enriched in severe compared to non-severe patients at all time points (Figure 4G, Figure S16C-D).

Neutrophils have also been shown to suppress T cell activation and proliferation, with some studies demonstrating suppressive neutrophil functions activated only in the final stages of neutrophil differentiation^54^. Therefore, we investigated the associations between neutrophil genes involved in T cell suppression and severity or neutrophil NMF subtype. *ARG1*, which contributes to T cell suppression by depleting L-arginine, was consistently enriched in severe patients (Figure 4H) and had the highest expression in NMF5 (G-MDSC) and NMF4 (Immature Activated) (Figure S16E). *CD274*, the gene encoding PD-L1 which suppresses T cells through engagement with PD-1, had enriched expression in severe patients on Days 3 and 7 (Figure 4H) and was most highly expressed in NMF3 (PD-L1^+^ ISG^+^) (Figure S16F). Notably, NMF1 (Pro-Neu) showed low expression of both genes, consistent with the previous finding that early progenitor neutrophils do not display MDSC functionality^54^. Other MDSC genes involved in T cell suppression were associated with severity on individual days but were less consistent (Figure S16G-I).

### Antibody isotype profiles are major drivers of neutrophil effector functions in COVID-19

Neutrophils function as phagocytes and are able to engulf antibody-opsonized pathogens in an Fc receptor-dependent manner^55^. As such, antibodies play an important role in triggering neutrophil effector functions. Therefore, we utilized systems serology framework^56^ on matched longitudinal plasma samples to explore antibody associations with disease severity and neutrophil phenotypes. We measured the levels of various antibody isotypes and subclasses (IgG1, IgG2, IgG3, IgG4, IgA1, IgM) for multiple SARS-CoV-2 antigens (S, S1, S2, N, RBD) and non-SARS-CoV-2 viral antigens as recently published^57^ (coronavirus OC43, influenza HA, cytomegalovirus CMV) (Supplementary Table S4). We found significantly higher levels of SARS-CoV-2 S-specific IgA1 antibodies in severe patients than non-severe patients on Day 7 (Figure 5A), though there was no difference in SARS-CoV-2 S-specific IgG1 antibodies (Figure S17A). Though we observed few significant associations between antibody types and severity across time points (Figure S17B), there were several associations between acuity level and antibody profiles on Days 3 and 7 among the most severe patients. We found significantly higher IgG1, IgG2, and IgG3 antibodies for a variety of SARS-CoV-2 antigens in Acuity_Max_2 patients on Day 3 as compared to Acuity_Max_1 patients (Figure 5B). On Day 7, IgG1 antibodies for all five SARS-CoV-2 antigens were significantly higher in Acuity_Max_2 patients versus Acuity_Max_1, consistent with previous publications linking delayed or diminished humoral responses to fatal COVID-19 outcomes^9^. We also observed associations between SARS-CoV-2 S-specific IgA1 antibodies and neutrophil NMF states, as well as ANC (Figure S17C-D).

**Figure 5.**
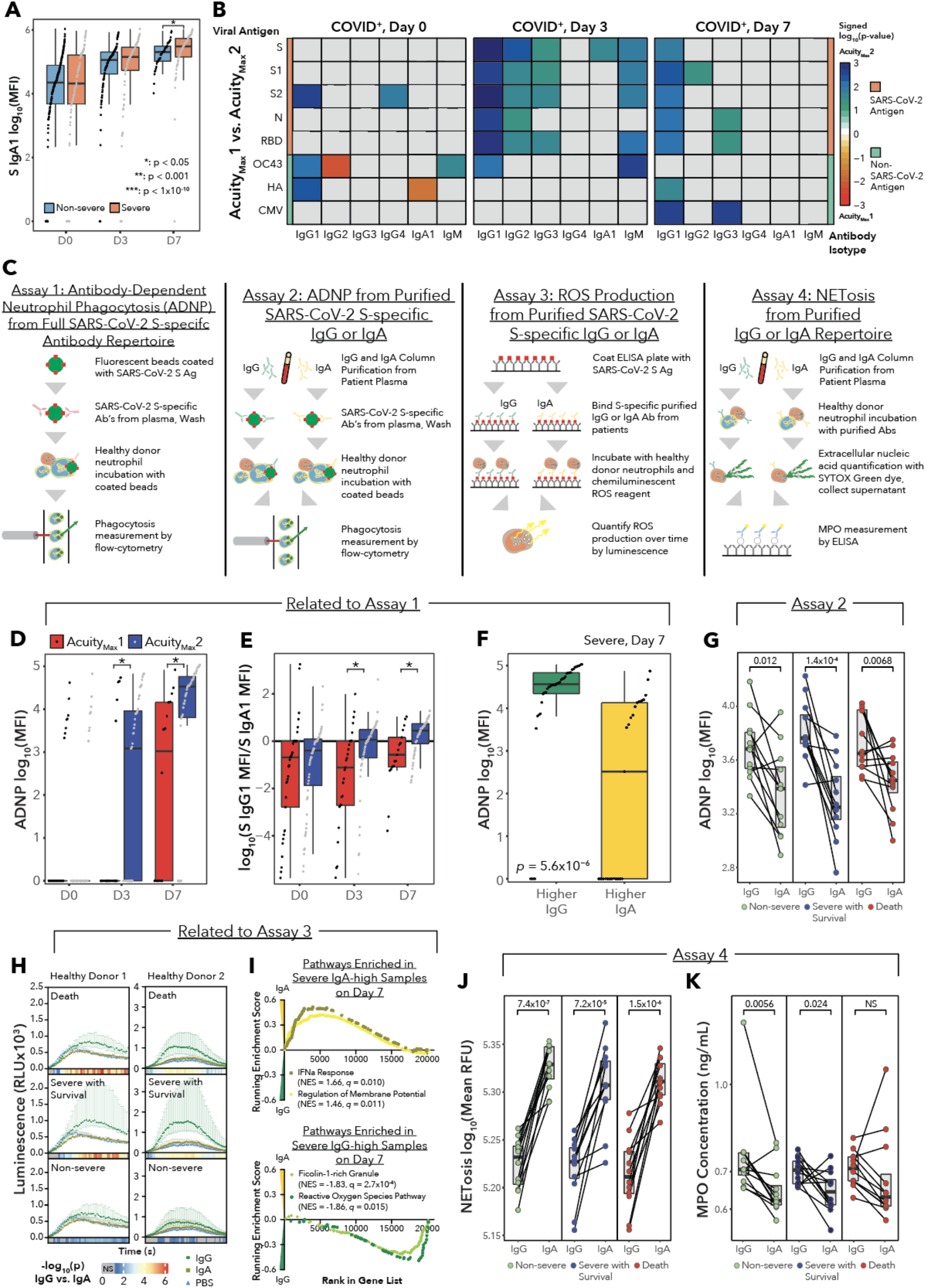
Antibody Profiles are Major Drivers of Neutrophil Function. **(A)** Box plots comparing SARS-CoV-2 spike protein-specific IgA1 log_10_(MFI) values in plasma across time, separated by Severity_Max_. Indicated p values for Wilcoxon rank-sum test. **(B)** Heatmaps displaying the signed (according to fold-change), −log_10_-transformed p values for the Wilcoxon rank-sum tests comparing levels of antigen-specific antibody isotypes between Acuity_Max_1 and Acuity_Max_2 for COVID-19-positive samples. The three main columns indicate Day. Within each heatmap, rows indicate viral antigens, color-coded by virus of origin: SARS-CoV-2 spike protein (S), SARS-CoV-2 spike protein S1 (S1), SARS-CoV-2 spike protein S2 (S2), SARS-CoV-2 nucleocapsid (N), SARS-CoV-2 receptor-binding domain (RBD), Human coronavirus OC43 (OC43), influenza hemagglutinin (HA), and cytomegalovirus (CMV). Within each heatmap, columns indicate antibody isotypes: IgG1, IgG2, IgG3, IgG4, IgA1, IgM. **(C)** Schematics for functional assays. **(D)** Box plot depicting the background-corrected antibody-dependent neutrophil phagocytosis (ADNP) assay log_10_(MFI) score for severe intubated patients, separated by Acuity_Max_ and Day, Acuity_Max_ of 1 (death) and Acuity_Max_of 2 (intubation, survival). P values are for the Wilcoxon rank-sum test. Points are ordered and equally spaced along the x-axis within Day and Acuity according to increasing ADNP values. **(E)** Box plot depicting the log_10_ ratio of the spike (S) protein-specific IgG1 MFI to the S-specific IgA1 MFI for severe intubated patients, separated by Acuity_Max_ and Day. Positive values indicate a ratio in favor of IgG1, while negative values indicate a ratio in favor of IgA1. P values are for the Wilcoxon rank-sum test. **(F)** Box plots of background-corrected ADNP log_10_(MFI) values for severe patients on Day 7, separated by IgG to IgA ratios. P values are for the Wilcoxon rank-sum test. **(G)** Paired-line graphs of ADNP log_10_(MFI) values showing the effect of isolated SARS-CoV-2 S-specific IgG or IgA antibodies from serum of patients who died (n = 12), patients with severe disease who survived (n = 12), and patients with non-severe disease (n = 12). P values are for the Wilcoxon rank-sum test. ADNP log_10_(MFI) values shown are the mean of ADNP values for neutrophils isolated from two separate healthy donors. **(H)** Point-range plots showing the luminescence of the reactive oxygen species reagent, luminol, over time when neutrophils from two healthy donors are exposed to IgG:S or IgA:S immune complexes using the same purified IgG and IgA antibodies as (G) or PBS. Point ranges are plotted as median +/− interquartile range. Color bar beneath each plot displays the −log_10_-transformed P values for the Wilcoxon rank-sum test between IgG and IgA values at each time point, with gray values indicating no significant difference. **(I)** GSEA enrichment plots for pathways enriched between samples with higher IgA:IgG or higher IgG:IgA ratios for COVID-19-positive samples from severe patients on Day 7. Pathways enriched in IgA-high samples are HALLMARK_INTERFERON_ALPHA_RESPONSE and GO_REGULATION_OF_MEMBRANE_POTENTIAL, and pathways enriched in IgG-high samples are HALLMARK_REACTIVE_OXYGEN_SPECIES_PATHWAY and GO_FICOLIN_1_RICH_GRANULE. **(J)** Paired-line graphs of mean SYTOX Green Nucleic Acid Stain log_10_(RFU) (quantification of NETosis) from neutrophils from two healthy donors when exposed to free antibodies from the same batch of IgG or IgA purification as in (G). P values are for the Wilcoxon rank-sum test. **(K)** Paired-line graphs of free MPO concentration as determined by ELISA from the supernatants of neutrophils from healthy donor 1 exposed to IgG or IgA from the NETosis assay in (J). P values are for the Wilcoxon rank-sum test.

To test whether patients’ antibody profiles impact neutrophil phagocytosis, we performed an Antibody-Dependent Neutrophil Phagocytosis (ADNP) assay^58^ (Figure 5C, Supplementary Table S4, Method Details). Though we did not observe any associations between ADNP and severity (Figure S17E), we did find significantly higher ADNP in Acuity_Max_2 patients over Acuity_Max_1 patients on Days 3 and 7 (Figure 5D). Decreased phagocytic activity in patients who died could indicate an inability of neutrophils to clear clots or debris from blood^59^. To understand why ADNP levels were divergent between the two groups, we evaluated differences in the antibody repertoire using the isotype measurements. We found that on Day 0, the majority of severe patients had higher SARS-CoV-2 S-specific IgA1 titers compared to S-specific IgG1; however, over time, the intubated survivors eventually shifted towards higher S-specific IgG1, whereas the non-survivors maintained higher S-specific IgA1 titers (Wilcoxon rank-sum test, D0: NS, D3: *p* = 0.0058, D7: *p* = 0.0090, Figure 5E). We did not observe this trend when only comparing severe and non-severe disease (Figure S17F). We then further stratified samples into two categories: higher S IgG1 titer or higher S IgA1 titer. Among severe patients on Day 7, ADNP was significantly elevated in the higher S IgG1 group (Figure 5F) and the same trend was found across all samples (Figure S17G).

Due to their unique FcR repertoire, including FcγRIIa (CD32a), FcγRIIIb (CD16b), FcγRI (CD64), and FcαRI (CD89), neutrophils can rapidly respond to antibody-complexes, leading to their degradation and clearance^60^. Depending on the FcRs triggered, distinct inflammatory consequences can evolve. Recent studies have demonstrated that while IgG antibodies can induce neutrophil phagocytosis, IgA:virus immune complexes are potent inducers of NETosis^21^. The two antibody isotypes interact with neutrophils through different receptors, with IgA binding FcαR and IgG binding FcγR. In addition to isotype and subclass, changes in Fc-glycosylation at conserved sites on the antibody heavy chain can alter antibody interactions with FcRs^61^. Therefore, we sought to determine whether neutrophil effector functions were differentially impacted by the plasma IgG-to-IgA ratio or whether antibodies from various COVID-19 severities differently modulate neutrophil functions. Thus, we separately purified the IgG and IgA fractions from severe COVID-19 survivors and non-survivors, as well as non-severe patients (*n* = 12 for each group), and performed ADNP, ROS generation, and NETosis assays (Figure 5C, Supplementary Table S4).

For the isotype-specific ADNP experiment, we generated IgG:S (SARS-CoV-2 spike protein) and IgA:S immune complexes and incubated them separately with healthy donor neutrophils to allow for phagocytosis. In all three disease categories (Non-severe, Severe Survivors, Death), only IgG:S immune complexes robustly triggered ADNP (Figure 5G, Figure S18A). Next, we incubated healthy donor neutrophils with immune complexes of both isotypes in the presence of a chemiluminescent reagent and measured the ROS production by neutrophils as a function of time. Across all three disease categories, IgG:S immune complexes induced higher ROS generation than IgA:S (Figure 5H). Notably, IgG:S immune complexes from severe survivors induced significantly higher ROS production than the Non-severe group (Figure S18B). This may be related to distinct IgG glycosylation patterns in severe COVID-19 patients, which could further promote more inflammatory response^62^. To cross-validate these experimental findings, we performed GSEA on neutrophil RNA-seq samples from severe patients on Day 7 comparing patients with higher IgA1:IgG1 ratios to patients with higher IgG1:IgA1 ratios. We found an enrichment for the ROS Pathway and the Ficolin-1-rich Granule pathway in samples with higher IgG1:IgA1 ratios, consistent with the findings of the ROS release assay (Figure 5I). In addition, we found that the IFNα response and Regulation of Membrane Potential pathways were enriched in samples with higher IgA1:IgG1 ratios. Changes in membrane potential are associated with many components of neutrophil activation such as chemotaxis and NETosis^63^.

Finally, we tested whether free IgA or IgG antibodies from patient serum could trigger NETosis, thereby potentially causing microvascular thrombosis among the most severe patients with higher IgA1:IgG1 ratios. We incubated healthy donor neutrophils with free IgA or IgG antibodies from the same purifications as the ADNP and ROS assays, in the presence of a fluorescent cell-impermeable nucleic acid dye (Sytox Green), to quantify the amount of DNA released by neutrophils in the form of NETs. We found strikingly higher NETosis from healthy neutrophils incubated with IgA than IgG, regardless of the disease severity of the patient from which the antibodies were isolated (Figure 5J, Figure S18C). This observation could contribute to the understanding of why severe patients with unresolved IgA1:IgG1 ratios in plasma were less likely to survive intubation. In addition to quantifying NETosis, we collected the supernatant from the assay and performed an ELISA for MPO. In the Non-severe and Severe Survivor groups, we observed significantly less MPO in the presence of IgA than the IgG (Figure 5K). This result is consistent with the ROS assay results which demonstrated higher ROS production after stimulation with purified IgG, as the function of MPO is to catalyze the formation of ROS. However, numerous other factors in the plasma can trigger NETosis in addition to antibodies, including complement and cytokines^64^, and these effects cannot be excluded.

### Plasma proteomics identifies neutrophil-driven secreted proteins and potential ligand-receptor interactions driving phenotypes

To further understand the role of neutrophils in COVID-19 in relation to other blood and immune cells, we analyzed the plasma proteome using our existing Olink Proximity Extension Assay dataset for this cohort. We began by searching for protein markers which were significantly and uniquely associated with neutrophil NMF clusters (Figure 6A, Figure S19, Supplementary Table S5). NMF5 (G-MDSC) in particular had strong upregulation of markers of disease severity and neutrophil activation such as S100A12, HGF, IL1RL1, IL1R2, DEFA1/1B, PADI4, and TGFB1 (Figure 6B). Of note, TGF-β has been shown to influence B cells to class-switch to IgA when stimulated with LPS in vitro^65^. NMF4 (Immature) had the highest levels of ACE2 potentially indicating tissue damage, while NMF3 (PD-L1^+^ ISG^+^) unsurprisingly showed enrichment for IFNL1, CXCL10, and IFNG, likely explaining the phenotype.

**Figure 6.**
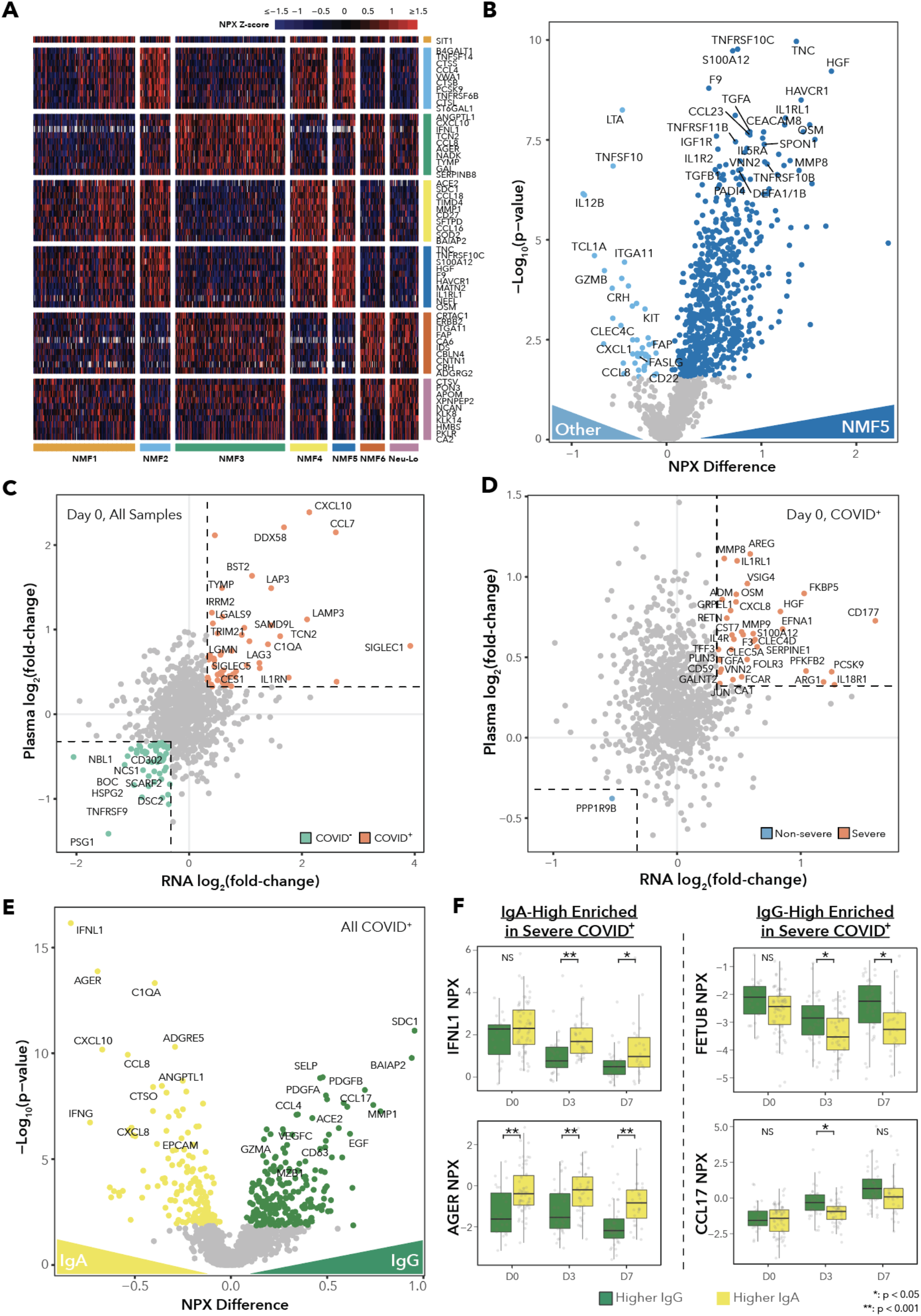
Alterations in the Plasma Proteome are Associated with Neutrophil Subtypes and Antibody Profiles. **(A)** Heatmap displaying scaled expression values for enriched protein markers for each neutrophil NMF cluster for all COVID-19-positive samples. **(B)** Volcano plot showing differentially expressed proteins in matched plasma based on NPX values between neutrophils in NMF5 versus all other clusters for all COVID-19-positive samples. Color-coded points indicate FDR q values < 0.05. **(C)-(D)** Scatter plot comparing the log_2_(fold-change) values for neutrophil RNA-Seq with the log_2_(fold-change) of the NPX difference of the corresponding protein in the plasma between (C) COVID-19-positive and COVID-19-negative patients on Day 0 or (D) COVID-19-positive severe and non-severe patients on Day 0. Differential expression analyses corrected for the clinical covariates of age, sex, ethnicity, heart disease, diabetes, hypertension, hyperlipidemia, pulmonary disease, kidney disease, immunocompromised status. Color-coded points indicate log_2_(fold-change) > 1.25 in both mRNA and protein. **(E)** Volcano plot showing differentially expressed proteins in matched plasma samples based on NPX values between samples with higher IgA:IgG or higher IgG:IgA ratios for all COVID-19-positive samples. Color-coded points indicate FDR q values < 0.05. **(F)** Box plots of NPX values for selected plasma proteins showing significant associations with IgG:IgA ratio for severe COVID-19-positive patients across Days 0, 3, and 7, separated by IgG:IgA ratio. P values are for the Wilcoxon rank-sum test.

Next, we sought to determine which severity-associated proteins in the plasma likely originated from neutrophils. To this end, we compared the log_2_(fold change) values between severe and non-severe patients on each separate time point on both the RNA and protein levels (Figure 6C-D, Figure S20A-B). For both protein and RNA analyses, we included age, sex, ethnicity, heart disease, diabetes, hypertension, hyperlipidemia, pulmonary disease, kidney disease, immuno-compromised status as covariates to control for confounding (and CIBERSORTx cell type fractions for the RNA). We identified several components of neutrophil granules (CD177, MMP8, MMP9, ARG1, S100A12, TGFA), factors involved in clotting (F3, SERPINE1), chemoattraction (CXCL8, IL4R), and inflammation/activation (FKBP5, FCAR, IL18R1, CLEC4D) upregulated in severe disease in both data types, suggesting that neutrophils are key contributors to the severity-associated plasma proteome.

Next, we searched for plasma proteins which were differentially expressed between patients with higher titers of either IgG or IgA to gain insight into the factors associated with antibody isotype switching in COVID-19 patients (Figure 6E-F, Figure S20C-E, Supplementary Table S5). The top protein associated with higher IgA titers was IFNL1. While no study to our knowledge has linked IFN-λ signaling with B cell isotype-switching to IgA, IFN-λ signaling is mainly targeted to epithelial cells in the same way that IgA antibodies are typically found at mucosal surfaces rather than in plasma^66^. Higher IgA titers were also associated with high plasma levels of soluble AGER (also known as RAGE), which is consistent with a recent publication showing significant associations between increased plasma sRAGE, disease severity and mortality in patients infected with SARS-CoV-2^67^. Many other plasma proteins associated with COVID-19 disease severity were enriched in the IgA-high samples, such as IFNG, CXCL10, and CXCL8. On the other hand, within severe samples, IgG-high samples were enriched for FETUB, a protein involved in fatty acid metabolism that can suppress inflammation and which has been shown to be depleted in severe COVID-19^68^. Additionally, IgG-high samples were enriched for CCL17, a Th2 chemokine which may be involved in the activation of class-switch recombination^69^.

Finally, we sought to determine whether any other soluble proteins could be responsible for driving neutrophil phenotypes or disease severity. We performed a ligand-receptor interaction analysis between plasma ligands and receptors differentially expressed by neutrophils between the NMF clusters (Figure 7A), and we tested the relationship between ligand/receptor pairs and disease statuses, separated by day (Figure 7B, Figure S21, Supplementary Table S5). Briefly, we identified differentially expressed ligands from plasma proteomics and differentially expressed neutrophil receptors from RNA-seq and scored each interaction by day based on the number of samples which showed expression of both ligand and receptor above the mean expression. Then we associated each ligand with the inferred cell type of origin using publicly available COVID-19 bronchoalveolar lavage fluid single-cell RNA-seq data^7^ (Figure S22; Method Details). This analysis highlights neutrophil receptor-ligand interactions which are enriched in each NMF subtype and severity group.

**Figure 7.**
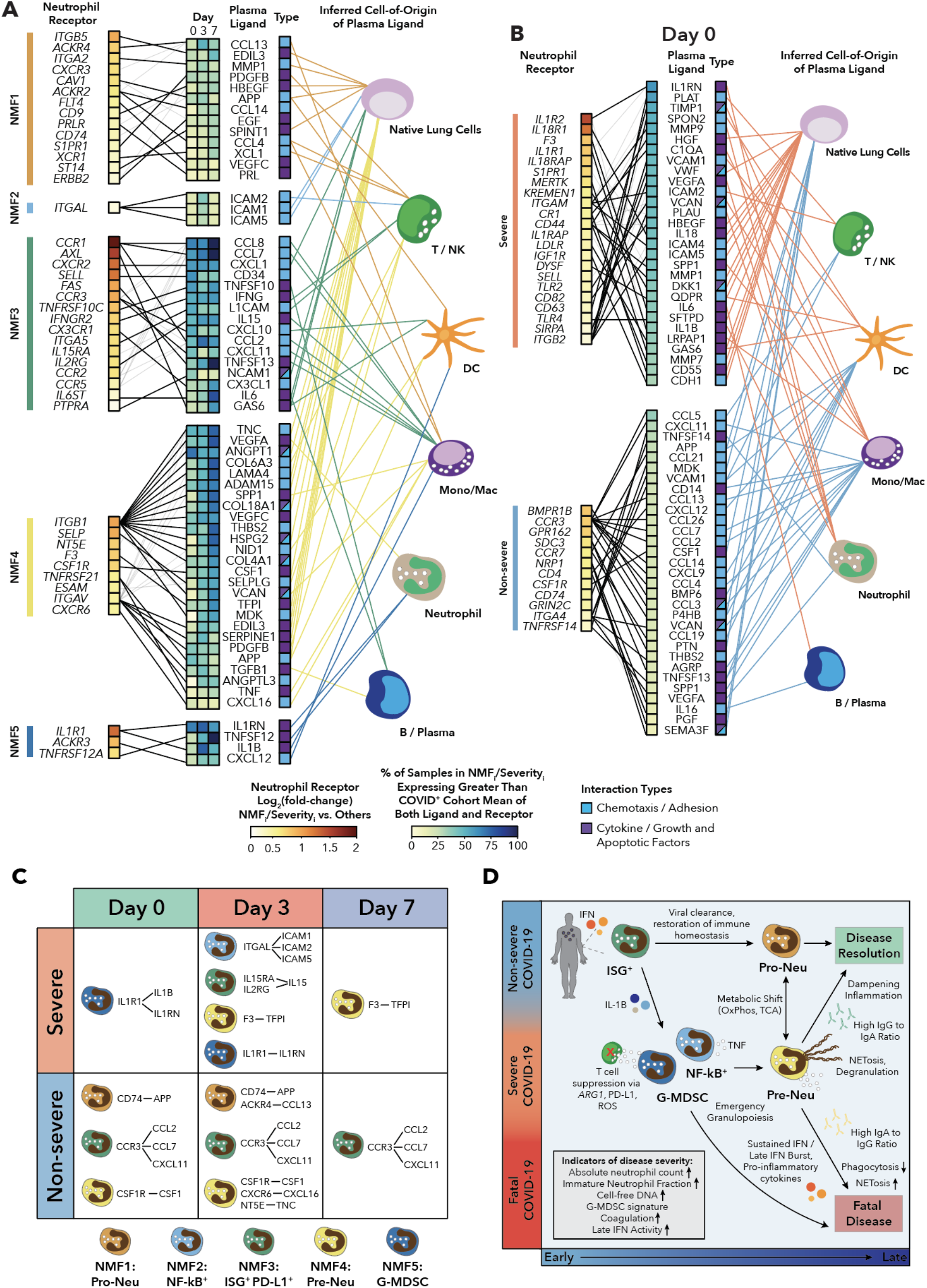
Ligand-Receptor Interactions in Plasma are Potential Drivers of Neutrophil Phenotype and Disease Severity. **(A)** Ligand-receptor analysis for differentially expressed ligands in plasma and receptors on neutrophils between NMF clusters for all COVID-19-positive samples. Ligands and receptors are color-coded by NMF cluster membership. Receptors are color-scaled according to the log_2_(fold-change) between expression in a given NMF cluster and all other clusters. Ligands are color-scaled according to the percentage of samples within the cluster for which the ligand and receptor are both expressed above the overall mean expression. Ligands matching with multiple receptors are color-scaled according to the highest percentage, and the secondary interactions are indicated with reduced line width. Ligands are then mapped to the inferred cell-of-origin based on BAL fluid single cell RNA-seq data from Bost *et al*, 2020. Ligand cell-of-origin is determined by the cluster with the highest average z-scored expression, and multiple cell types are indicated if the mean z-score of the second highest cluster has a z-score difference of less than 0.1. **(B)** Ligand-receptor analysis for differentially expressed ligands in plasma and receptors on neutrophils between COVID-19-positive severe and non-severe samples on Day 0. Ligands and receptors are color-coded by severity. Receptors are color-scaled according to the log_2_(fold-change) between severity groups. Ligands are color-scaled according to the percentage of samples within the severity group for which the ligand and receptor are both expressed above the overall mean expression. **(C)** Table highlighting the overlap between the unique neutrophil NMF subtype ligand-receptor interactions and the interactions associated with severe or non-severe disease, broken by Day. **(D)** Model summarizing neutrophil contributions to COVID-19 pathogenesis.

Among the more severe subtypes, NMF5 (G-MDSC) had the highest expression of the IL1 receptor *IL1R1* and was marked by the highest levels of the matched plasma ligands IL1RN and IL1B. These IL1 family ligands originate from neutrophils, suggesting that the G-MDSC-like phenotype, characteristic of severe COVID-19 disease, may be driven by a positive feedback loop of neutrophil-derived IL1B signaling. NMF4 (Immature Activated) had the highest expression of *ITGB1* and *ITGAV*, which both interact with a large number of potential ligands, the majority of which originate from native lung cells. In particular, the *IL1R1* receptor on neutrophils was associated with both NMF5 and severe disease, and the *F3*-TFPI interaction, which is implicated in coagulation, was associated with both NMF4 and severe disease, consistent with the many other indicators that NMF4 is involved in NETosis (Figure 7C).

In the less severe-specific subtypes, the enriched potential interactions between NMF1 (Pro-Neu) neutrophils and plasma ligands featured many growth factor signaling pathways (PDGFB, HBEGF, EGF, EREG, VEGFC, PRL), and the majority of ligands originated from native lung cells. NMF3 (PD-L1^+^ ISG^+^) showed strong upregulation of receptors involved in migration and activation (*CCR1*, *CXCR2*, *SELL*, *CCR3*) and their ligands (CCL8, CCL7, CD34). As expected, the interaction between *IFNGR2* and IFNG was identified in this cluster, explaining the overall phenotype of the ISG neutrophils. A higher fraction of ligands was mapped back to monocytes/macrophages for NMF3 than any other cluster. *CCR3* interactions were identified in both NMF3 and non-severe disease, and *CD74* interactions were associated with both NMF1 and non-severe disease (Figure 7C).

Similarly, the ligand-receptor interaction analysis for differentially expressed ligands and receptors between severe and non-severe patients revealed several interactions driving severity within the neutrophil compartment, including the neutrophil ligands IL1RN, MMP9, VEGFA, PLAU, and IL1B. Of note, we found at least one potential interaction within the uPA/uPAR system in severe patients across all three days (PLAU-*ITGB2*, PLAU-*ITGAM*, PLAU-*PLAUR*, SERPINE1-*PLAUR*, MMP12-*PLAUR*). PLAU/uPA, which was mapped back to neutrophils in COVID-19 bronchoalveolar lavage (BAL) fluid, has been shown to amplify NF-kB responses in neutrophils including increased production of IL1B, MIP-2, and TNFa, which can result in acute lung injury^70^.

## DISCUSSION

Here, we present a comprehensive characterization of circulating neutrophils from hospitalized COVID-19 patients. We obtained a total of 781 samples across 388 patients, with disease severity stratified from death to healthy controls. Using bulk transcriptomic analysis of enriched neutrophils, plasma proteomics, and high-throughput antibody profiling, we identified diverging neutrophil phenotypes between severe and non-severe disease and potential drivers of these phenotypes, indicating several potential pathways for intervention. We highlighted similarities between COVID-19 neutrophil dysregulation and phenotypes found across multiple disease pathologies, suggesting that the mechanistic insight from this study could be applicable in other disease contexts. Finally, we validated our unbiased findings using functional antibody-dependent assays probing neutrophil phagocytosis, reactive oxygen species production, and NETosis (Figure 7D).

We first utilized unbiased NMF clustering to define six distinct neutrophil states associated with COVID-19 and SARS-CoV-2-negative respiratory disease. Despite the more limited cellular resolution of bulk transcriptomics, we compared the gene signatures of each cluster with single-cell RNA-sequencing data from an independent cohort of COVID-19 patients and found strong overlap between the two^11^. Furthermore, our network analysis across disease contexts demonstrates that there is a common set of transcriptionally-distinct neutrophil states that exists across sepsis^43^, cancer^42^, and acute viral infection, each having their own associations with disease severity or resolution. Therefore, the characteristics associated with each neutrophil state and potential therapeutic interventions targeting specific states may be applicable across disease contexts. While these neutrophil states are detected across multiple studies, future studies will be required to isolate phenotypically distinct cell populations and assess their regulatory or inflammatory properties. Additionally, while previous studies have focused on analysis of the immune response to SARS-CoV-2 at a single time point, our analysis of longitudinal samples allowed us to distinguish signatures associated with outcome at initial hospitalization from signatures that developed over time. We observed that all patients, regardless of disease outcome, exhibit an interferon-driven viral response neutrophil phenotype upon hospitalization, but this signature decreases with time and is replaced either by a suppressive G-MDSC-like signature in severe patients or a neutrophil progenitor signature in non-severe patients. Furthermore, we observed that relative to patients who survived, patients who died maintained higher levels of interferon response on Days 3 and 7 of hospitalization, potentially indicating that interferon signaling may be a longitudinal biomarker of severe disease. Finally, multimodal analysis integrating transcriptomics and proteomics from matched plasma revealed a potential positive feedback loop of neutrophil IL1B signaling in severe patients. Overall, our findings emphasize the importance of a well-powered cohort with both the granularity to define neutrophil states and longitudinal sample collection to capture the dynamics of disease progression or resolution.

Our evolving understanding of the differential impact of IgA and IgG antibody isotypes on neutrophil effector functions has extensive therapeutic implications. The observation that patients who died maintained a higher IgA1-to-IgG1 ratio over patients who were intubated but survived directly implicates the humoral response and neutrophil effector functions in the most extreme outcomes in COVID-19. As it has been reported, SARS-CoV-2 infection begins in the nasal passages and thus triggers a strong mucosal IgA response followed by a wave of IgA antibody-secreting cells entering circulation^71^. Based on the results of our antibody isolation experiments, we hypothesize that this IgA-heavy humoral response may promote systemic circulating neutrophil dysregulation with higher rates of NETosis in otherwise uninfected parts of the body. While IgA-induced NETosis would have a beneficial component in mucosal linings by preventing viral entry, it would be ineffective or even harmful in other locations, as neutrophils in the bloodstream perform their most protective phagocytic functions in response to IgG antibodies. Many studies have shown that NETosis is a defining feature of severe disease^49,72,73^, and here, we demonstrate that NETosis can be strongly induced by IgA antibodies, which may be the dominant effector function occurring in patients with high ratios of IgA1-to-IgG1 antibodies in plasma. Several potential therapeutics have been suggested for use in autoimmune disease contexts aimed at inhibiting NETosis such as PAD4 inhibition or administration of recombinant human thrombomodulin^74^, and similar strategies could be applied to NETosis in the context of severe COVID-19. Additionally, clinical trials targeting IL1B aimed at decreasing NETosis are underway (ClinicalTrials.gov identifier: NCT04594356). We also hypothesize that patients with high abundance of G-MDSCs would benefit the most from these therapeutics, as the NMF subtype ligand-receptor interaction analysis suggests that G-MDSCs may be induced by a positive feedback loop of IL1B signaling in circulating neutrophils.

As a potential therapeutic intervention, we hypothesize that infusion of convalescent plasma enriched for IgG and depleted for IgA antibodies would have a stronger impact on patient recovery than non-enriched plasma. Additionally, given the association between the antibody profiles and survival with patient intubation status but not with disease severity overall, we predict that this would provide the most benefit in the cases of the most severe patients, and would likely not be as effective in preventing progression to severe disease. Recently, an analysis of a convalescent plasma infusion trial which was not enriched for IgG showed that plasma infusion conferred a survival benefit to hospitalized COVID-19 patients who did not receive mechanical ventilation^75^. It may be possible that survival among the mechanically ventilated group would increase once the additional IgA antibodies were depleted and the neutrophil response was shifted towards phagocytosis and away from additional NETosis. Additionally, our observations have implications for vaccine development. There is a possibility that intranasal vaccines which promote a higher IgA response could be even more effective at preventing viral entry, whereas IgG responses would initiate once IgG antibodies in blood come into contact with the virus.

While manipulation of the antibody landscape seems to hold promise for effective interventions, the drivers of the humoral response and class-switch recombination in COVID-19 are still poorly understood. In this study, we identify a strong association between higher IgA1-to-IgG1 ratios in plasma and circulating IFNL1, though no study to date has drawn a connection between type III interferons and isotype switching to IgA. Future studies should aim to determine which plasma cells are responsible for the serum IgA secretion in response to SARS-CoV-2 infection, whether it is from secondary lymphoid organs, or expanding B lineage cells in mucosal tissues, or distant plasmablasts stimulated by circulating factors unique to COVID-19. A recent study suggested that TNFα-secreting cells could be responsible for the loss of germinal centers in the secondary lymphoid organs of severe COVID-19 patients^76^. Though neutrophils produce lower levels of TNFα than highly-inflammatory macrophage populations, the consistent robust enrichment of the TNFα signaling via NFkB pathway in neutrophils suggest that neutrophils may also play a role in the loss of germinal centers and weakening of humoral responses in severe COVID-19 patients. Many recent studies in patients treated with TNFα blockers for autoimmune diseases such as rheumatoid arthritis, psoriasis, and inflammatory bowel disease demonstrated clinical benefit from these therapeutics, but results from full-scale clinical trials are still needed^77–79^.

We acknowledge several limitations of our study. First, we performed bulk transcriptomics rather than single-cell RNA-sequencing, so the neutrophil state gene signatures reflect a mixture of true neutrophil subtypes. Second, our samples were enriched for neutrophils via magnetic bead negative selection, and high purity of all samples across the cohort could not be guaranteed; thus, we used estimated cell type proportions as covariates in all analyses, but the expression of contaminating cell type-specific genes and cytokines cannot be excluded. Third, our time course data was collected on Days 0, 3, and 7 of hospitalization, though patients may have been infected for varying amounts of time prior to enrollment, so this cannot be considered an exact time course. Fourth, we only collected longitudinal samples from hospitalized patients, so we were unable to study patients pre-hospitalization or patients who were infected but never hospitalized. Fifth, sample collection at later time points was biased towards sicker patients as they needed to stay in the hospital for a longer period of time. Sixth, we did not collect a validation cohort, so our findings will need to be validated in external cohorts with similar multimodal data structures. Seventh, our study relies on blood draws and thus only provides insights into circulating factors that play a role in COVID-19 disease pathology, yet the findings regarding type III interferons and the IgA-dominant humoral response strongly implicate mucosal immunity; future studies should focus on longitudinal SARS-CoV-2 immunity occurring along mucosal barriers. Finally, samples were collected at the onset of the pandemic before there were recommended treatment options, and thus it is not known how different drug treatments such as dexamethasone, tocilizumab, or other monoclonal antibody therapeutics would affect neutrophil phenotypes; furthermore, with the advent of COVID-19 vaccines, it is not known how the adenovirus or mRNA vaccines modulate neutrophil function by the antibody profiles that they produce.

In summary, our study elucidates how circulating neutrophils and their interactions with soluble factors drive COVID-19 disease severity, providing insight into this crucial and abundant cell type. We propose a model of SARS-CoV-2 infection in which antibody profiles drive neutrophils either to aid in disease resolution through phagocytosis or contribute to tissue damage via NETosis. Further, we hypothesize that therapies which simultaneously aim to ablate suppressive G-MDSC-like neutrophils and prevent excessive NETosis in circulation have the potential to aid with disease resolution in severe patients.

## STAR METHODS

Detailed methods are provided in the online version of this paper and include the following:

- KEY RESOURCES TABLE
- RESOURCE AVAILABILITY

- Lead contact
- Materials availability
- Data and code availability
- EXPERIMENTAL MODEL AND SUBJECT DETAILS

- MGH patients cohort description
- METHOD DETAILS

- Neutrophil isolation and lysis
- Patient matched plasma isolation
- Cell-free DNA (cfDNA) quantification
- Smart-Seq2 cDNA preparation
- Library construction and sequencing
- IgG subclass, isotype, and FcγR binding
- Antibody-dependent neutrophil phagocytosis (ADNP) assay
- SARS-CoV-2 spike specific IgG and IgA isolation
- Antibody-dependent neutrophil activation and ROS release
- NETosis Assay
- MPO ELISA
- QUANTIFICATION AND STATISTICAL ANALYSIS

- RNA-seq alignment
- Quality control
- Neutrophil fraction estimation and contamination control
- Dimensionality reduction and visualization
- Differential expression analysis
- Gene set enrichment analysis
- NMF clustering analysis
- Sample pathway scoring
- Clustering Analysis for Single-cell Blood Neutrophils from Sepsis Patients
- Neutrophil state network analysis
- Schulte-Schrepping single-cell RNA-seq reanalysis for early-late threshold
- ARDS log fold-change comparisons
- Day:Severity interaction analysis
- Logistic regression models to predict severe COVID-19 on Day 0
- Plasma proteomic markers of neutrophil subtypes
- Comparison of differential expression and plasma proteomic data
- Ligand-receptor interaction analysis

### KEY RESOURCES TABLE

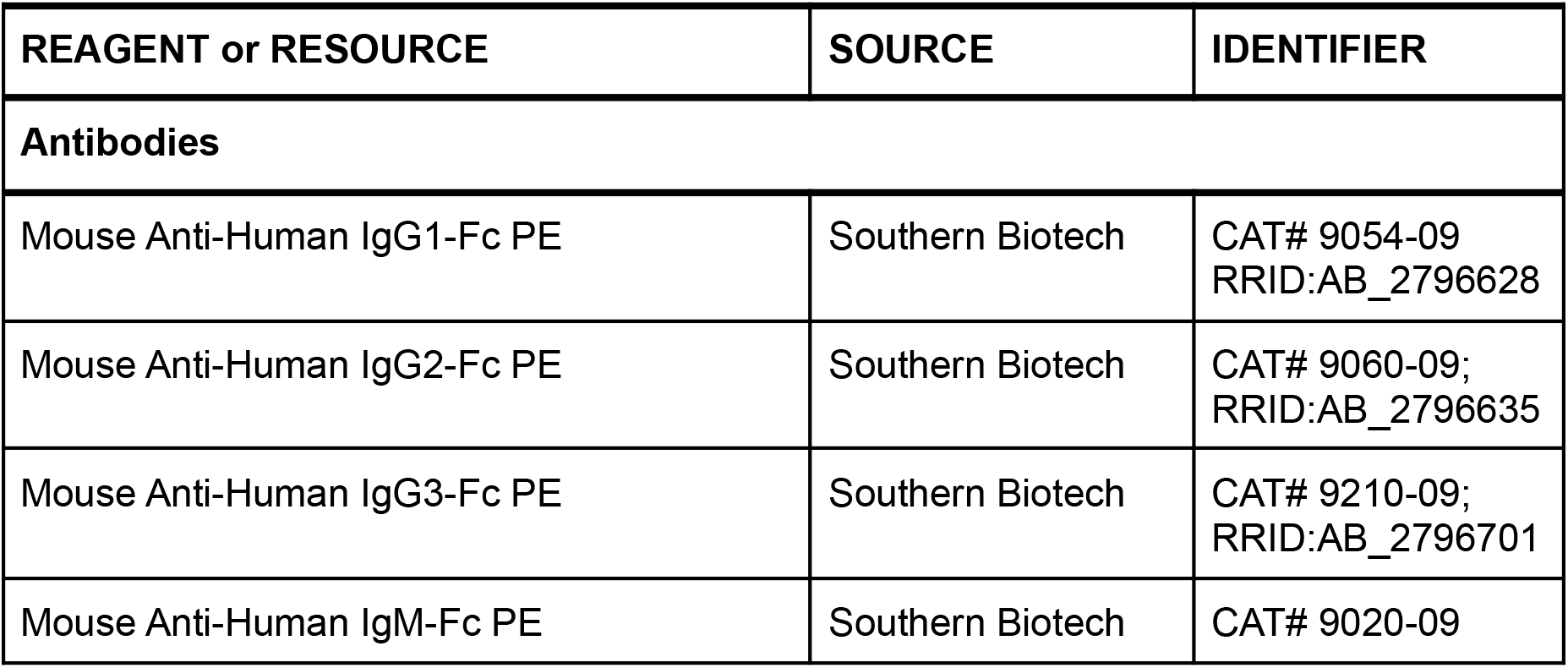
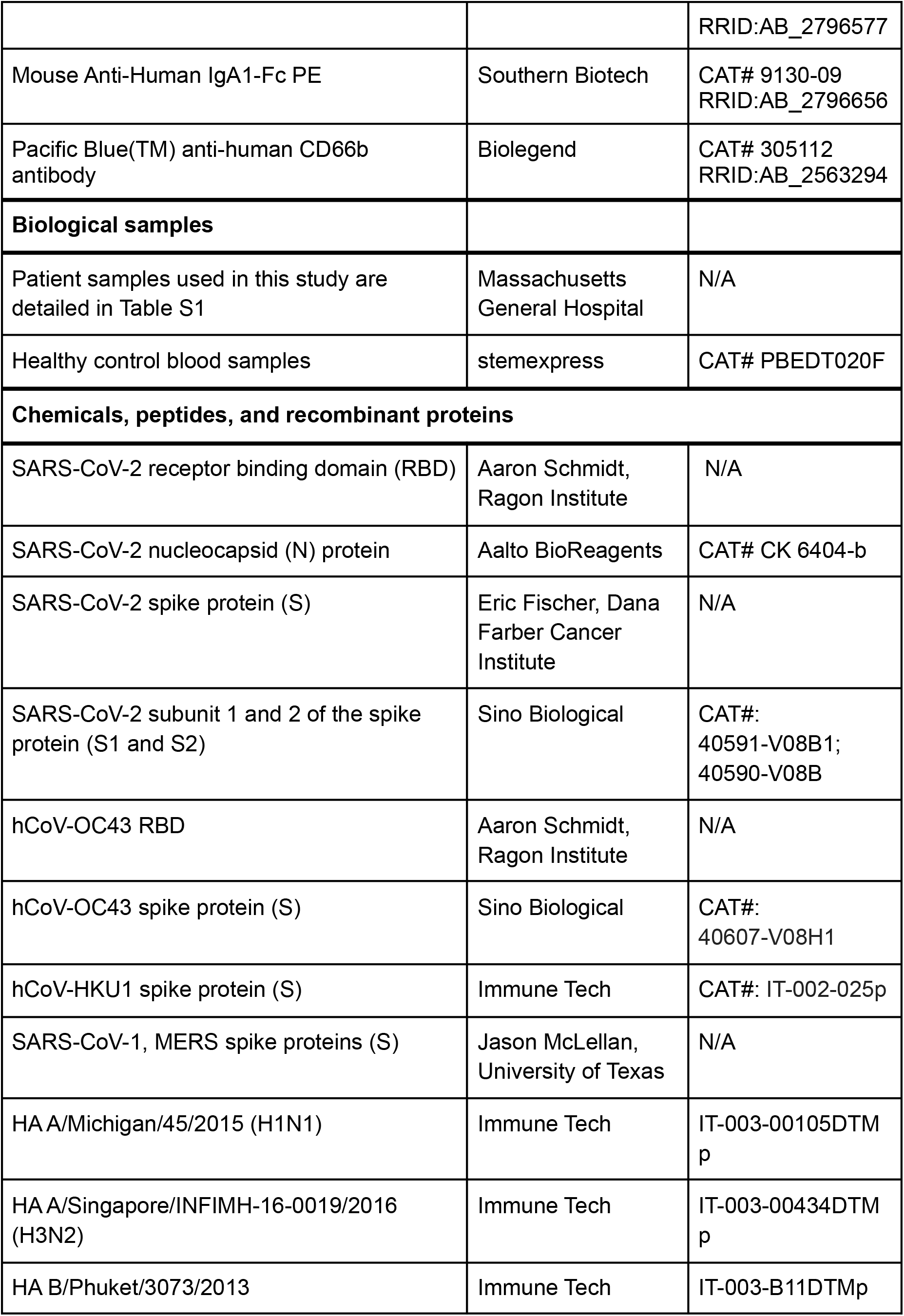
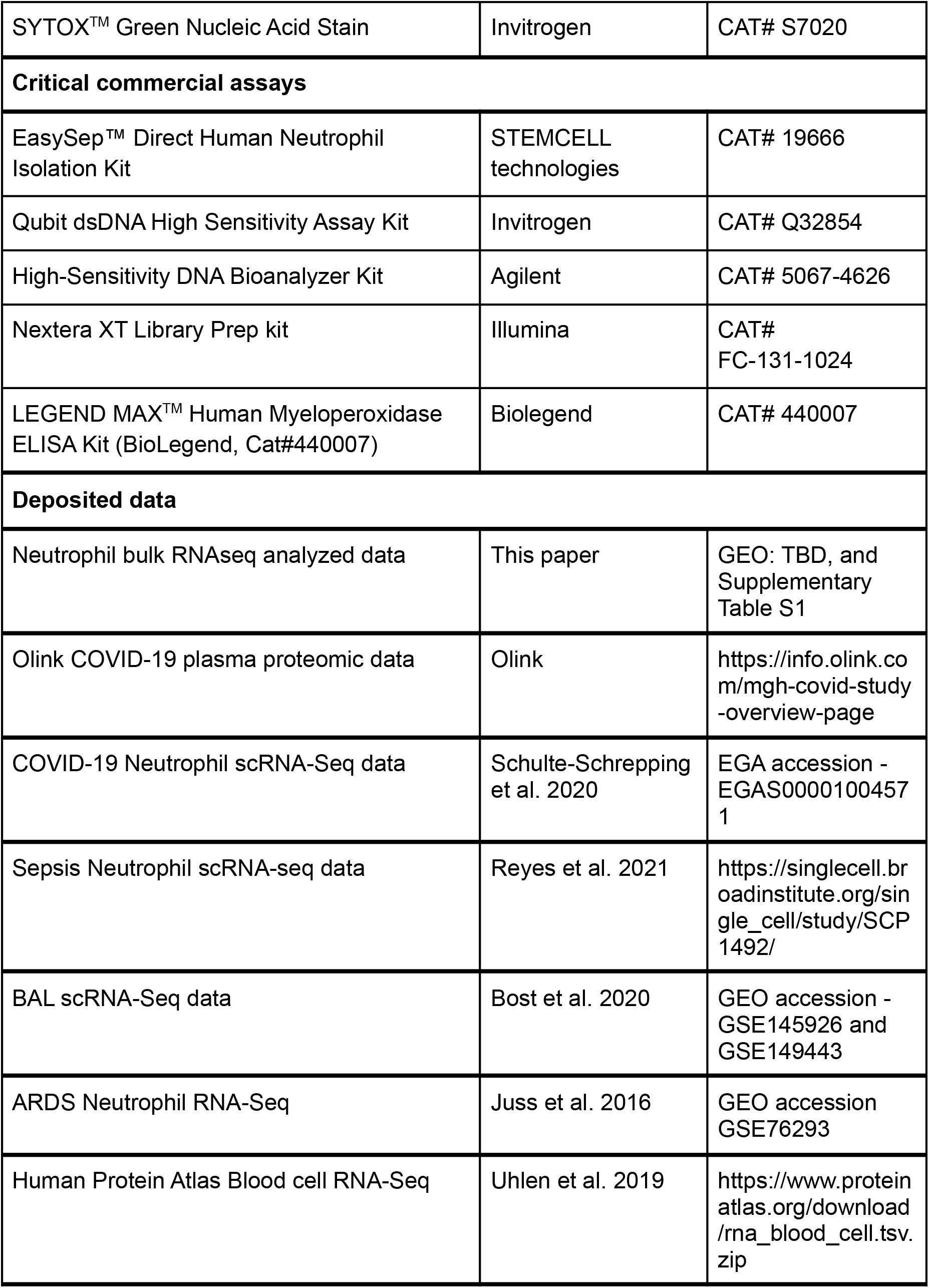
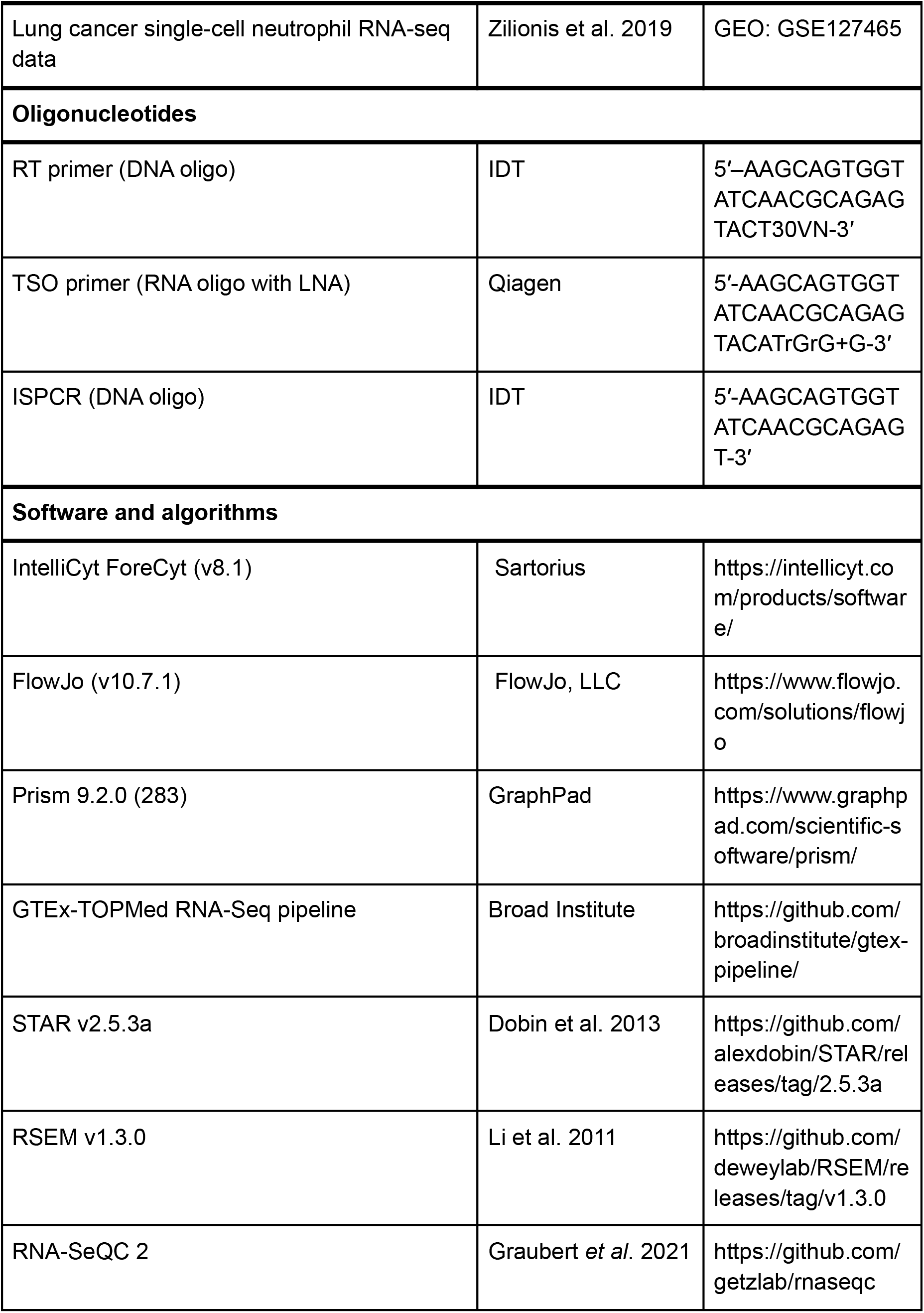
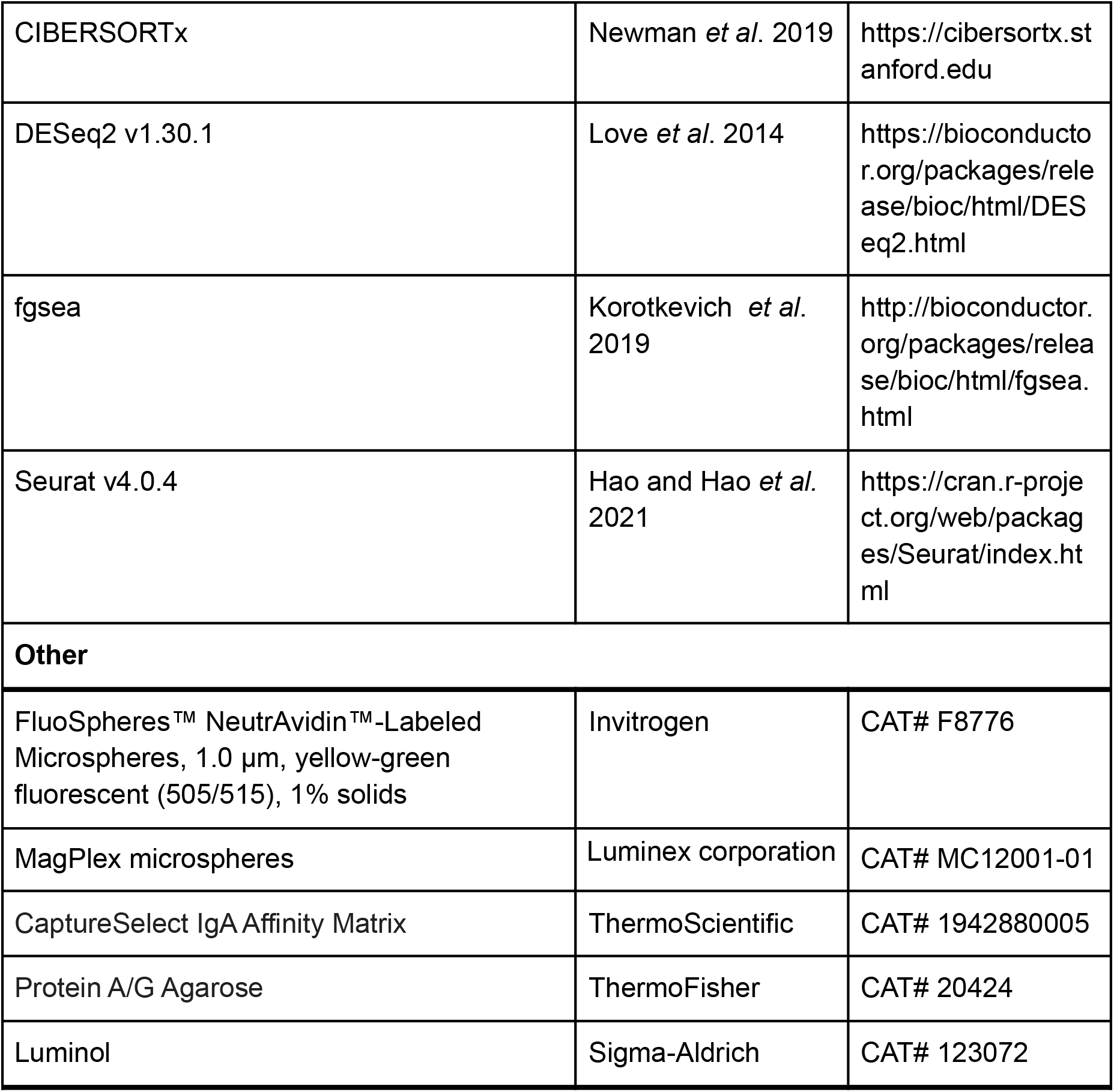

### RESOURCE AVAILABILITY

#### Lead contact

Further information and requests for resources and reagents should be directed to and will be fulfilled by the lead contact, Moshe Sade-Feldman.

#### Materials availability

This study did not generate new unique reagents.

#### Data and code availability

Due to IRB consent limitations, raw sequencing data is not publicly available. However, the read count matrix and TPM matrix used in this study will be available in GEO and Supplementary Table S1. All code used for the analysis is deposited in GitHub at https://github.com/lasalletj/COVID_Neutrophils and all additional files required to run the code will be deposited in Zenodo upon acceptance.

### EXPERIMENTAL MODEL AND SUBJECT DETAILS

#### MGH patients cohort description

Between March to May 2020 during the peak of the COVID-19 pandemic, we enrolled a total of 380 patients 18 years or older who presented in Massachusetts General Hospital Emergency Department (ED) with acute respiratory distress and clinical concern for COVID-19. The study was approved by the Mass General Brigham Institutional Review Board under protocol 2017P001681, with an approval for a waiver of informed consent in compliance with the 45CFR 46, 2018 Common rule. Out of the 380 patients enrolled in this study, 304 tested positive for SARS-CoV-2 (COVID-19^+^), while 76 patients that were admitted to the ED with similar symptoms were tested negative (COVID-19^−^) and were used as controls in this study. Additionally we also collected blood samples from 8 healthy donors. For each patient, medical history and clinical data were collected and are presented in Table S1 and as previously described^26^. Samples were collected at three different time points: Day 0 upon admission to the ED (n=374 samples); Day 3 (n=212 samples) and Day 7 (n=143 samples) for COVID-19^+^ hospitalized patients. In addition, in some cases up to day 28 post-admission to the ED, a fourth blood sample was collected upon a major change in the clinical status, and was termed as an event driven sample (n=44 samples). Acuity categories were classified into five classes (A1-A5) using the WHO ordinal outcomes scale as recently described in Filbin *et al.* 2021, with the following classifications: A1 and A2 were classified as severe disease, with A1 defined as death within 28 days (n=40 patients and 96 samples), and A2 for patients that survived within 28 days but required mechanical ventilation and/or intubation (n=67 patients and 222 samples). Groups A3-A5 were defined as non-severe, with A3 classified as patients that required supplemental oxygen (n=133 patients and 298 samples), A4 hospitalized but no need for supplemental oxygen (n=41 patients and 45 samples), and A5 classified as patients that were discharged from ED in the first 24 hours and did not return to the hospital within 28 days (n=23 patients and 23 samples; Figure S1). Primary outcomes for each patient (Acuity_Max_ or Severity_Max_) were defined as the most severe disease level with 28 days of enrollment. Since this study included the enrollment of patients with an approval for a waiver of informed consent, demographic information, and other clinical parameters described in this study (e.g., blood counts, LDH, CRP etc.) are limited and reported in quintiles.

### METHOD DETAILS

#### Neutrophil isolation and lysis

Blood samples were collected in EDTA vacutainer tubes and transported to the laboratory. Neutrophils were isolated from whole blood via negative selection using the EasySep™ Direct Human Neutrophil Isolation Kit (STEMCELL Technologies, Cat# 19666). All described procedures in this section were done at room temperature. Between 0.25-0.5 mL whole blood was lysed with ACK Lysis Buffer (ThermoFisher Scientific, Cat# A1049201) in a 15 mL conical tube and white blood cells were pelleted at 300 x*g* for 5 min. Following aspiration of the lysed red blood cells and resuspension of the pellet in 250 µL 1 mM EDTA in PBS, 50 µL each of the RapidSpheres and Isolation Cocktail were added to the cell suspension. Following a 5 min incubation, sample volumes were completed to 4 mL with 1 mM EDTA in PBS, mixed gently, and placed on an EasyEights™ EasySep™ Magnet (STEMCELL Technologies, Cat# 18103) for 5 min. Next, supernatants were transferred to new 15 mL conical tubes, 25 µL RapidSpheres were added, and the samples were gently mixed and incubated for 5 min. Samples were then placed on the magnet, and after 5 min incubation supernatants were transferred to new tubes, and were placed immediately on the magnet for a second incubation before the supernatants containing the enriched neutrophil populations were collected, pelleted, and resuspended in 1 mM EDTA in PBS. Cells were counted on a TC20™ Automated Cell Counter (Bio-Rad Laboratories, Inc., Cat# 1450102) with trypan blue staining for dead cell exclusion. Neutrophils were then lysed in TCL Buffer (QIAGEN, Cat# 1031576) with 1% 2-Mercaptoethanol at a concentration of 1000 cells/µL, flash-frozen on dry ice, and then stored at −80 °C until use.

#### Patient matched plasma isolation

Following the aliquoting of 0.25-0.5 mL whole blood for neutrophil isolation, remaining blood volumes were diluted 1:3 with room temperature RPMI. Each diluted sample was then added carefully to a SepMate tube (STEMCELL Technologies, Cat# 85450 or 85415) that had been prefilled with 15 mL Ficoll (VWR, Cat#21008-918). Samples were spun at 1200 x*g* for 20 min at 20 °C with maximum acceleration and the brake on. After centrifugation, the plasma layer was transferred into a clean conical tube and spun at 1000 x*g* for 5 mins at 4 °C to pellet any remaining cell debris. Without disturbing the pellet, each sample was aliquoted into 1.5 mL Cryovials (VWR, Cat# 66008-710) and frozen at −80 °C until analysis.

#### Cell-free DNA (cfDNA) quantification

cfDNA was quantified using the Qubit dsDNA High Sensitivity Assay Kit (Invitrogen, Cat# Q32854). 98 µL of DNA dye was aliquoted into each well of a 96-well black clear bottom plate (Corning, Cat# 3904). Plasma samples which had been pre-aliquoted into 96-well Eppendorf PCR plates were thawed at RT, vortexed, and spun down briefly. 2 uL of plasma sample was added to each well of the assay plate. Fluorescence was quantified on a Cytation 5 Microplate reader at 523 nm.

#### Smart-Seq2 cDNA preparation

cDNA was prepared from bulk populations of 2×10^4^ neutrophils per sample via the Smart-Seq2 protocol^80^ with some modifications to the reverse transcription step as previously described^81^. 20 µL (at a concentration of 1000 cells/µL) of neutrophil lysates were thawed on ice and plated into 96-well plates prior to centrifugation at 1500 rpm for 30 s. RNA was purified with Agencourt RNAClean XP SPRI beads (Beckman Coulter, Cat# A63987) and then the samples were resuspended in 4 µL of Mix-1 [Per 1 sample: 1 µl (10 µM) RT primer (DNA oligo) 5′–AAGCAGTGGTATCAACGCAGAGTACT30VN-3′; 1 µl (10 µM) dNTPs; 1µl (10%, 4 U/µl) recombinant RNase inhibitor; 1 µl nuclease-free water], denatured at 72 °C for 3 min and placed immediately on ice for 1 min before 7 µL of Mix-2 [Per 1 sample: 0.75 µl nuclease-free water; 2 µl 5X RT buffer (Thermo Fisher Scientific, Cat# EP0753); 2 µl (5 M) betaine; 0.9 µl (100 mM) MgCl2; 1 µl (10 µM) TSO primer (RNA oligo with LNA) 5′-AAGCAGTGGTATCAACGCAGAGTACATrGrG+G-3′; 0.25 µl (40 U/µl) recombinant RNase inhibitor; 0.1 µl (200 U/µl) Maxima H Minus Reverse Transcriptase] was added. Reverse transcription reactions were performed at 50 °C for 90 min, followed by 5 min incubation at 85 °C. Then, 14 µL of Mix-3 [Per 1 sample: 1 µl nuclease-free water; 0.5 µl (10 µM) ISPCR primer (DNA oligo) 5′-AAGCAGTGGTATCAACGCAGAGT-3′; 12.5 µl 2X KAPA HiFi HotStart ReadyMix] was added to each well and the whole-transcriptome amplification step was performed at 98 °C for 3 min, followed by 16 cycles of [98 °C for 15 s, 67 °C for 20 s, and 72°C for 6 min], and final extension at 72C for 5 min. cDNA was purified using AgencourtAMPureXP SPRI beads (Beckman Coulter, Cat# A63881) as described^81^, to remove all primer residue. Quality control was performed on samples prior library construction and included: (1) concentration measurements via the Qubit dsDNA high sensitivity assay kit (Invitrogen, Cat# Q32854) on the Cytation 5 Microplate Reader (BioTek); (2) cDNA size distribution using the High-Sensitivity DNA Bioanalyzer Kit (Agilent, Cat# 5067-4626).

#### Library construction and sequencing

Libraries were generated using the Nextera XT Library Prep kit (Illumina, Cat# FC-131-1024) with custom indexing adapters^81^ in a 384-well PCR plate, followed by a cleanup step to remove residual primer dimers. Pooled libraries containing 384 samples were then sequenced on a NovaSeq S4 (Illumina) using paired-end 150-base reads. Additionally, 16 samples were sequenced on a NextSeq 500 sequencer (Illumina), using paired-end 38-base reads. This approach insured an appropriate coverage for all samples analyzed in this study.

#### IgG subclass, isotype, and FcγR binding

SARS-CoV-2 and eCoV-specific antibody subclass/isotype levels were assessed using a 384-well based customized multiplexed Luminex assay, as previously described^27^. SARS-CoV-2 receptor binding domain (RBD) (kindly provided by Aaron Schmidt, Ragon Institute), SARS-CoV-2 nucleocapsid (N) protein (Aalto BioReagents), and SARS-CoV-2 spike protein (S) (kindly provided by Eric Fischer, Dana Farber), SARS-CoV-2 subunit 1 and 2 of the spike protein (S1 and S2) (Sino Biological), as well as human eCoV antigens: hCoV-OC43 RBD (kindly provided by Aaron Schmidt, Ragon Institute), hCoV-OC43 spike protein (S) (Sino Biological), hCoV-HKU1 spike protein (S) (Immune Tech), SARS-CoV-1, MERS spike proteins (S) (kindly provided by Jason McLellan, University of Texas) were used to profile specific humoral immune response. A mix of HA A/Michigan/45/2015 (H1N1), HA A/Singapore/INFIMH-16-0019/2016 (H3N2), HA B/Phuket/3073/2013 (Immune Tech) was used as a control. Antigens were coupled to magnetic Luminex beads (Luminex Corp) by carbodiimide-NHS ester-coupling (Thermo Fisher). Antigen-coupled microspheres were washed and incubated with plasma samples at an appropriate sample dilution (1:500 for IgG1 and 1:100 for all other readouts) for 2 hours at 37°C in 384-well plates (Greiner Bio-One). Unbound antibodies were washed away, and antigen-bound antibodies were detected by using a PE-coupled detection antibody for each subclass and isotype (IgG1, IgG2, IgG3, IgG4, IgA1, and IgM; Southern Biotech). After 1h incubation, plates were washed, and flow cytometry was performed with an IQue (Intellicyt), and analysis was performed on IntelliCyt ForeCyt (v8.1). PE median fluorescence intensity (MFI) is reported as a readout for antigen-specific antibody titers.

#### Antibody-dependent neutrophil phagocytosis (ADNP) assay

ADNP was conducted as previously described^58^. SARS-CoV-2 Spike proteins were biotinylated using EDC (Thermo Fisher) and Sulfo-NHS-LC-LC biotin (Thermo Fisher) and coupled to Cat. To form immune complexes, antigen-coupled beads were incubated for 2 hours at 37°C with serum and then washed to remove unbound antibodies. The immune complexes were incubated for 1 hour with RBC-lysed whole blood. Following the incubation, neutrophils were stained for CD66b+ (Biolegend, Cat# 305112) and fixed in 4% PFA.

Flow cytometry was performed to identify the percentage of cells that had phagocytosed beads as well as the number of beads that had been phagocytosed (phagocytosis score = % positive cells × Median Fluorescent Intensity of positive cells/10000). Flow cytometry was performed with an IQue (Intellicyt) or LSRII(BD), and analysis was performed using IntelliCyt ForeCyt (v8.1) or FlowJo V10.7.1.

#### SARS-CoV-2 spike specific IgG and IgA isolation

IgA were purified from human plasma samples using CaptureSelect IgA Affinity Matrix (Thermo Fisher Scientific, Cat#1942880005), and flowthrough was used to purify the IgG with Protein A/G Agarose (Thermo Fisher Scientific, Cat#:20424). For both, the capture matrices were washed three times with Binding Buffer (0.1 M phosphate, 0.15 M sodium chloride; pH 7.2) and incubated overnight with 1:5 diluted plasma samples. Antibodies bound to matrices were washed 3x with PBST by centrifugation and eluted with Elution Buffer (0.1 M glycine, pH 2-3). The antibodies were collected to tubes containing Neutralization Buffer (1 M Tris, pH 8-9) and used for further analysis. The presence of IgA and IgG was confirmed by ELISA.

#### Antibody-dependent neutrophil ROS release

A high-binding 96-well plate was coated with SARS-CoV-2 Spike protein (5ug/ml) and blocked with 5% BSA. Isolated antibodies were added and incubated for 2h at RT; afterward, the plate was washed three times with PBST. Neutrophils were isolated from fresh blood using the EasySep™ Direct Human Neutrophil Isolation Kit (STEMCELL Technologies, Cat# 19666) and adjusted to the concentration of 10^6^ cells/mL. Luminol (Sigma-Aldrich, Cat#123072) was diluted in DMSO and added to neutrophils at the final concentration of 0.2 mg/mL. Cells with luminol were added to each well, and chemiluminescence was read immediately on a plate reader (for around two hours). ROS release was quantified as chemiluminescence count/second.

#### NETosis Assay

Methods were adapted from a previous publication^47^. All reagents used in this section were allowed to equilibrate to RT before use.

*Poly-L-lysine plate coating*: 96-well black clear bottom plates (Corning, Cat# 3904) were coated in 40µL of a 1:10 dilution of 0.01% poly-L-lysine (Sigma-Aldrich, Cat#P4707-50ML) in sterile water. Plates were incubated at 37°C for one hour and subsequently washed twice with sterile water, and were allowed to dry for at least two hours before use.

*Enhanced neutrophil isolation*: Fresh blood was collected from healthy donors, moved to a 50mL conical, and diluted 1:3 with room temperature RPMI. Diluted samples were added to a SepMate tubes (Stemcell Technologies, Cat# 85450) that had been prefilled with 16 mL Ficoll (VWR, Cat#21008-918). Samples were spun at 1200 x*g* for 20 min at 20 °C with maximum acceleration and the brake on. Plasma and PBMCs were removed, and the high density layer containing erythrocytes and granulocytes was moved to a 50mL tube. Samples then underwent two rounds of red blood cell lysis using ACK Lysis Buffer (ThermoFisher Scientific, Cat# A1049201) and centrifugation for 5 minutes at 1500g, RT. Pellets were resuspended in 500µL of 1 mM EDTA in PBS per 10mL of blood, and 250uL aliquots were moved to 15mL conicals. Negative selection for neutrophils was then performed with the EasySep™ Direct Human Neutrophil Isolation Kit (STEMCELL Technologies, Cat# 19666) with custom modifications. 75 µL each of the RapidSpheres and Isolation Cocktail were added to the cell suspension. Following a 5 min incubation, sample volumes were completed to 4 mL with 1 mM EDTA in PBS, mixed gently, and placed on an EasyEights™ EasySep™ Magnet (STEMCELL Technologies, Cat# 18103) for 5 min. Next, supernatants were transferred to new 15 mL conical tubes, 37.5 µL RapidSpheres were added, and the samples were gently mixed and incubated for 5 min. Samples were then placed on the magnet, and after 5 min incubation supernatants were transferred to new tubes, and were placed immediately on the magnet for a second incubation before the supernatants containing the enriched neutrophil populations were collected, pelleted, and resuspended in PBS. Cells were counted on a TC20™ Automated Cell Counter (Bio-Rad Laboratories, Inc., Cat# 1450102) with trypan blue staining for dead cell exclusion.

*NETosis induction and quantification*: Using the highly-enriched neutrophil samples, 50,000 cells were plated in each well of the poly-L-lysine-coated 96-well black clear bottom plates. Plates were then incubated for 20 minutes at 37°C and 5% CO_2_ to allow neutrophils to adhere. Supernatant was then gently removed and immediately replaced with 32µL RPMI + L-glu with 625nM SYTOX^TM^ Green Nucleic Acid Stain (Invitrogen, Cat#S7020). 8µL of patient-isolated antibody was then added to each well for a total of 40µL per well with a final 1:5 dilution of free antibody and 500nM SYTOX Green. Plates were then incubated for 4 hours at 37°C and 5% CO_2_. Cells were gently removed from the incubator and fluorescence was quantified on a Cytation 5 Microplate reader at 485nm and 523nm using the area scan setting from the bottom of the plate. Absorbance at 485nm was subtracted from the absorbance at 523nm to obtain corrected RFU values.

#### MPO ELISA

Supernatants from the NETosis assay were collected gently without disturbing cell adhesion and were stored at −80°C until the assay was performed. Samples were thawed at RT and spun down at 1500rpm for 5 minutes, and supernatants were transferred to a new plate, leaving a small amount of liquid at the bottom of each well. The ELISA for quantification of MPO was performed using the LEGEND MAX^TM^ Human Myeloperoxidase ELISA Kit (BioLegend, Cat#440007) according to manufacturer specifications. The standard curve was fitted with a 4-parameter logistic curve-fitting algorithm using the dr4pl package in R.

### Quantification and Statistical Analysis

#### RNA-seq alignment

A custom FASTA was generated from the Homo sapiens (human) genome assembly GRCh38 (hg38) following exclusion of ALT, HLA, and Decoy contigs according to documentation in the Broad Institute GTEx-TOPMed RNA-seq pipeline (https://github.com/broadinstitute/gtex-pipeline/), with an appended SARS-CoV2 genome. GENCODE v35 with the appended SARS-CoV2 GTF was used for annotation. Raw FASTQ files were aligned to the custom genome FASTA in the Terra platform with the Broad Institute GTEx pipeline using STAR v2.5.3a, and expression quantification based on a collapsed annotation was performed using RSEM v1.3.0.

#### Quality control

RNA-SeQC 2^30^ (https://github.com/getzlab/rnaseqc) was used to calculate quality control metrics for each sample. Samples were excluded if they did not meet the following criteria: 1) percentage of mitochondrial reads less than 20%, 2) greater than 10,000 genes detected with at least 5 unambiguous reads, 3) median exon CV less than 1, 4) exon CV MAD less than 0.75, 5) exonic rate greater than 25%, 6) median 3’ bias less than 90%. This filtration kept 698 out of 781 samples (89.4%) (Figure S1). Genes were included in the analysis if they were expressed at a level of 0.1 TPM in at least 20% of samples and if there were at least 6 counts in 20% of samples. In total, 20283 genes passed the filtration criteria.

#### Neutrophil fraction estimation and contamination control

CIBERSORTx^38^ was used to estimate the proportions of mature neutrophils, immature neutrophils, T/NK cells, B cells, plasmablasts, and monocytes in each sample. To generate the signature matrix for deconvolution, we utilized the single-cell RNA-seq data of PBMCs and neutrophils from whole blood from Cohort 2 of the Schulte-Schrepping et al dataset^11^. Using the designations provided in the public data, we created pseudobulks for each cell type per patient by summing the counts of a given cell type, and we excluded pseudobulked cell type samples from individual patients if the cell type had less than 5000 counts. To generate the CIBERSORTx signature matrix, we set limits of 50 to 100 marker genes per cell type, and filtered for only hematopoietic genes. Following the generation of a signature matrix, we ran CIBERSORTx with default parameters to estimate the proportions of each cell type.

Given the levels of immunoglobulin genes from contaminating plasma cells, we created an immunoglobulin score for each sample to use as a covariate for regression (Figure S6). To select genes, we chose the top 115 immunoglobulin genes which were differentially expressed in Day 0 COVID+ vs. COVID- patients (DESeq2, no covariates) and assigned each sample a score according to a previously described method^82^. Briefly, the score was defined as the average log_2_(TPM+1) expression of the immunoglobulin gene set, minus the average log_2_(TPM+1) expression of a control gene set. The control gene set was selected by sorting the entire list of genes by aggregate counts across all samples, breaking the list into 25 bins, and for each gene in the immunoglobulin gene set, selecting 100 genes at random from the same expression bin. Using this method, the control gene set has a comparable distribution of expression levels relative to the immunoglobulin gene set and accounts for the varying complexity between samples.

#### Dimensionality reduction and visualization

PCA and UMAP were performed in R using prcomp() and umap() with default parameters.

#### Differential expression analysis

Differential expression analyses were performed using the DESeq2 package in R^83^. For each analysis *i*, we excluded genes with less than 5 counts in *x_i_* samples. To determine *x_i_*, we generated a curve plotting the required number of samples having ≥5 counts as the independent variable and the number of genes satisfying this condition as the dependent variable. We then selected the inflection point of this curve to be *x_i_*.

#### Gene set enrichment analysis

We performed gene set enrichment analysis using the fgsea package in R using the following pathway sets from MSigDB Release v7.2: H, C5 GO BP. We also performed a search of MSigDB using the keyword “neutrophil” and added the following pathways:

- BIOCARTA_NEUTROPHIL_PATHWAY,
- BIOCARTA_NEUTROPHIL_PATHWAY,
- GO_AZUROPHIL_GRANULE,
- GO_AZUROPHIL_GRANULE_LUMEN,
- GO_AZUROPHIL_GRANULE_MEMBRANE,
- GO_FICOLIN_1_RICH_GRANULE,
- GO_NEGATIVE_REGULATION_OF_NEUTROPHIL_ACTIVATION,
- GO_NEGATIVE_REGULATION_OF_NEUTROPHIL_MIGRATION,
- GO_NEUTROPHIL_CHEMOTAXIS,
- GO_NEUTROPHIL_EXTRAVASATION,
- GO_NEUTROPHIL_MIGRATION,
- GO_POSITIVE_REGULATION_OF_NEUTROPHIL_MIGRATION,
- GO_REGULATION_OF_NEUTROPHIL_ACTIVATION,
- GO_REGULATION_OF_NEUTROPHIL_CHEMOTAXIS,
- GO_REGULATION_OF_NEUTROPHIL_DEGRANULATION,
- GO_REGULATION_OF_NEUTROPHIL_EXTRAVASATION,
- GO_REGULATION_OF_NEUTROPHIL_MEDIATED_CYTOTOXICITY,
- GO_REGULATION_OF_NEUTROPHIL_MIGRATION,
- GO_SPECIFIC_GRANULE,
- GO_SPECIFIC_GRANULE_LUMEN,
- GO_SPECIFIC_GRNAULE_MEMBRANE,
- GO_TERTIARY_GRANULE,
- HP_ABNORMAL_NEUTROPHIL_COUNT,
- HP_ABNORMALITY_OF_NEUTROPHIL_MORPHOLOGY,
- HP_ABNORMALITY_OF_NEUTROPHIL_PHYSIOLOGY,
- HP_ABNORMALITY_OF_NEUTROPHILS,
- HP_IMPAIRED_NEUTROPHIL_BACTERICIDAL_ACTIVITY,
- MARTINELLI_IMMATURE_NEUTROPHIL_DN, MARTINELLI_IMMATURE_NEUTROPHIL_UP,
- NICK_RESPONSE_TO_PROC_TREATMENT_DN,
- NICK_REPSONSE_TO_PROC_TREATMENT_UP,
- REACTOME_NEUTROPHIL_DEGRANULATION.

In addition to these pathways, we added gene sets corresponding to various neutrophil states and signatures: genes up- or down-regulated more than threefold in blood neutrophils from ARDS patients^44^, single-cell neutrophil clusters in blood or lung tissue of patients with lung cancer^42^, single-cell neutrophil clusters from blood of patients with sepsis^43^, and single-cell neutrophil clusters from COVID-19 patients and healthy controls^11^. For single-cell cluster markers, if there were more than 100 marker genes per cluster, gene sets were selected as the top 100 genes ranked by p-value for enrichment in a given cluster. In addition, we included the NMF cluster gene markers from this study as neutrophil state gene sets. The GMT file containing all genes per pathway used in this analysis will be available on Zenodo, and the lists are included in Supplementary Table S1.

#### NMF clustering analysis

In order to identify neutrophil subtypes, we performed NMF clustering of bulk RNA-Seq samples with CIBERSORTx estimated neutrophil fraction > 50% (mature neutrophils and immature neutrophils combined). We used a previously described Bayesian NMF approach which identified 6 clusters^40,84,85^.

#### Sample pathway scoring

Bulk RNA-seq samples were scored for expression of genes in a gene set according to a previously described method used to control for sample complexity, as we anticipated that cells with higher complexity resulting from contamination from other cell types would have more genes detected and thus score higher for any gene set^82^. Briefly, the score for each sample was defined as the average expression of the genes in the gene set minus the average expression of genes in a control gene set. To define the control gene set, all genes were ranked according to average expression across all samples and divided into 25 bins. Next for each gene in the gene set, 100 genes were selected from the same expression bin to create a gene set with comparable expression levels which is 100-fold larger.

#### Clustering Analysis for Single-cell Blood Neutrophils from Sepsis Patients

The gene expression matrix was imported into R using Seurat 4.0.4. Cells were excluded with fewer than 100 genes. Data were normalized using the NormalizeData function and expression values were scaled using the ScaleData function in Seurat. 40 PCs were selected for building the neighborhood graph. Clustering was performed with the Louvain algorithm with a resolution of 0.6 which resulted in 6 clusters. Cluster markers were determined using the FindMarkers function in Seurat, and p value corrections were performed with the Benjamini-Hochberg method.

#### Neutrophil state network analysis

Neutrophil state gene signatures were taken from the same GMT file used for GSEA analysis in Figure 2F. The network was built using the igraph package in R. Edges were drawn between nodes if the Jaccard index between the two gene signature lists was greater than 0.05. Edge width was scaled according to the overlap coefficient between the gene sets, and nodes were scaled according to gene set size and colored according to the number of neighbors in the graph.

#### Schulte-Schrepping single-cell RNA-seq reanalysis for early-late threshold

The single-cell fresh whole blood neutrophil data from Bonn cohort 2, originally analyzed by Schulte-Schrepping et al.^11^, was reanalyzed for cluster membership according to day using day 11 as the threshold for late disease. For each cluster, we created a running metric for how many cells were classified as “early” by calculating the percentage of cells collected from Day 0 to Day x (Figure S10D).

#### ARDS log fold-change comparisons

Log_2_(fold-change) (LFC) values in blood neutrophil microarray gene expression between non-COVID-19 ARDS patients and healthy volunteers was obtained from the study from Juss et al^44^. Linear regression on the LFC values in ARDS vs. healthy volunteers and severe COVID-19 vs. mild COVID-19 was performed using the lm package in R. To generate a ranked list of genes based on the differences in LFC values, ARDS LFC values were z-scored, and mild vs. severe COVID-19 LFC values were z-scored on each individual day. GSEA was then performed on the lists using the difference in LFC z-score as the ranking metric.

#### Day:Severity interaction analysis

To identify diverging patterns of gene expression between severity groups with time, we built models using DESeq2 for COVID-19-positive samples on Days 0, 3, and 7. The full model included CIBERSORTx estimated cell type fractions, the immunoglobulin score, and the terms for Day, Severity_Max_, and the Day:Severity_Max_ interaction term, while the reduced model did not include the interaction term, and we used the likelihood ratio test in DESeq2 to compare these models. Log(fold-change) values and p-values were extracted to generate a ranked list of genes according to signed p-values for GSEA.

#### Logistic regression models to predict severe COVID-19 on Day 0

Logistic regression models were built using the glm package in R. In order to ensure the stability and interpretability of the coefficients in the model, we included only COVID-19-positive patients on Day 0 who were not immediately discharged from the ED (Acuity_Max_1-4) and who had complete data for ANC, ALC, D-dimer, CRP, LDH, and BMI measured at Day 0. For patients with Acuity_Max_ 1-4, 7 patients had missing clinical data, and these 7 missing patients were not biased towards a particular severity according to Fisher’s exact test. All parameters used were broken into discrete quintiles unless insufficient samples belonged to one category, in which case factor levels were combined in order to minimize the standard error of the coefficient estimation. We combined factor levels for age, LDH, and BMI, leaving 4 factor levels for age and LDH, and 5 factor levels for BMI (BMI was the only category scored from 0 to 5). Models were built according to three tiers of parameters. Model 1: clinical characteristics (age, sex, ethnicity, heart disease, diabetes, hypertension, hyperlipidemia, lung disease, kidney disease, immunocompromised status, BMI), Model 2: clinical characteristics plus clinical laboratory values (ANC, ALC, Creatinine, CRP, D-dimer, LDH), and Model 3: clinical characteristics plus clinical laboratory values plus neutrophil gene signature scores (NMF1, NMF2, NMF3, NMF4, NMF5, NMF6, ARDS Up - Juss, ARDS Down - Juss). ROC curves and AUC values were calculated using the pROC package in R. Significance of model improvement was determined using the likelihood ratio test using the lrtest package in R.

Feature selection for the best predictors on Day 0 of severity within 28 days among the variables used in Model 3 was performed using LASSO with the glmnet package in R with 100 repeats of 5-fold cross validation^26^. Model tuning was performed using the caret package in R. We ranked features according to the number of cross-validation folds in which they were selected for the LASSO model (Figure 3D).

#### Plasma proteomic markers of neutrophil subtypes

To identify plasma proteins associated with neutrophil NMF subtypes, we performed a Wilcoxon rank-sum test for all of the 1472 proteins measured in the Olink plasma proteomic assay between samples from NMF cluster_i_ versus all other clusters (including Neu-Lo). We used the updated Olink proteomics data (https://info.olink.com/broad-covid-study-overview-download) which had the following modifications: 1) scale correction factors were no longer used, and 2) limits of detection were calculated on a per plate basis rather than the whole project. This resulted in the recovery of 43 assays which were not included in the original version; using the new method, no assays had 100% of samples below the limit of detection. Results for each cluster were filtered for p_adj_ > 0.05, first selecting only positive markers (higher protein levels in cluster_i_), and next selecting only negative markers (lower protein levels in cluster_i_). The strongest positive markers were selected by filtering out all markers which did not satisfy the criteria that 1) the highest expression of the protein was in the given NMF cluster and 2) the step ratio, defined as the NPX difference between the given NMF cluster and the second-highest expressing cluster, was at least 0.1. A similar method filtering out markers that did not have the lowest expression in the given NMF cluster and markers with a step ratio for the second lowest cluster of at least 0.1 was used for negative markers. Heatmaps of the protein markers per cluster were generated with the pheatmap package in R, with genes ordered according to p value.

#### Comparison of differential expression and plasma proteomic data

To compare log_2_(fold-change) values on the plasma protein level and neutrophil RNA transcriptional level, we performed differential expression analyses for each. For plasma proteins, we fit linear models using the lm packages in R for each protein using the following clinical covariates: age, sex, ethnicity, heart disease, diabetes, hypertension, hyperlipidemia, pulmonary condition, kidney disease, immunocompromised status. For RNA-seq data, we used DESeq2 differential expression analysis in R with the same clinical covariates as well as the CIBERSORTx estimated cell type fractions. The LFC values were compared for COVID-19-positive vs. COVID-19-negative samples, as well as severe vs. non-severe samples on Days 0, 3, and 7 separately.

#### Ligand-receptor interaction analysis

A curated ligand-receptor pair database from FANTOM5 was used to search for interactions between neutrophil receptors and plasma ligands on either the basis of Severity_Max_ or neutrophil NMF cluster^86^. The database was filtered on ligand-receptor interactions identified as “literature-supported” or “putative”, and was further filtered for receptors with non-zero expression in granulocytes according to the Human Protein Atlas^87^. To identify neutrophil receptors associated with specific NMF clusters, differential expression was performed using DESeq2 for NMF cluster_i_ versus all other clusters irrespective of Day. Only positive gene markers were kept with p_adj_ < 0.05. Differentially expressed receptors which were not unique to a single NMF cluster were excluded. Similarly, differential expression of plasma proteins was performed using lm in R comparing NMF cluster_i_ vs all other clusters (including Neu-Lo), and proteins were kept with p_adj_ < 0.05. Thus a list of potential interactions was generated using the database. To determine whether the neutrophil receptors and plasma proteins were differentially expressed within the same sample rather than the aggregated group, the percentage of samples within a given NMF cluster on a specific day which had higher than mean expression across all COVID-19+ samples of both neutrophil receptor and plasma protein were calculated. In Figures 7A, 7B and Figure S21, ligands matching with multiple receptors were then colored according to the interaction which had the highest percentage of above-mean expression, and secondary interactions were indicated with reduced line width. Plasma ligands were then mapped to the inferred cell-of-origin using single-cell data from bronchoalveolar lavage fluid from COVID-19 patients as previously described^7,26^.

## ACKNOWLEDGMENTS

We acknowledge support from the Ragon Institute of MGH, MIT and Harvard, the Massachusetts Consortium on Pathogen Readiness (MassCPR), the NIH (3R37AI080289-11S1, R01AI146785, U19AI42790-01, U19AI135995-02, U19AI42790-01, 1U01CA260476 – 01, CIVIC75N93019C00052, T32 GM007592), the American Lung Association, and the MGH Executive Committee on Research. A.-C.V. acknowledges funding support from the COVID-19 Clinical Trials Pilot grant from the Executive Committee on Research at MGH, a COVID-19 Chan Zuckerberg Initiative grant (2020-216954), and funds from the Manton Foundation and the Klarman Family Foundation. We thank Arnav Mehta for his help with the proteomics data and for the code which we adapted from Filbin et al. 2021. We also thank Mikael Pittet for his expertise in neutrophil biology.

## AUTHOR CONTRIBUTIONS

Conceptualization: T.J.L., A.L.K.G., S.S.F., P.K., G.A., N.H., and M.S.F.

Methodology: T.J.L., A.L.K.G., S.S.F., P.K., G.A., M.B.G., M.R.F., A.C.V., N.H., and M.S.F.

Validation: T.J.L., P.K., and M.S.F.

Formal Analysis: T.J.L., and S.S.F.

Investigation: T.J.L., A.L.K.G., P.K., I.G., K.R.K., K.M., J.T., M.R-L., B.C.R., N.S., M.F.T., K.M.L-P., B.M.L., B.N.M., N.C.C., H.K.K., C.L.L., J.D.M., E.M.B., P.B.L., R.P.B., B.A.P., M.B.G., M.R.F., A.C.V., and M.S.F.

Resources: M.B.G., M.R.F., A.C.V., G.A., N.H. and M.S.F.

Data Curation: T.J.L., S.S.F., N.H., and M.S.F.

Writing- Original Draft: T.J.L., A.L.K.G., S.S.F., P.K., and M.S.F.

Writing- Review and Editing: T.J.L., A.L.K.G., S.S.F., P.K., I.G., G.A., M.B.G., M.R.F., A.C.V., N.H., and M.S.F.

Visualization: T.J.L., A.L.K.G., S.S.F., P.K., and M.S.F.

Supervision: G.A., N.H., and M.S.F.

## Supplementary Materials

### List of supplementary figures

**Figure S1.** Overview of Cohort and Quality Control Filtration, Related to Figure 1.

**Figure S2.** Estimation of Sample Purity Using CIBERSORTx, Related to Figure 1.

**Figure S3.** Determinants of Neutrophil Sample Purity, Related to Figure 1.

**Figure S4**. Comparison of CIBERSORTx Neutrophil Fractions with Clinical Parameters, Related to Figure 1.

**Figure S5.** The UMAP Landscape of Bulk RNA-seq Neutrophils in the Cohort, and CIBERSORTx Fractions Across Disease Severity, Related to Figure 1.

**Figure S6.** Defining an Immunoglobulin Score to Regress Plasmablast Contamination, Related to Figure 1.

**Figure S7.** Characterizing NMF Clustering Results by COVID-19 Status, Severity, Time, and Sample Purity, Related to Figure 2.

**Figure S8.** Breakdown of Clinical Parameters Across Neutrophil Subtypes, Related to Figure 2.

**Figure S9.** Differential Gene Expression Between Immature NMF Clusters, and NMF Cluster Comparisons with Previously-Defined Neutrophil States, Related to Figure 2.

**Figure S10.** Differentially Expressed Genes between Severe and Non-severe Samples, and Neutrophil State Dynamics, Related to Figure 2.

**Figure S11.** Blood Neutrophils in Severe COVID-19 Transcriptionally Resemble Blood Neutrophils from COVID-19-negative Severe ARDS Patients, Related to Figure 2.

**Figure S12.** Genes and Pathways that Vary with Time According to Severity, Related to Figure 2.

**Figure S13.** Predicting COVID-19 Disease Severity on Day 0, and Comparisons Among Highest Acuity Patients, Related to Figure 3.

**Figure S14.** Patterns of NETosis in RNA and Plasma, Related to Figure 4.

**Figure S15.** Cell-free DNA in COVID-19, Related to Figure 4.

**Figure S16.** Neutrophil Degranulation and T Cell Suppression Mechanisms in COVID-19, Related to Figure 4.

**Figure S17.** Immunoglobulin Levels in Plasma, and Antibody-Dependent Neutrophil Phagocytosis Assay, Related to Figure 5.

**Figure S18.** Differential Effects of IgG versus IgA Antibodies on Neutrophil Effector Functions, Related to Figure 5.

**Figure S19.** Differentially Expressed Plasma Proteins Associated with Neutrophil States, Related to Figure 6.

**Figure S20.** RNA and Protein Log-Fold-Change Comparisons for COVID-19 status and Severity Across Days, and Differentially-expressed Plasma Proteins Associated with Higher IgG or IgA Titers, Related to Figure 6.

**Figure S21.** Ligand-receptor Interaction Analysis for Severe versus Non-severe Patients, Related to Figure 7.

**Figure S22.** Inferring Cell-of-origin for Plasma Ligands Utilizing Single-cell RNA-seq Data, Related to Figure 7.

### List of supplementary tables

**Table S1.** Clinical Metadata, Gene Expression Matrices, Genomic Parameters, and Information for Figure 1 and Related Supplementary Figures S1-S6.

**Table S2.** Information for Figure 2 and Related Supplementary Figures S7-S12.

**Table S3.** Information for Figure 3 and Related Supplementary Figure S13.

**Table S4.** Information for Figure 5 and Related Supplementary Figures S17-S18.

**Table S5.** Information for Figures 6 and 7 and Related Supplementary Figures S19-S22.

