## Supplementary Figures for "Longitudinal characterization of circulating neutrophils uncovers distinct phenotypes associated with disease severity in hospitalized COVID-19 patients"

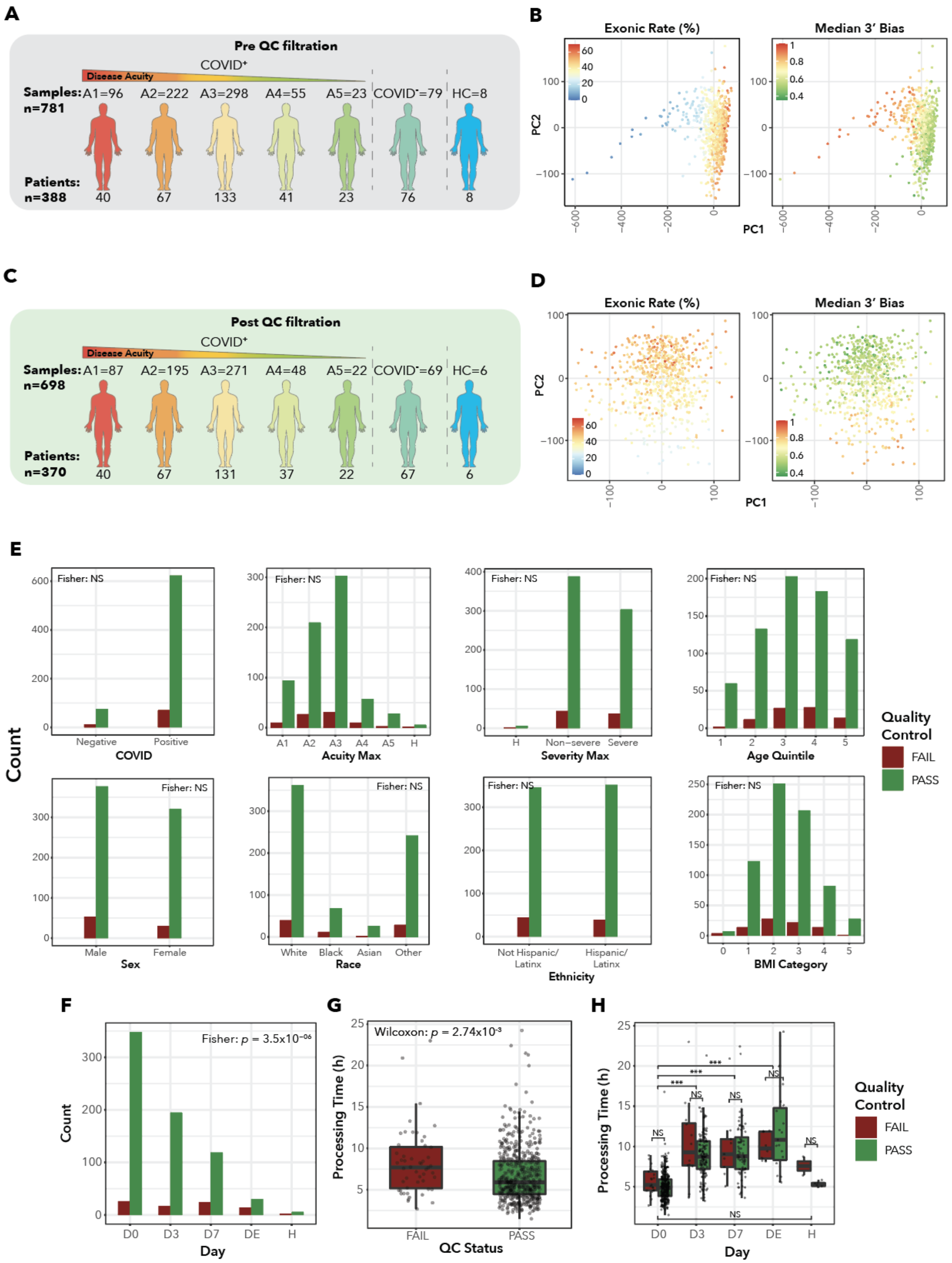

A
Pre QC filtration
COVID+
Disease Acuity
Samples: A1=96 A2=222 A3=298 A4=55 A5=23 COVID-=79 HC=8
n=781
Patients: n=388 40 67 133 41 23 76 8
B
Exonic Rate (%)
Median 3' Bias
PC2
PC1
C
Post QC filtration
COVID+
Disease Acuity
Samples: A1=87 A2=195 A3=271 A4=48 A5=22 COVID-=69 HC=6
n=698
Patients: n=370 40 67 131 37 22 67 6
D
Exonic Rate (%)
Median 3' Bias
PC2
PC1
E
Count
Fisher: NS
COVID
Acuity Max
Severity Max
Age Quintile
Sex
Race
Ethnicity
BMI Category
Quality Control
FAIL
PASS
F
Fisher: p = 3.5x10-06
Count
Day
G
Wilcoxon: p = 2.74x10-3
Processing Time (h)
QC Status
H
Processing Time (h)
Day
Quality Control
FAIL
PASS

**Figure S1. Overview of Cohort and Quality Control Filtration, Related to Figure 1.**
**(A)** Stratification of cohort according to disease status and severity before quality control.
**(B)** Principal Component Analysis (PCA) of all neutrophil-enriched bulk RNA-seq samples before quality control filtration, color-coded by Exonic Rate and Median 3' Bias (left to right).
**(C)** Overview of cohort post-quality control filtration.
**(D)** PCA of all neutrophil-enriched bulk RNA-seq samples passing quality control, color-coded by Exonic Rate and Median 3' Bias (left to right).
**(E)** Bar plots of (left to right) COVID-19 status, $\text{Acuity}_{\text{Max}}$ within 28 days, $\text{Severity}_{\text{Max}}$ within 28 days, Age category, Sex, Race, Ethnicity, and BMI category, split between samples which passed or failed quality control. Fisher's exact test performed to confirm no significant differences in distributions.
**(F)** Bar plots of samples passing or failing quality control by day. Fisher's exact test performed to determine significance.
**(G)** Box plots of sample processing time for samples passing or failing quality control. Wilcoxon rank-sum test performed to determine significance.
**(H)** Box plots of sample processing time versus day of collection, split by quality control status. Wilcoxon rank-sum test performed within day and between days.

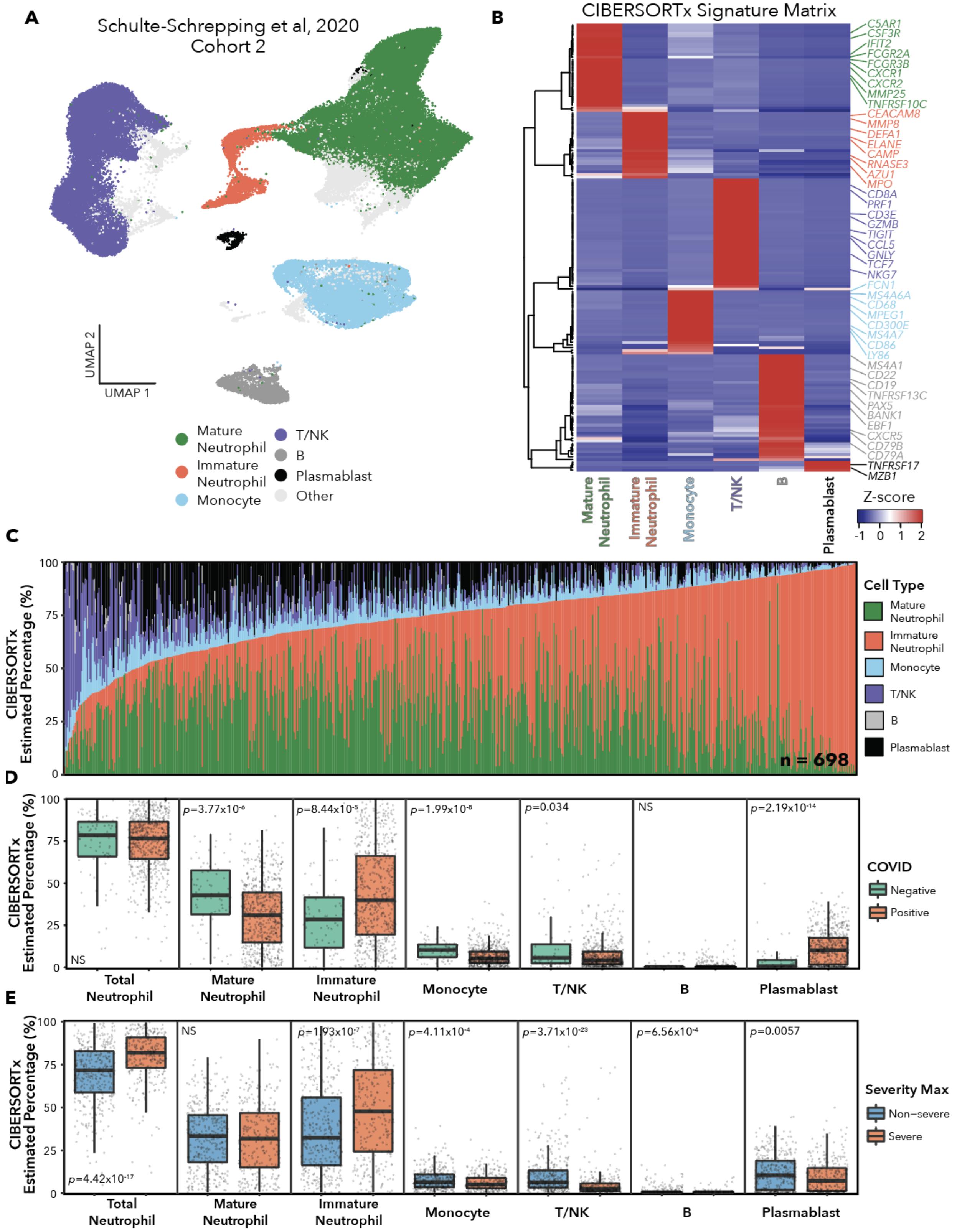
A
Schulte-Schrepping et al, 2020
Cohort 2
UMAP 2
UMAP 1
Mature Neutrophil
Immature Neutrophil
Monocyte
T/NK
B
Plasmablast
Other
B
CIBERSORTx Signature Matrix
C5AR1
CSF3R
IFIT2
FCGR2A
FCGR3B
CXCR1
CXCR2
MMP25
TNFRSF10C
CEACAM8
MMP8
DEFA1
ELANE
CAMP
RNASE3
AZU1
MPO
CD8A
PRF1
CD3E
GZMB
TIGIT
CCL5
GNLY
TCF7
NKG7
FCN1
MS4A6A
CD68
MPEG1
CD300E
MS4A7
CD86
LY86
MS4A1
CD22
CD19
TNFRSF13C
PAX5
BANK1
EBF1
CXCR5
CD79B
CD79A
TNFRSF17
MZB1
Mature Neutrophil
Immature Neutrophil
Monocyte
T/NK
B
Plasmablast
Z-score
-1 0 1 2
C
CIBERSORTx Estimated Percentage (%)
n = 698
Cell Type
Mature Neutrophil
Immature Neutrophil
Monocyte
T/NK
B
Plasmablast
D
CIBERSORTx Estimated Percentage (%)
NS
p=3.77x10-6
p=8.44x10-5
p=1.99x10-8
p=0.034
NS
p=2.19x10-14
Total Neutrophil
Mature Neutrophil
Immature Neutrophil
Monocyte
T/NK
B
Plasmablast
COVID
Negative
Positive
E
CIBERSORTx Estimated Percentage (%)
p=4.42x10-17
NS
p=1.93x10-7
p=4.11x10-4
p=3.71x10-23
p=6.56x10-4
p=0.0057
Total Neutrophil
Mature Neutrophil
Immature Neutrophil
Monocyte
T/NK
B
Plasmablast
Severity Max
Non−severe
Severe

**Figure S2. Estimation of Sample Purity Using CIBERSORTx, Related to Figure 1.**
**(A)** UMAP (Uniform Manifold Approximation and Projection) plot of single-cell RNA-seq data of fresh whole blood from COVID-19-positive patients and controls from Schulte-Schrepping et al. Cohort 2. Cells are colored according to their major lineage (neutrophil, monocyte, T/NK, B, Plasmablast, Other), with neutrophils split between mature and immature.
**(B)** CIBERSORTx scaled expression signature matrix, generated from pseudobulked Schulte-Schrepping cell types using the “Create Signature Matrix” module, used to deconvolute the neutrophil-enriched bulk RNA-seq samples.
**(C)** Distribution of the CIBERSORTx estimated cell type percentages for each sample. Each column is a single sample, and columns are ordered by increasing Total Neutrophil content (Immature Neutrophil Fraction + Mature Neutrophil Fraction).
**(D)-(E)** Box plots showing the distribution across all samples of CIBERSORTx Estimated Percentages of (left to right) Total Neutrophils, Mature Neutrophils, Immature Neutrophils, Monocytes, T/NK, B, and Plasmablasts, split by (D) COVID-19 status and (E) $Severity_{Max}$ within 28 days. Wilcoxon rank-sum test performed for significance.

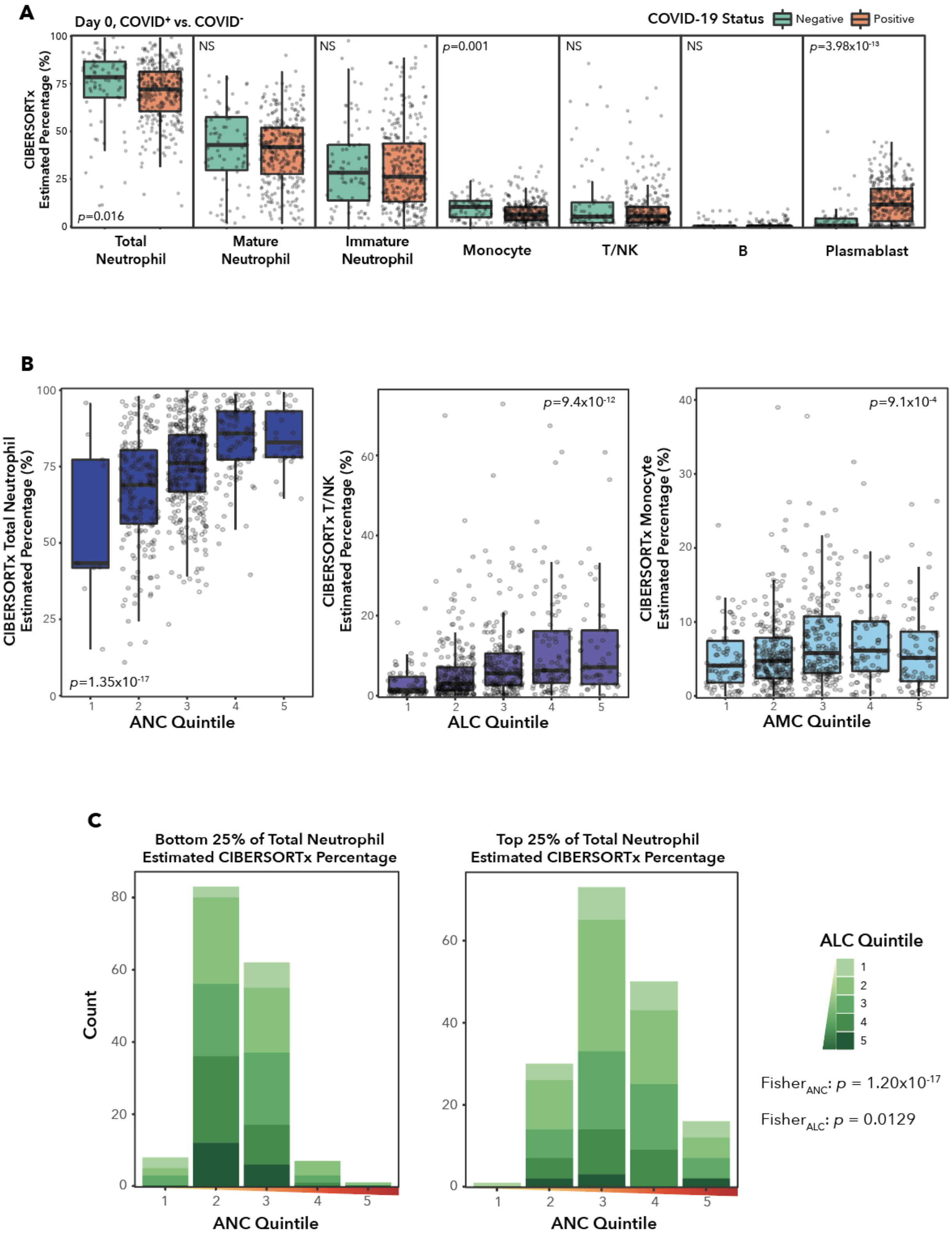
A
Day 0, COVID+ vs. COVID-
COVID-19 Status
Negative
Positive
CIBERSORTx Estimated Percentage (%)
p=0.016
NS
NS
p=0.001
NS
NS
p=3.98x10-13
Total Neutrophil
Mature Neutrophil
Immature Neutrophil
Monocyte
T/NK
B
Plasmablast
B
CIBERSORTx Total Neutrophil Estimated Percentage (%)
p=1.35x10-17
ANC Quintile
CIBERSORTx T/NK Estimated Percentage (%)
p=9.4x10-12
ALC Quintile
CIBERSORTx Monocyte Estimated Percentage (%)
p=9.1x10-4
AMC Quintile
C
Bottom 25% of Total Neutrophil Estimated CIBERSORTx Percentage
Top 25% of Total Neutrophil Estimated CIBERSORTx Percentage
Count
ANC Quintile
ANC Quintile
ALC Quintile
Fisher ANC: p = 1.20x10-17
Fisher ALC: p = 0.0129

**Figure S3. Determinants of Neutrophil Sample Purity, Related to Figure 1.**
**(A)** Box plots comparing CIBERSORTx Estimated Cell Type Percentages across COVID-19 status on Day 0. Wilcoxon rank-sum test performed to test significance.
**(B)** Box plots comparing (left) absolute neutrophil counts to CIBERSORTx Estimated Total Neutrophil Percentage, (middle) absolute lymphocyte counts to CIBERSORTx Estimated T/NK Percentage, and (right) absolute monocyte counts to CIBERSORTx Estimated Monocyte Percentage. Kruskal-Wallis test performed to determine significance.
**(C)** Stacked bar plots showing the distribution of absolute neutrophil counts within (left) the bottom 25% of CIBERSORTx Total Neutrophil Estimated Percentage or (right) the top 25% of CIBERSORTx Total Neutrophil Estimated Percentage. Bars are colored according to absolute lymphocyte counts. Fisher's exact test used to compare distributions.

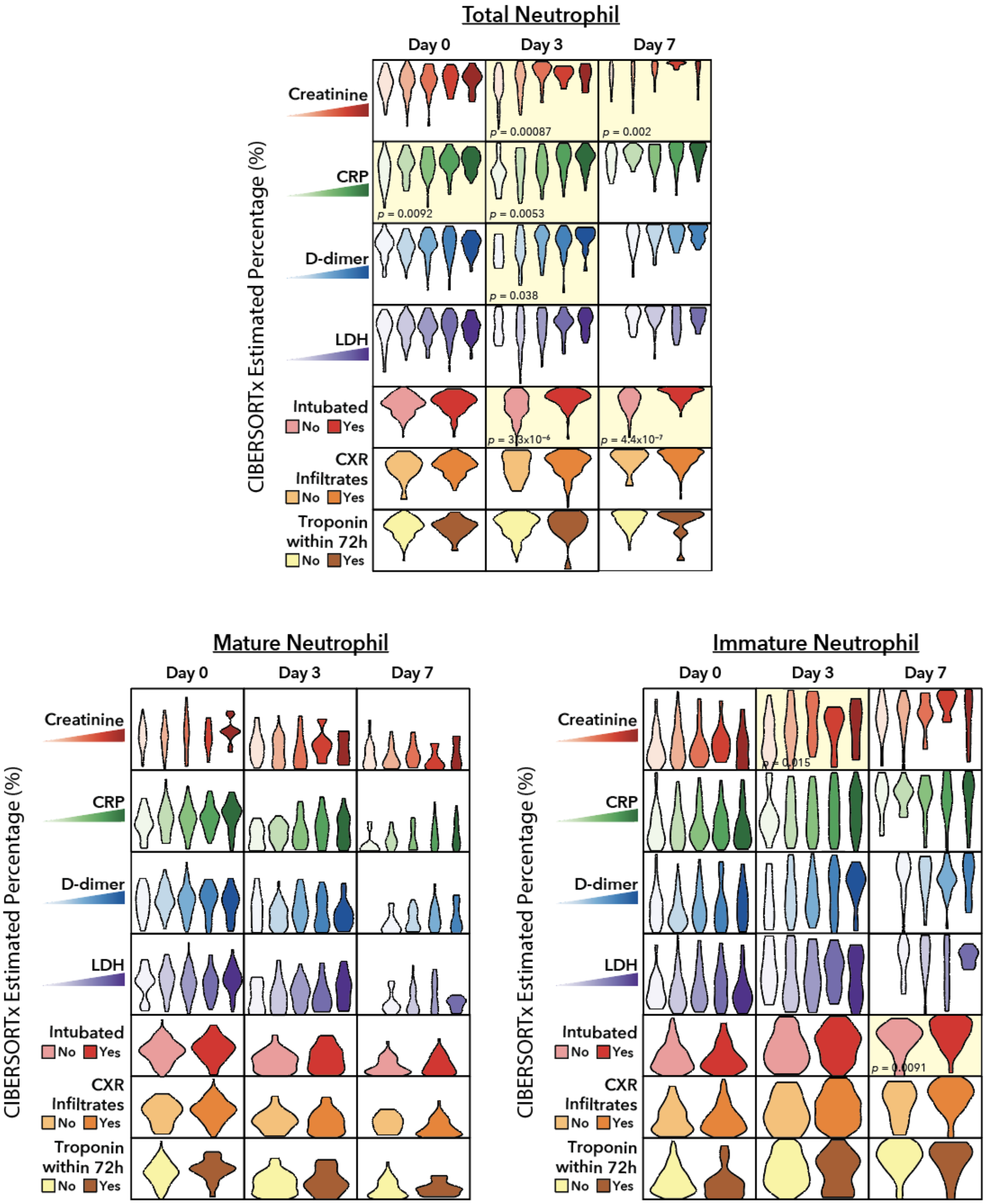
Total Neutrophil
Day 0
Day 3
Day 7
CIBERSORTx Estimated Percentage (%)
Creatinine
CRP
D-dimer
LDH
Intubated
No
Yes
CXR Infiltrates
No
Yes
Troponin within 72h
No
Yes
p = 0.00087
p = 0.002
p = 0.0092
p = 0.0053
p = 0.038
p = 4.4x10⁻⁷
Mature Neutrophil
Day 0
Day 3
Day 7
CIBERSORTx Estimated Percentage (%)
Creatinine
CRP
D-dimer
LDH
Intubated
No
Yes
CXR Infiltrates
No
Yes
Troponin within 72h
No
Yes
Immature Neutrophil
Day 0
Day 3
Day 7
CIBERSORTx Estimated Percentage (%)
Creatinine
CRP
D-dimer
LDH
Intubated
No
Yes
CXR Infiltrates
No
Yes
Troponin within 72h
No
Yes

**Figure S4**. **Comparison of CIBERSORTx Neutrophil Fractions with Clinical Parameters, Related to Figure 1.**

Violin plots comparing CIBERSORTx (top) Total Neutrophil, (bottom left) Mature Neutrophil, and (bottom right) Immature Neutrophil Percentages with (top to bottom) Creatinine, CRP, D-dimers, LDH, Intubation status, CXR infiltrates, and Troponin detection within 72 hours, for COVID-19-positive samples on Days 0, 3, or 7. Cells highlighted in yellow show significant differences in CIBERSORTx percentages across the clinical categories. Kruskal-Wallis test used for ordinal variables, and Wilcoxon rank-sum test used for binary variables. P values shown are corrected for multiple comparisons (FDR).

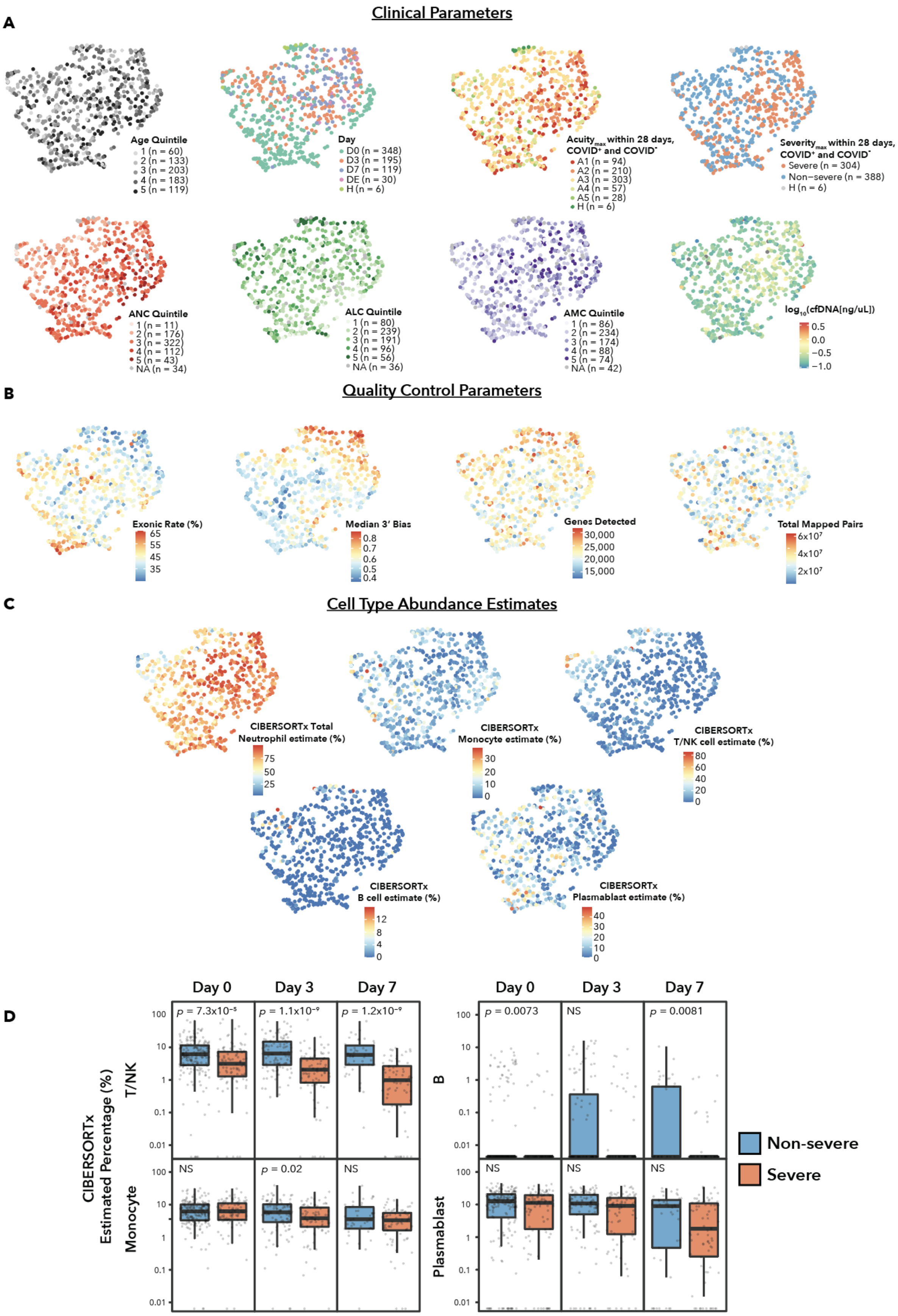
Clinical Parameters
A
Age Quintile
1 (n = 60)
2 (n = 133)
3 (n = 203)
4 (n = 183)
5 (n = 119)
Day
D0 (n = 348)
D3 (n = 195)
D7 (n = 119)
DE (n = 30)
H (n = 6)
Acuity max within 28 days, COVID+ and COVID-
A1 (n = 94)
A2 (n = 210)
A3 (n = 303)
A4 (n = 57)
A5 (n = 28)
H (n = 6)
Severity max within 28 days, COVID+ and COVID-
Severe (n = 304)
Non−severe (n = 388)
H (n = 6)
ANC Quintile
1 (n = 11)
2 (n = 176)
3 (n = 322)
4 (n = 112)
5 (n = 43)
NA (n = 34)
ALC Quintile
1 (n = 80)
2 (n = 239)
3 (n = 191)
4 (n = 96)
5 (n = 56)
NA (n = 36)
AMC Quintile
1 (n = 86)
2 (n = 234)
3 (n = 174)
4 (n = 88)
5 (n = 74)
NA (n = 42)
log10(cfDNA[ng/uL])
0.5
0.0
−0.5
−1.0
B
Quality Control Parameters
Exonic Rate (%)
65
55
45
35
Median 3' Bias
0.8
0.7
0.6
0.5
0.4
Genes Detected
30,000
25,000
20,000
15,000
Total Mapped Pairs
6x10^7
4x10^7
2x10^7
C
Cell Type Abundance Estimates
CIBERSORTx Total Neutrophil estimate (%)
75
50
25
CIBERSORTx Monocyte estimate (%)
30
20
10
0
CIBERSORTx T/NK cell estimate (%)
80
60
40
20
0
CIBERSORTx B cell estimate (%)
12
8
4
0
CIBERSORTx Plasmablast estimate (%)
40
30
20
10
0
D
Day 0
Day 3
Day 7
CIBERSORTx Estimated Percentage (%)
T/NK
Monocyte
B
Plasmablast
p = 7.3x10^-5
p = 1.1x10^-9
p = 1.2x10^-9
NS
p = 0.02
NS
p = 0.0073
NS
p = 0.0081
NS
NS
NS
100
10
1
0.1
0.01
Non-severe
Severe

**Figure S5. The UMAP Landscape of Bulk RNA-seq Neutrophils in the Cohort, and CIBERSORTx Fractions Across Disease Severity, Related to Figure 1.**

**(A)** UMAP plots colored by the following clinical parameters (left to right): Age category, Day of blood draw, $\text{Acuity}_{\text{Max}}$ within 28 days (including COVID-19-negative Acuity categorizations), $\text{Severity}_{\text{Max}}$ within 28 days (including COVID-19-negative Severity categorizations), absolute neutrophil count category, absolute lymphocyte count category, absolute monocyte count category, and cell-free DNA concentration in plasma.

**(B)** UMAP plots colored by the following quality control parameters (left to right): Exonic Rate, Median 3' Bias, Genes Detected, and Total Mapped Pairs, according to RNA-SeQC.

**(C)** UMAP plots colored by the following CIBERSORTx Cell Type Percentage Estimates (left to right): Total Neutrophil, Monocyte, T/NK, B, and Plasmablast.

**(D)** Box plots of CIBERSORTx Estimated Cell Type Percentages for T/NK, Monocyte, B, and Plasmablast, broken down by $\text{Severity}_{\text{Max}}$ and day. Wilcoxon rank-sum test used to determine significance.

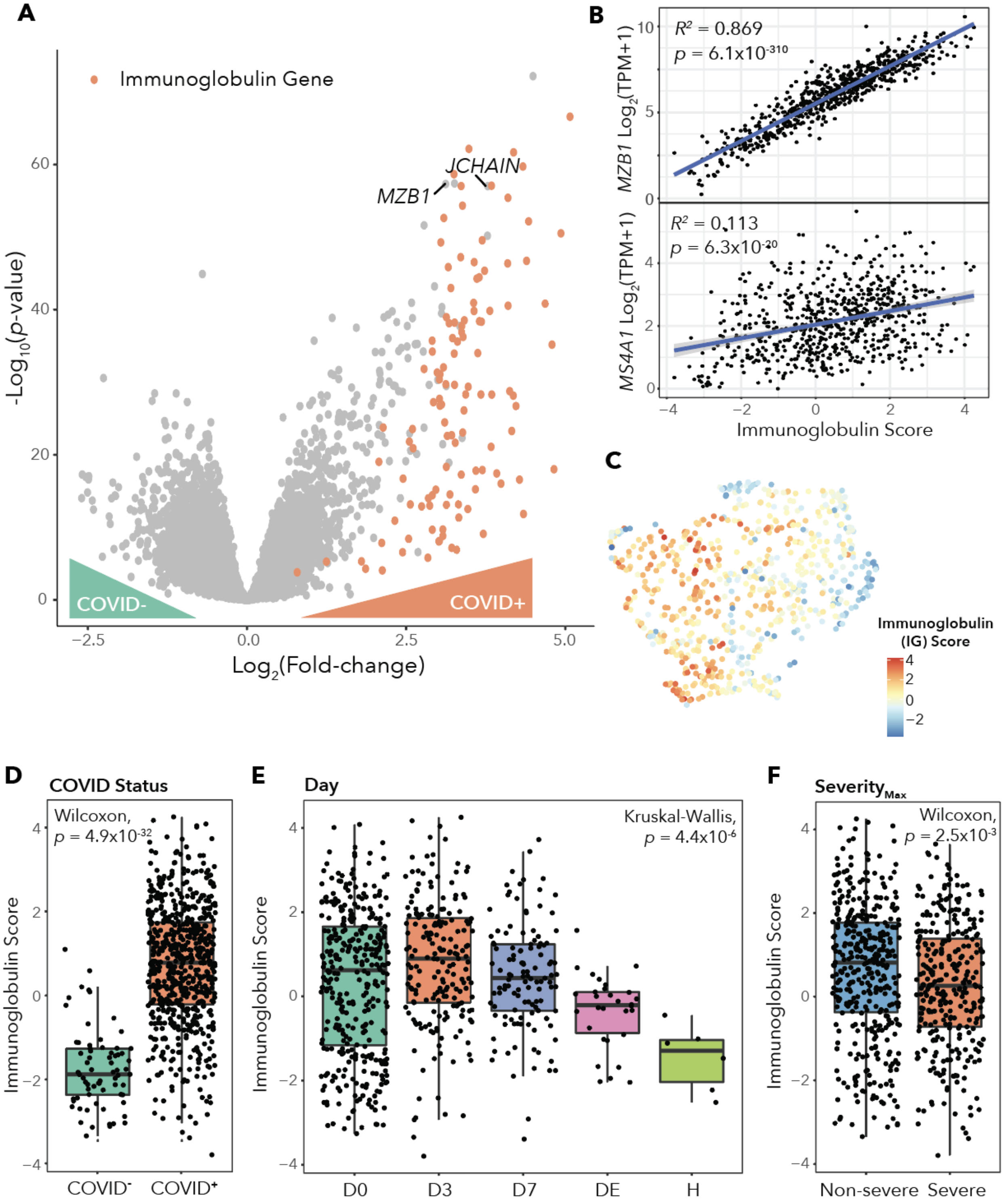
A
Immunoglobulin Gene
JCHAIN
MZB1
COVID-
COVID+
-Log10(p-value)
Log2(Fold-change)
B
R² = 0.869
p = 6.1x10-310
MZB1 Log2(TPM+1)
R² = 0.113
p = 6.3x10-20
MS4A1 Log2(TPM+1)
Immunoglobulin Score
C
Immunoglobulin (IG) Score
D
COVID Status
Wilcoxon, p = 4.9x10-32
Immunoglobulin Score
COVID-
COVID+
E
Day
Kruskal-Wallis, p = 4.4x10-6
D0
D3
D7
DE
H
F
SeverityMax
Wilcoxon, p = 2.5x10-3
Non-severe
Severe

**Figure S6. Defining an Immunoglobulin Score to Regress Plasmablast Contamination, Related to Figure 1.**

**(A)** Volcano plot showing differentially expressed genes between COVID-19-positive and COVID-19-negative samples on Day 0. Immunoglobulin genes (all of which are used in the score) are highlighted in red, and plasmablast marker genes *MZB1* and *JCHAIN* are annotated.

**(B)** Scatter plots and linear regression of immunoglobulin score versus $log_2$(TPM+1) expression of (top) plasmablast marker gene *MZB1* and (bottom) B cell marker *MS4A1*.

**(C)** UMAP plot of all bulk RNA-seq samples color-coded by immunoglobulin score.

**(D)** Box plots comparing immunoglobulin score across COVID-19 status for all time points. Wilcoxon rank-sum test performed to determine significance.

**(E)** Box plots comparing immunoglobulin score across Day for all samples. Kruskal-Wallis test used to determine significance.

**(F)** Box plot comparing immunoglobulin score across $Severity_{Max}$ within 28 days for all samples. Wilcoxon rank-sum test performed to determine significance.

A

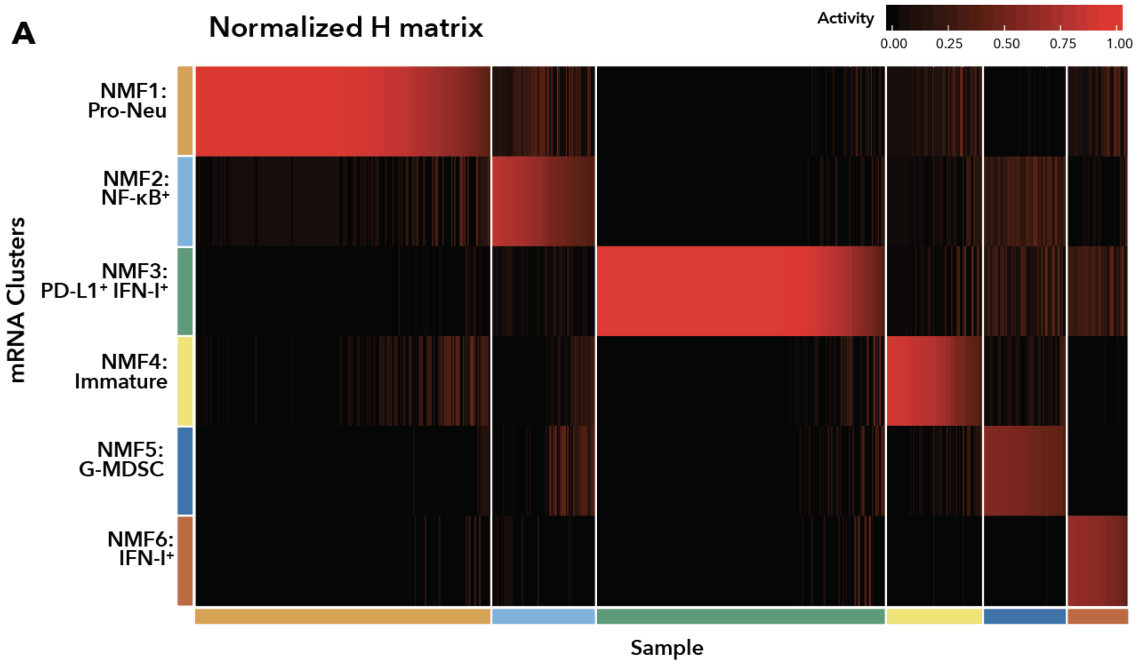


B

| | NMF1 | NMF2 | NMF3 | NMF4 | NMF5 | NMF6 | Neu-Lo |
|---|---|---|---|---|---|---|---|
| COVID⁺ | 183 | 53 | 190 | 64 | 40 | 42 | 51 |
| COVID⁻ | 19 | 18 | 7 | 2 | 17 | 0 | 12 |

C

| | NMF1 | NMF2 | NMF3 | NMF4 | NMF5 | NMF6 | Neu-Lo |
|---|---|---|---|---|---|---|---|
| Severe | 72 | 46 | 86 | 44 | 38 | 7 | 11 |
| Non-severe | 130 | 25 | 111 | 22 | 19 | 35 | 52 |

D

| | NMF1 | NMF2 | NMF3 | NMF4 | NMF5 | NMF6 | Neu-Lo |
|---|---|---|---|---|---|---|---|
| A1 | 16 | 12 | 36 | 8 | 14 | 5 | 3 |
| A2 | 56 | 34 | 50 | 36 | 24 | 2 | 8 |
| A3 | 106 | 20 | 87 | 20 | 17 | 24 | 29 |
| A4 | 12 | 4 | 13 | 2 | 2 | 6 | 18 |
| A5 | 7 | 1 | 11 | 0 | 0 | 5 | 4 |
| H | 5 | 0 | 0 | 0 | 0 | 0 | 1 |

E

| | NMF1 | NMF2 | NMF3 | NMF4 | NMF5 | NMF6 | Neu-Lo |
|---|---|---|---|---|---|---|---|
| Day 0 | 45 | 24 | 168 | 12 | 28 | 34 | 37 |
| Day 3 | 78 | 26 | 22 | 25 | 17 | 6 | 21 |
| Day 7 | 56 | 18 | 6 | 27 | 6 | 2 | 4 |
| Day E | 18 | 3 | 1 | 2 | 6 | 0 | 0 |
| H | 5 | 0 | 0 | 0 | 0 | 0 | 1 |

F

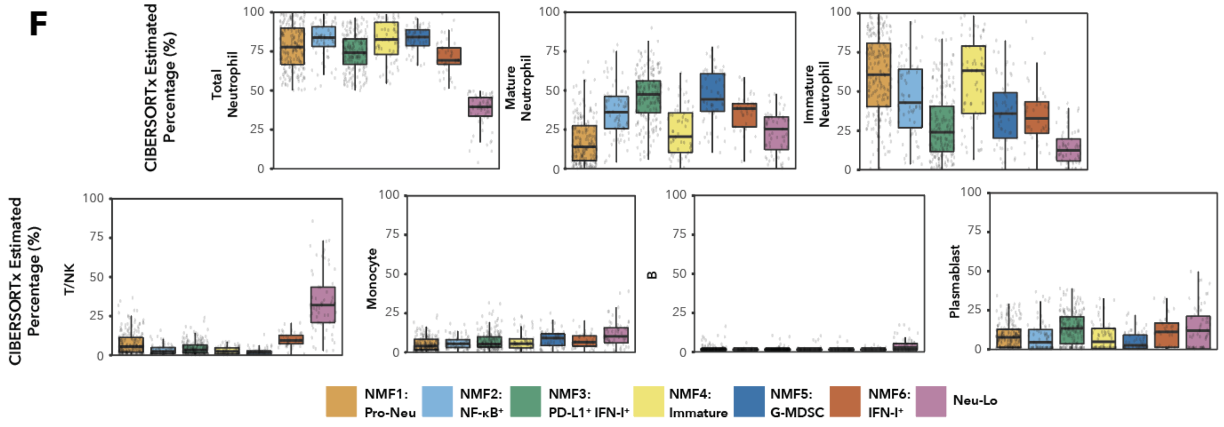

**Figure S7. Characterizing NMF Clustering Results by COVID-19 Status, Severity, Time, and Sample Purity, Related to Figure 2.**

**(A)** NMF Normalized H matrix of neutrophil-enriched bulk RNA-seq samples with CIBERSORTx Estimated Total Neutrophil Percentage above 50%. Clustering identified 6 subtypes. Activity corresponds to the probability that a sample is included in a given cluster. Samples are ordered according to activity value within a given cluster.

**(B)-(E)** Tables showing the distribution of samples in each cluster across (C) COVID-19 status, (D) Severity$_{Max}$ within 28 days (includes COVID-19-negative samples), (E) Acuity$_{Max}$ within 28 days (includes COVID-19-negative samples), and (F) day of blood draw.

**(F)** Box plots of CIBERSORTx Estimated Cell Type Percentage for (left to right) Total Neutrophil, Mature Neutrophil, Immature Neutrophil, T/NK, Monocyte, B, and Plasmablast according to NMF subtype for all samples.

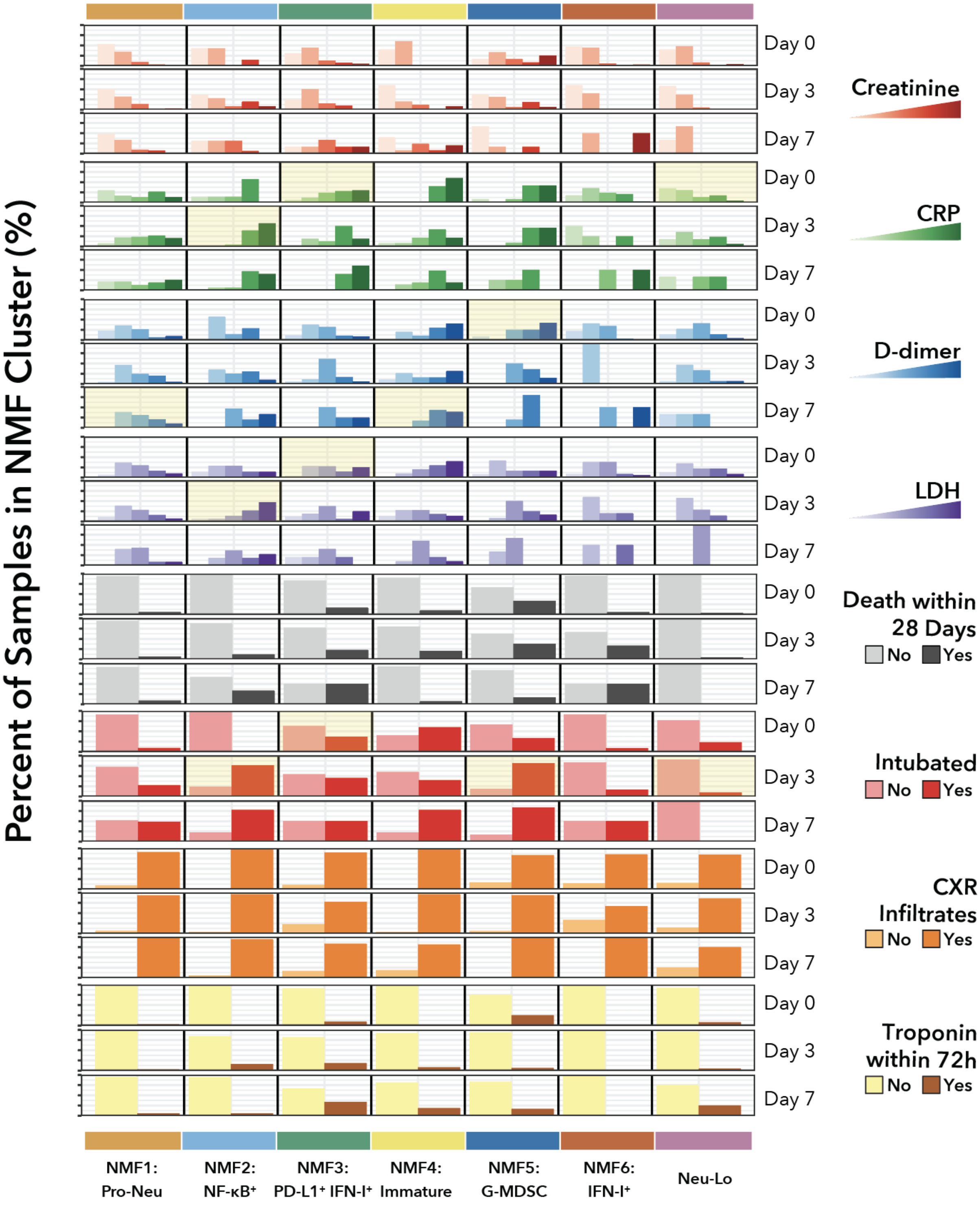
Percent of Samples in NMF Cluster (%)
Day 0
Day 3
Day 7
Creatinine
CRP
D-dimer
LDH
Death within 28 Days
No
Yes
Intubated
No
Yes
CXR Infiltrates
No
Yes
Troponin within 72h
No
Yes
NMF1: Pro-Neu
NMF2: NF-κB⁺
NMF3: PD-L1⁺ IFN-I⁺
NMF4: Immature
NMF5: G-MDSC
NMF6: IFN-I⁺
Neu-Lo

**Figure S8. Breakdown of Clinical Parameters Across Neutrophil Subtypes, Related to Figure 2.**
Bar plots of the following clinical parameters per NMF subtype, broken down by day (top to bottom): Creatinine, CRP, D-dimer, LDH, Death with 28 days, Intubated, CXR Infiltrates, Troponin detected in blood within 72 hours. Plots highlighted in yellow show significant differences from the distribution of parameters outside the selected cluster according to Fisher's exact test, FDR $q < 0.05$.

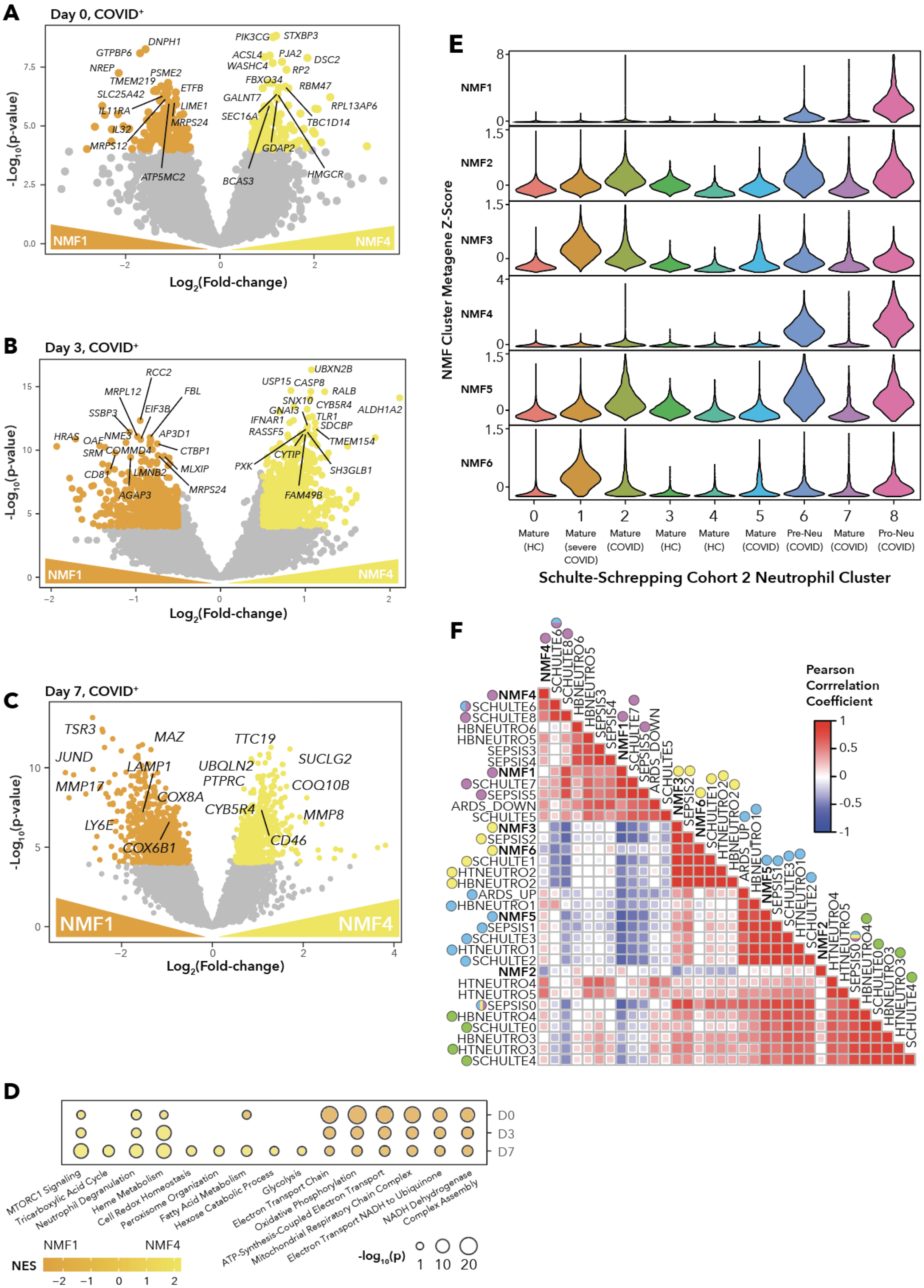
A
Day 0, COVID+
DNPH1
GTPBP6
NREP
PSME2
TMEM219
ETFB
SLC25A42
IL11RA
LIME1
MRPS24
IL32
MRPS12
ATP5MC2
PIK3CG
STXBP3
ACSL4
PJA2
DSC2
WASHC4
RP2
FBXO34
RBM47
GALNT7
RPL13AP6
SEC16A
TBC1D14
GDAP2
BCAS3
HMGCR
NMF1
NMF4
-Log10(p-value)
Log2(Fold-change)
B
Day 3, COVID+
RCC2
MRPL12
FBL
SSBP3
EIF3B
HRAS
OAF
NME3
AP3D1
SRM
COMMD4
CTBP1
CD81
LMNB2
MLXIP
AGAP3
MRPS24
UBXN2B
USP15
CASP8
RALB
SNX10
CYB5R4
GNAI3
ALDH1A2
IFNAR1
TLR1
RASSF5
SDCBP
TMEM154
CYTIP
PXK
SH3GLB1
FAM49B
NMF1
NMF4
-Log10(p-value)
Log2(Fold-change)
C
Day 7, COVID+
TSR3
MAZ
TTC19
JUND
SUCLG2
LAMP1
UBQLN2
MMP17
PTPRC
COQ10B
COX8A
CYB5R4
MMP8
LY6E
CD46
COX6B1
NMF1
NMF4
-Log10(p-value)
Log2(Fold-change)
D
D0
D3
D7
MTORC1 Signaling
Tricarboxylic Acid Cycle
Neutrophil Degranulation
Heme Metabolism
Cell Redox Homeostasis
Peroxisome Organization
Fatty Acid Metabolism
Hexose Catabolic Process
Glycolysis
Electron Transport Chain
Oxidative Phosphorylation
ATP-Synthesis-Coupled Electron Transport
Mitochondrial Respiratory Chain Complex
Electron Transport NADH to Ubiquinone
NADH Dehydrogenase Complex Assembly
NMF1
NMF4
NES
-log10(p)
E
NMF1
NMF2
NMF3
NMF4
NMF5
NMF6
NMF Cluster Metagene Z-Score
0 Mature (HC)
1 Mature (severe COVID)
2 Mature (COVID)
3 Mature (HC)
4 Mature (HC)
5 Mature (COVID)
6 Pre-Neu (COVID)
7 Mature (COVID)
8 Pro-Neu (COVID)
Schulte-Schrepping Cohort 2 Neutrophil Cluster
F
Pearson Corrrelation Coefficient
NMF4
SCHULTE6
SCHULTE8
HBNEUTRO6
HBNEUTRO5
SEPSIS3
SEPSIS4
NMF1
SCHULTE7
SEPSIS5
ARDS_DOWN
SCHULTE5
NMF3
SEPSIS2
NMF6
SCHULTE1
HTNEUTRO2
HBNEUTRO2
ARDS_UP
HBNEUTRO1
NMF5
SEPSIS1
SCHULTE3
HTNEUTRO1
SCHULTE2
NMF2
HTNEUTRO4
HTNEUTRO5
SEPSIS0
HBNEUTRO4
SCHULTE0
HBNEUTRO3
HTNEUTRO3
SCHULTE4

**Figure S9. Differential Gene Expression Between Immature NMF Clusters, and NMF Cluster Comparisons with Previously-Defined Neutrophil States, Related to Figure 2.**
**(A)-(C)** Volcano plots showing differentially expressed genes between COVID-19-positive samples in clusters NMF1 versus NMF4 on (A) Day 0, (B) Day 3, and (C) Day 7. Color-coded points indicate $\log_2$(fold-change) > 0.5 and $p < 10^{-4}$.
**(D)** Gene set enrichment analysis on genes differentially expressed between COVID-19-positive samples in clusters NMF1 versus NMF4 on Days 0, 3, and 7, specifically highlighting metabolic pathways. Bubble size corresponds to $-\log_{10}(p)$ and color corresponds to NES.
**(E)** Violin plots of the metagene z-score for each NMF cluster signature across the Schulte-Schrepping single-cell fresh whole blood neutrophil data.
**(F)** Pairwise Pearson correlation heatmap for the Z-scores of each gene set on all samples in the cohort. Color-coded dots indicate the network group membership from Figure 2C.

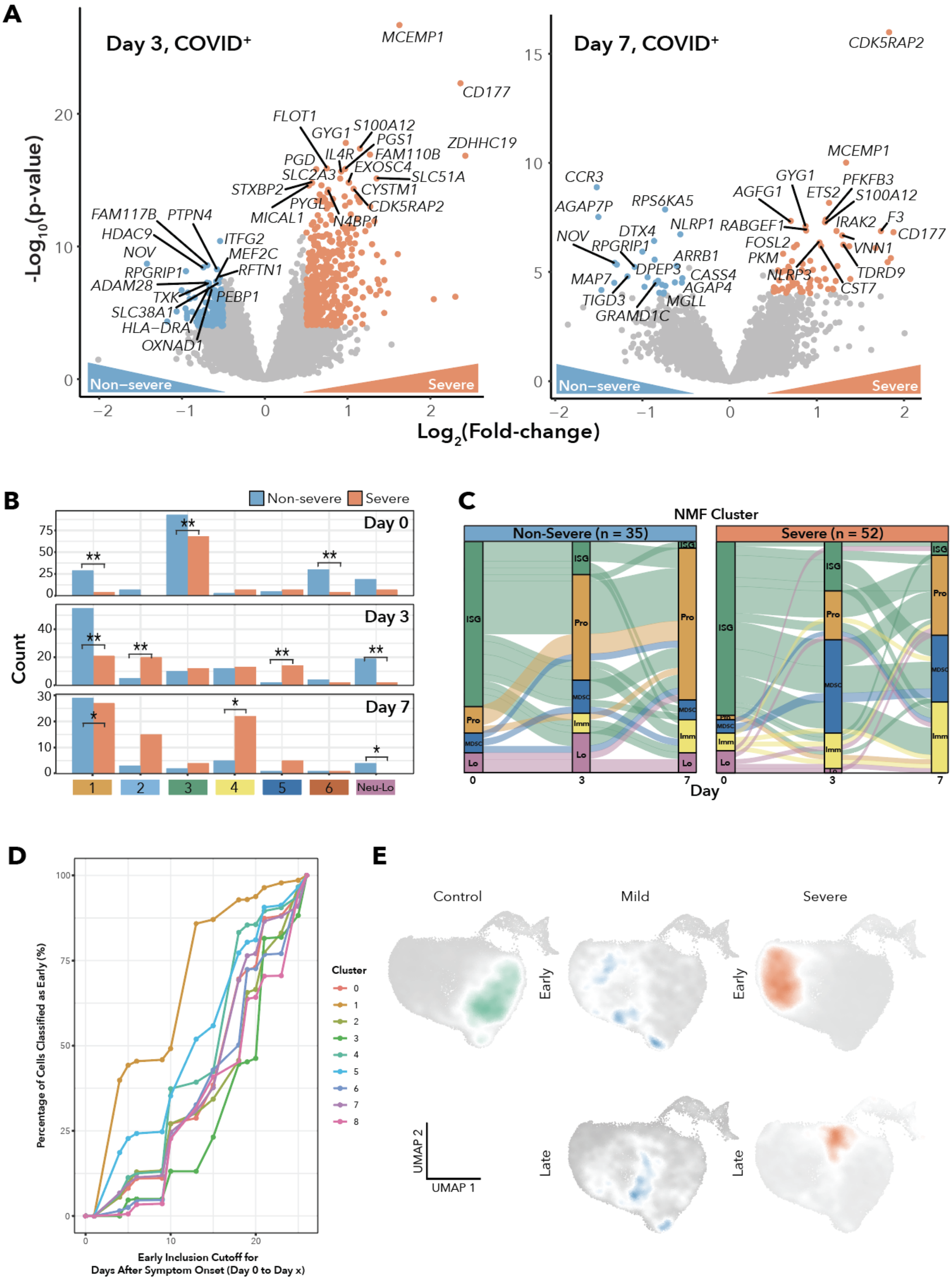
A
Day 3, COVID+
Day 7, COVID+
-Log10(p-value)
Log2(Fold-change)
Non−severe
Severe
MCEMP1
CD177
ZDHHC19
CDK5RAP2
B
Non-severe
Severe
Day 0
Day 3
Day 7
Count
Neu-Lo
C
NMF Cluster
Non-Severe (n = 35)
Severe (n = 52)
ISG
Pro
MDSC
Imm
Lo
Day
D
Percentage of Cells Classified as Early (%)
Early Inclusion Cutoff for
Days After Symptom Onset (Day 0 to Day x)
Cluster
E
Control
Mild
Severe
Early
Late
UMAP 1
UMAP 2

**Figure S10. Differentially Expressed Genes between Severe and Non-severe Samples, and Neutrophil State Dynamics, Related to Figure 2.**

**(A)** Volcano plots showing differentially expressed genes between COVID-19-positive severe and non-severe samples on (left) Day 3 and (right) Day 7. Color-coded points indicate $log_2$(fold-change) > 0.5 and $p < 10^{-4}$.

**(B)** Bar plots showing the number of samples per NMF cluster per day, separated by $Severity_{Max}$. Asterisks denote significance according to Fisher's exact test: single asterisk $p < 0.05$ and double asterisk $p < 0.01$.

**(C)** Alluvial diagrams displaying the change in NMF cluster membership over time for patients who had all three blood draws which all passed quality control, split by non-severe (n = 35 patients) and severe (n = 52 patients). "ISG" indicates NMF3 and NMF6, "Pro" indicates NMF1, "MDSC" indicates NMF5, "Immature" indicates NMF4, and "Lo" indicates samples with CIBERSORTx Estimated Total Neutrophil Percentage less than 50%.

**(D)** Line chart showing the percentage of cells within a cluster that are categorized as early as a function of inclusion cutoff for days following symptom onset for the Schulte-Schrepping Cohort 2 fresh whole-blood neutrophil data. Data shown are for all COVID-19-positive samples. The early cutoff was defined as 0-10 days after disease onset by Schulte-Schrepping et al.

**(E)** Density plots of healthy controls, disease severity, and time point overlain on the Schulte-Schrepping Cohort 2 neutrophil UMAP, classifying early time points as days 0-13 following symptom onset.

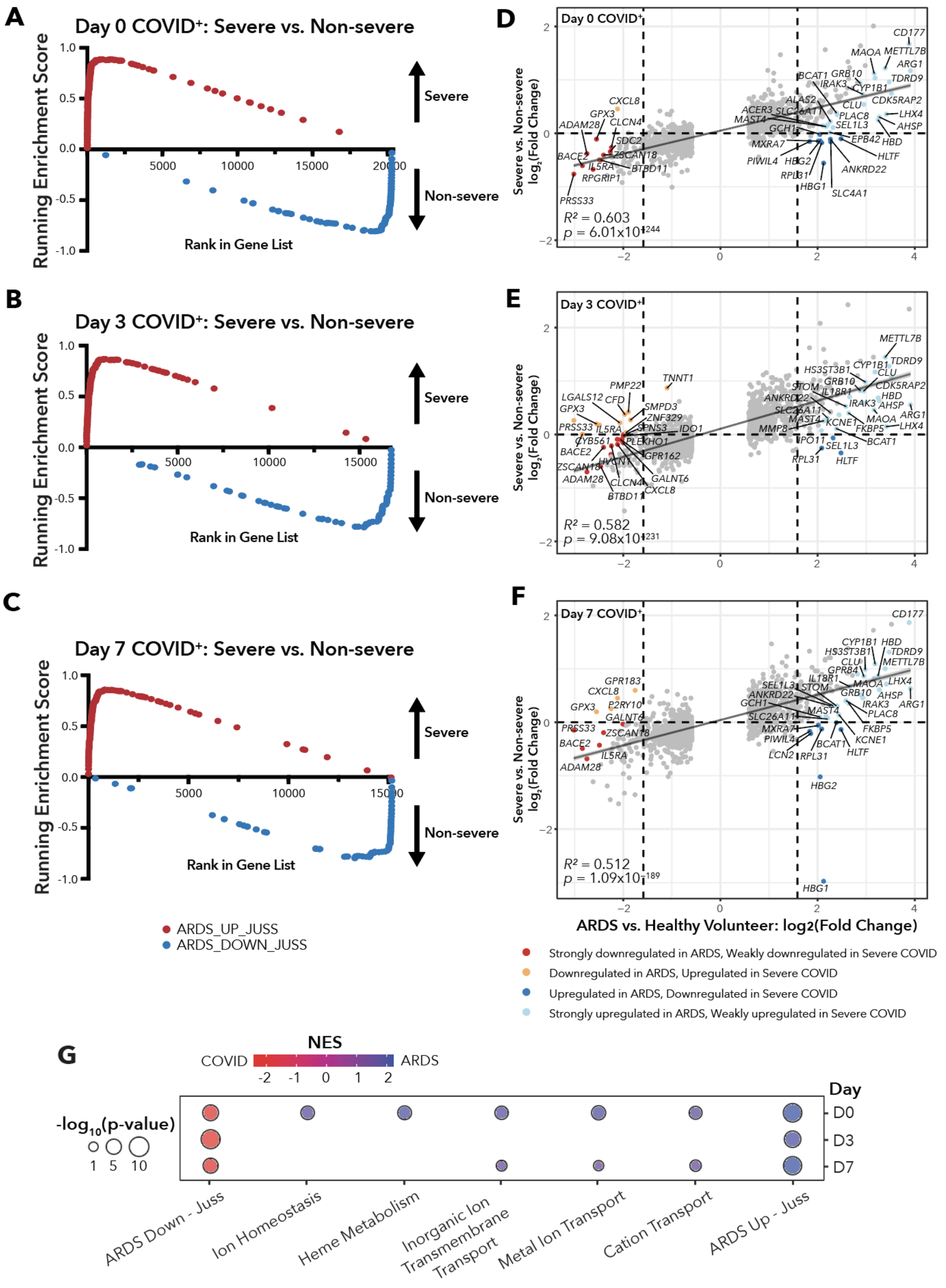
A
Day 0 COVID+: Severe vs. Non-severe
Running Enrichment Score
Rank in Gene List
Severe
Non-severe
B
Day 3 COVID+: Severe vs. Non-severe
Running Enrichment Score
Rank in Gene List
Severe
Non-severe
C
Day 7 COVID+: Severe vs. Non-severe
Running Enrichment Score
Rank in Gene List
Severe
Non-severe
ARDS_UP_JUSS
ARDS_DOWN_JUSS
D
Day 0 COVID+
R² = 0.603
p = 6.01x10-244
E
Day 3 COVID+
R² = 0.582
p = 9.08x10-231
F
Day 7 COVID+
R² = 0.512
p = 1.09x10-189
Severe vs. Non-severe log2(Fold Change)
ARDS vs. Healthy Volunteer: log2(Fold Change)
Strongly downregulated in ARDS, Weakly downregulated in Severe COVID
Downregulated in ARDS, Upregulated in Severe COVID
Upregulated in ARDS, Downregulated in Severe COVID
Strongly upregulated in ARDS, Weakly upregulated in Severe COVID
G
NES
COVID
ARDS
-log10(p-value)
Day
D0
D3
D7
ARDS Down - Juss
Ion Homeostasis
Heme Metabolism
Inorganic Ion Transmembrane Transport
Metal Ion Transport
Cation Transport
ARDS Up - Juss

**Figure S11. Blood Neutrophils in Severe COVID-19 Transcriptionally Resemble Blood Neutrophils from COVID-19-negative Severe ARDS Patients, Related to Figure 2.**
**(A)-(C)** Gene set enrichment plots for the ARDS_UP_JUSS and ARDS_DOWN_JUSS gene sets among differentially expressed genes between COVID-19-positive severe and non-severe samples on (A) Day 0, (B) Day 3, and (C) Day 7. The ARDS gene signatures are defined as genes which are altered with fold-change > 3 in ARDS blood neutrophils compared to healthy volunteers.
**(D)-(F)** Scatter plots and linear regression comparing $log_2$(fold-change) of COVID-19-positive severe versus non-severe on (D) Day 0, (E) Day 3, and (F) Day 7 to $log_2$(fold-change) of ARDS versus Healthy Volunteers at its only time point. For each gene, the difference between the ARDS and COVID-19 fold-changes was calculated. Genes with a difference in fold-change greater than two standard deviations are color-coded according to the legend.
**(G)** Gene set enrichment analysis on the gene list ranked according to differences in fold-changes between COVID-19-negative ARDS and COVID-19.

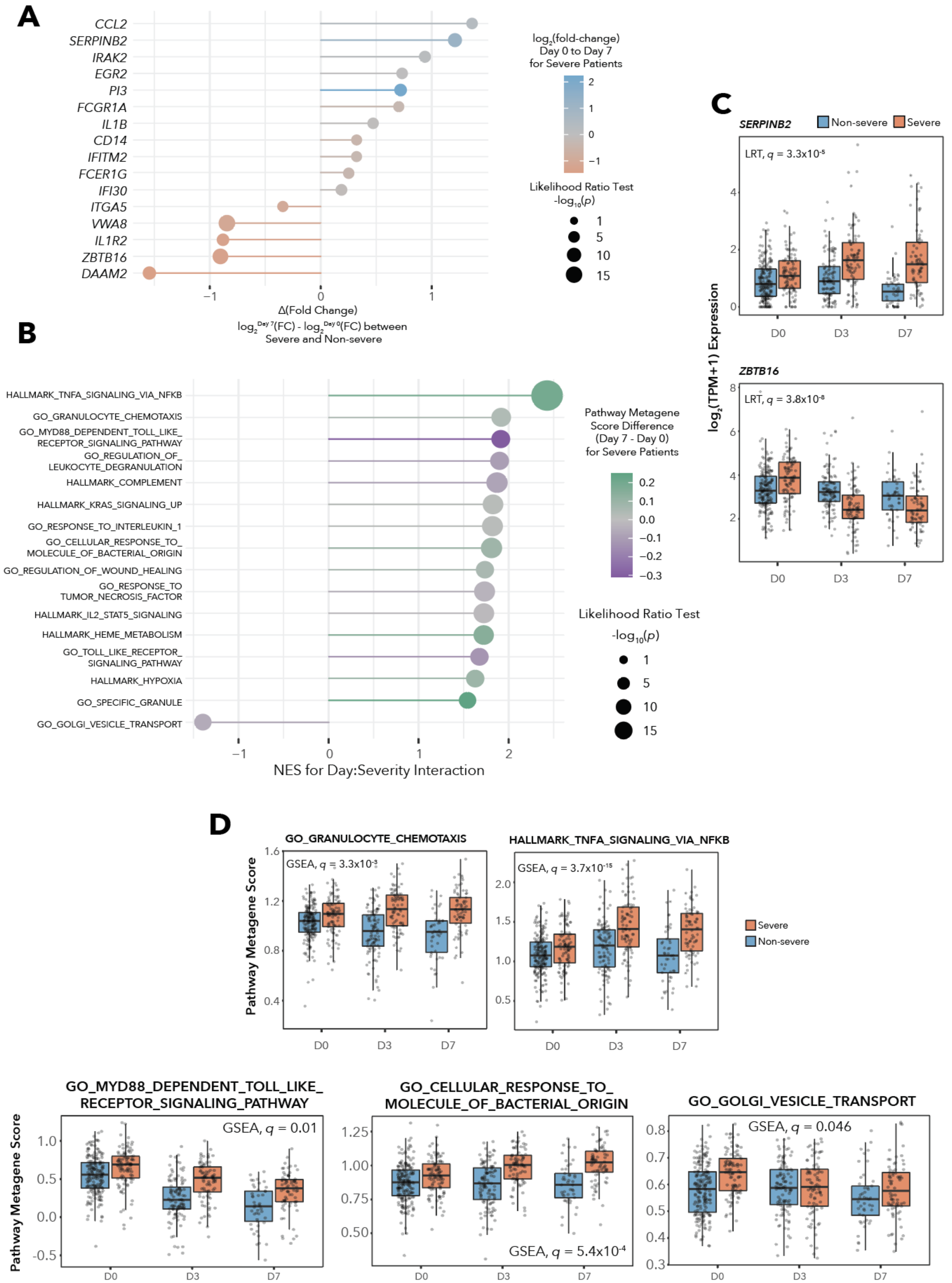
A
CCL2
SERPINB2
IRAK2
EGR2
PI3
FCGR1A
IL1B
CD14
IFITM2
FCER1G
IFI30
ITGA5
VWA8
IL1R2
ZBTB16
DAAM2
Δ(Fold Change)
log2 Day 7(FC) - log2 Day 0(FC) between
Severe and Non-severe
log2(fold-change)
Day 0 to Day 7
for Severe Patients
Likelihood Ratio Test
-log10(p)
B
HALLMARK_TNFA_SIGNALING_VIA_NFKB
GO_GRANULOCYTE_CHEMOTAXIS
GO_MYD88_DEPENDENT_TOLL_LIKE_RECEPTOR_SIGNALING_PATHWAY
GO_REGULATION_OF_LEUKOCYTE_DEGRANULATION
HALLMARK_COMPLEMENT
HALLMARK_KRAS_SIGNALING_UP
GO_RESPONSE_TO_INTERLEUKIN_1
GO_CELLULAR_RESPONSE_TO_MOLECULE_OF_BACTERIAL_ORIGIN
GO_REGULATION_OF_WOUND_HEALING
GO_RESPONSE_TO_TUMOR_NECROSIS_FACTOR
HALLMARK_IL2_STAT5_SIGNALING
HALLMARK_HEME_METABOLISM
GO_TOLL_LIKE_RECEPTOR_SIGNALING_PATHWAY
HALLMARK_HYPOXIA
GO_SPECIFIC_GRANULE
GO_GOLGI_VESICLE_TRANSPORT
NES for Day:Severity Interaction
Pathway Metagene Score Difference (Day 7 - Day 0) for Severe Patients
Likelihood Ratio Test
-log10(p)
C
SERPINB2
Non-severe
Severe
LRT, q = 3.3x10-5
ZBTB16
LRT, q = 3.8x10-8
log2(TPM+1) Expression
D0
D3
D7
D
GO_GRANULOCYTE_CHEMOTAXIS
GSEA, q = 3.3x10-3
HALLMARK_TNFA_SIGNALING_VIA_NFKB
GSEA, q = 3.7x10-15
Pathway Metagene Score
Severe
Non-severe
GO_MYD88_DEPENDENT_TOLL_LIKE_RECEPTOR_SIGNALING_PATHWAY
GSEA, q = 0.01
GO_CELLULAR_RESPONSE_TO_MOLECULE_OF_BACTERIAL_ORIGIN
GSEA, q = 5.4x10-4
GO_GOLGI_VESICLE_TRANSPORT
GSEA, q = 0.046

**Figure S12. Genes and Pathways that Vary with Time According to Severity, Related to Figure 2.**

**(A)** Selected genes which display a significant interaction between severity and time according to the Likelihood Ratio Test comparing models with or without the interaction term Day:$Severity_{Max}$. Point size indicates $-log_{10}(p)$, distance from center indicates the difference in $log_2$(fold-change) between Day 7 and Day 0 for severe versus non-severe patients, and color indicates the $log_2$(fold-change) between Day 7 and Day 0 within severe patients.

**(B)** Gene set enrichment analysis on the gene list ranked according to the Likelihood Ratio Test for the interaction term Day:$Severity_{Max}$. Point size indicates $-log_{10}(p)$, distance from center indicates NES, and color indicates the difference in the pathway metagene score between Day 7 and Day 0 within severe patients.

**(C)** Box plots of $log_2$(TPM+1) expression over time of *SERPINB2* and *ZBTB16*, two genes which show significant interactions between Day and $Severity_{Max}$ according to the DESeq2 Likelihood Ratio Test.

**(D)** Box plots displaying the interaction between day and $Severity_{Max}$ for five pathway metagene scores from (B): GO_GRANULOCYTE_CHEMOTAXIS, HALLMARK_TNFA_SIGNALING_VIA_NFKB, GO_MYD88_DEPENDENT_TOLL_LIKE_RECEPTOR_SIGNALING_PATHWAY, GO_CELLULAR_RESPONSE_TO_MOLECULE_OF_BACTERIAL_ORIGIN, and GO_GOLGI_VESICLE_TRANSPORT. Indicated FDR q values are for the GSEA performed on the gene list ranked by strength of interaction between Day and $Severity_{Max}$ according to the LRT.

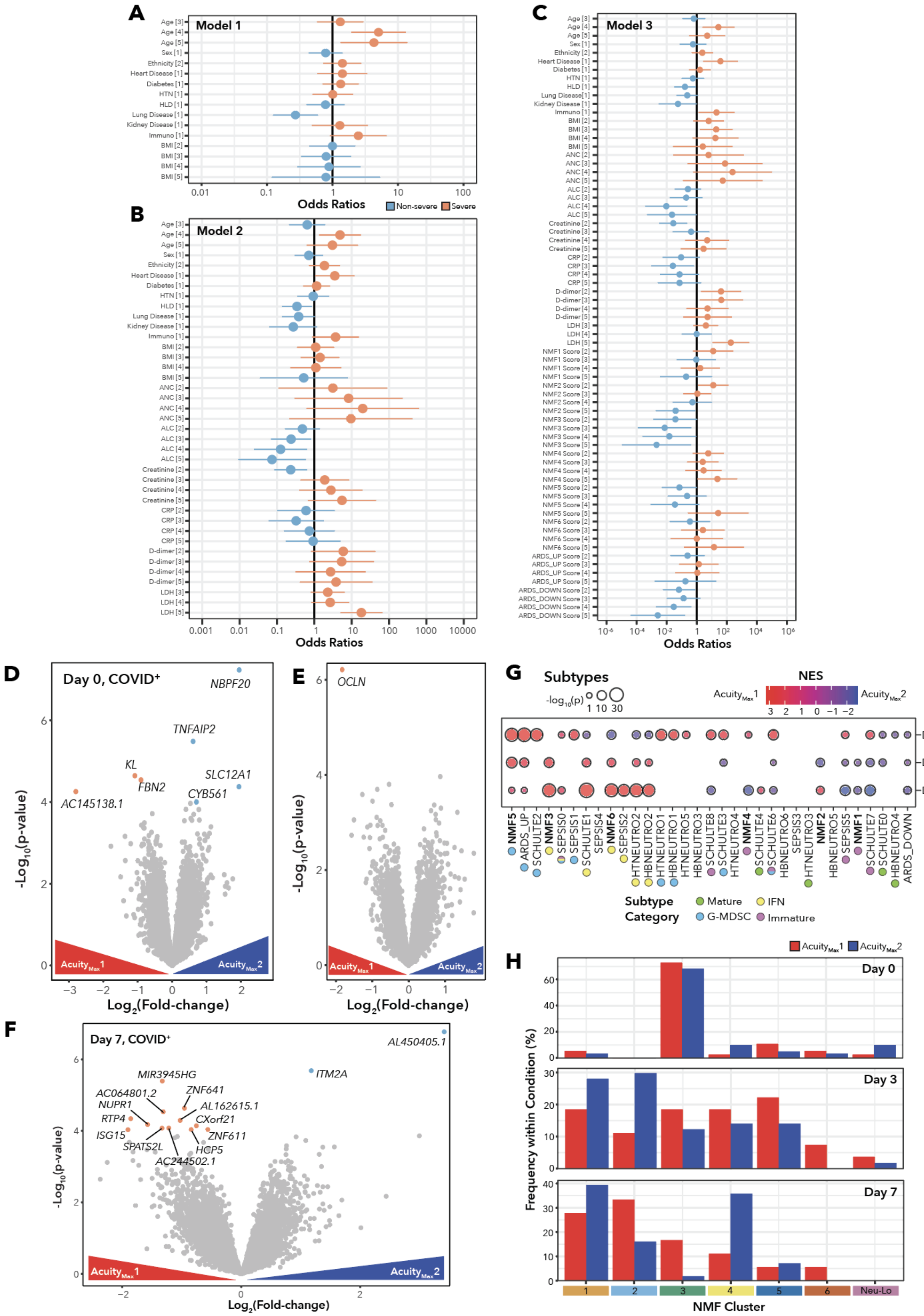
A
Model 1
Odds Ratios
Non-severe
Severe
B
Model 2
Odds Ratios
C
Model 3
Odds Ratios
D
Day 0, COVID+
NBPF20
TNFAIP2
KL
FBN2
SLC12A1
CYB561
AC145138.1
-Log10(p-value)
Log2(Fold-change)
E
OCLN
-Log10(p-value)
Log2(Fold-change)
F
Day 7, COVID+
AL450405.1
ITM2A
MIR3945HG
AC064801.2
ZNF641
NUPR1
AL162615.1
RTP4
CXorf21
ISG15
ZNF611
SPATS2L
HCP5
AC244502.1
-Log10(p-value)
Log2(Fold-change)
G
Subtypes
NES
Subtype Category
Mature
IFN
G-MDSC
Immature
H
Day 0
Day 3
Day 7
Frequency within Condition (%)
NMF Cluster
Neu-Lo

**Figure S13. Predicting COVID-19 Disease Severity on Day 0, and Comparisons Among Highest Acuity Patients, Related to Figure 3.**

**(A)-(C)** Forest plots displaying the odds ratio estimates for each factor level of the terms incorporated in the logistic regression models in Figure 3B for predicting COVID-19 disease severity on Day 0 of hospitalization. (A) includes only clinical characteristics, (B) adds clinical laboratory values, and (C) adds neutrophil gene signature scores. Lab values and gene signatures variables are broken into quintiles with factor quintile levels: 1 = lowest, 2 = low, 3 = mid, 4 = high, 5 = highest.

**(D)-(F)** Volcano plots showing differentially expressed genes between COVID-19-positive $\text{Acuity}_{\text{Max}}1$ versus $\text{Acuity}_{\text{Max}}2$ samples on (D) Day 0, (E) Day 3, and (F) Day 7. Color-coded points indicate $\log_2$(fold-change) > 0.5 and $p < 10^{-4}$.

**(G)** Gene set enrichment analysis for the differentially expressed genes between COVID-19-positive $\text{Acuity}_{\text{Max}}1$ (death) and $\text{Acuity}_{\text{Max}}2$ (intubation with survival) patients on Days 0, 3, and 7. Gene sets correspond to the neutrophil states in Figure 2F. Bubble size is scaled to $-\log_{10}$(p-value) and color corresponds to normalized enrichment score (NES).

**(H)** Bar plots comparing the distribution of NMF subtypes between COVID-19-positive $\text{Acuity}_{\text{Max}}1$ and $\text{Acuity}_{\text{Max}}2$ samples on (top) Day 0, (middle) Day 3, and (bottom) Day 7, showing frequency within a given Acuity. Fisher’s exact test did not indicate any significant differences.

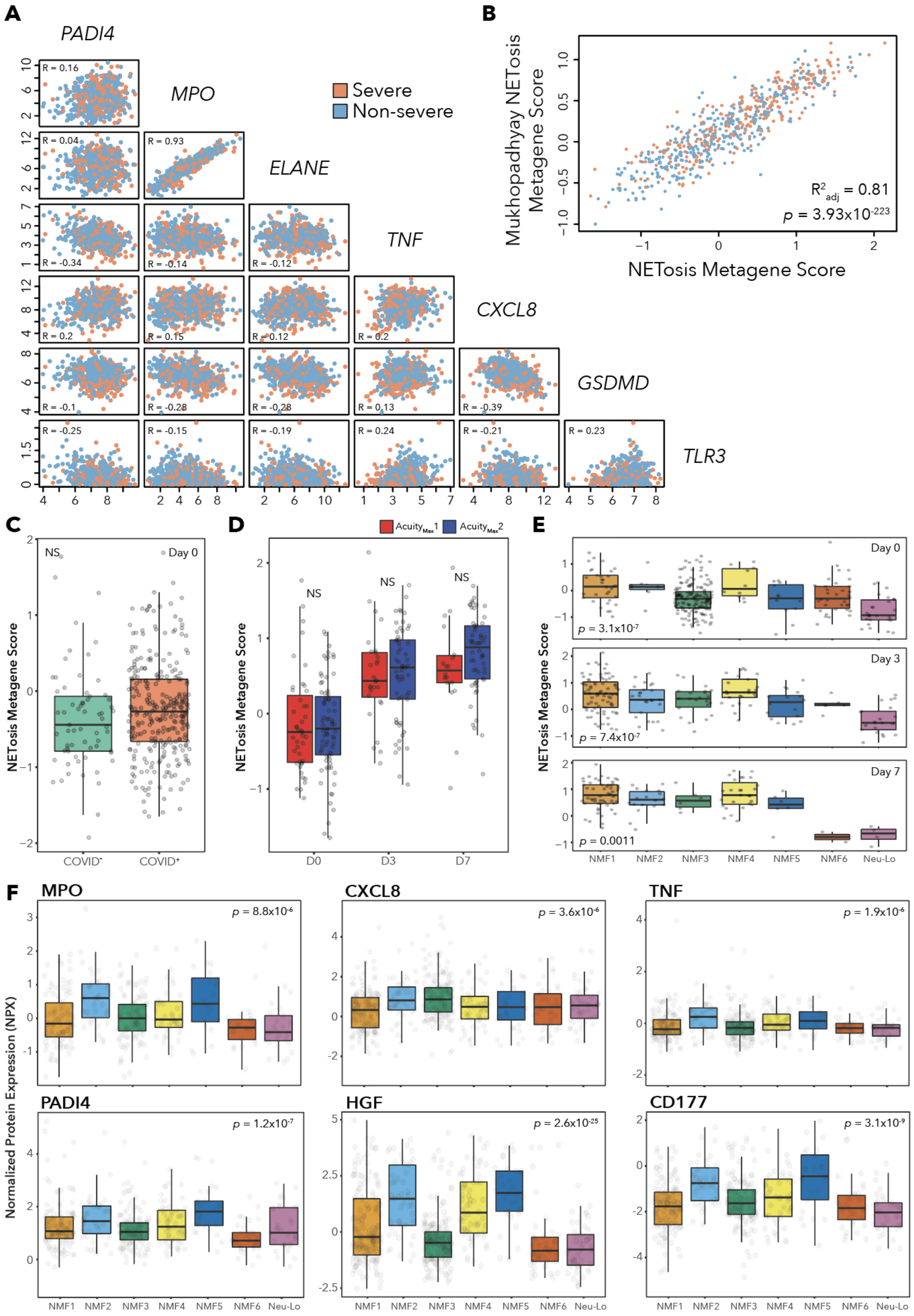
A
PADI4
MPO
ELANE
TNF
CXCL8
GSDMD
TLR3
Severe
Non-severe
R = 0.16
R = 0.04
R = 0.93
R = -0.34
R = -0.14
R = -0.12
R = 0.2
R = 0.15
R = 0.12
R = 0.2
R = -0.1
R = -0.28
R = -0.28
R = 0.13
R = -0.39
R = -0.25
R = -0.15
R = -0.19
R = 0.24
R = -0.21
R = 0.23
B
Mukhopadhyay NETosis Metagene Score
NETosis Metagene Score
$R^2_{adj}$ = 0.81
$p$ = 3.93x$10^{-223}$
C
NETosis Metagene Score
NS
Day 0
COVID⁻
COVID⁺
D
Acuity$_{Max}$1
Acuity$_{Max}$2
NETosis Metagene Score
NS
NS
NS
D0
D3
D7
E
NETosis Metagene Score
Day 0
$p$ = 3.1x$10^{-7}$
Day 3
$p$ = 7.4x$10^{-7}$
Day 7
$p$ = 0.0011
NMF1
NMF2
NMF3
NMF4
NMF5
NMF6
Neu-Lo
F
Normalized Protein Expression (NPX)
MPO
$p$ = 8.8x$10^{-6}$
CXCL8
$p$ = 3.6x$10^{-6}$
TNF
$p$ = 1.9x$10^{-6}$
PADI4
$p$ = 1.2x$10^{-7}$
HGF
$p$ = 2.6x$10^{-25}$
CD177
$p$ = 3.1x$10^{-9}$
NMF1
NMF2
NMF3
NMF4
NMF5
NMF6
Neu-Lo

**Figure S14. Patterns of NETosis in RNA and Plasma, Related to Figure 4.**
**(A)** Scatter plots comparing $log_2$(TPM+1) expression of genes contributing to the NETosis metagene score (*PADI4*, *MPO*, *ELANE*, *TNF*, *CXCL8*, *GSDMD*, *TLR3*). Points are color-coded according to $Severity_{Max}$. Pearson correlation coefficients are indicated.
**(B)** Scatter plot comparing the NETosis metagene score to a previously defined NETosis gene set from Mukhopadhyay et al. (*CR1*, *ITGAM*, *CFH*, *CFB*, *C5*, *C5AR1*, *C3*, *CFP*, *MPO*, *ELANE*, *CTSG*, *HMGB1*, *AGER*, *TLR2*, *TLR4*, *H4C1*, *TF*, *TFPI*, *F2*, *FGB*, *PLG*, *VWF*, *PF4*, *CCL5*, *DNASE1*, *ITGB2*, *CD33*, *CEACAM8*). $R^2$ and p value determined using lm() in R.
**(C)** Box plots comparing NETosis metagene score across COVID-19 status. No significant difference measured according to the Wilcoxon rank-sum test.
**(D)** Box plots comparing NETosis metagene score across time, separated by $Acuity_{Max}$1 versus $Acuity_{Max}$2. No significant difference measured according to the Wilcoxon rank-sum test.
**(E)** Box plots comparing NETosis metagene score across NMF clusters, separated by Day. Indicated p values are for the Kruskal-Wallis test.
**(F)** Box plots indicating the matched plasma protein levels in NPX for COVID-19-positive samples on Days 0, 3, and 7 across NMF clusters, for the following protein markers of NETosis (left to right): MPO, CXCL8, TNF, PADI4, HGF, CD177. Indicated p values are for the Kruskal-Wallis test.

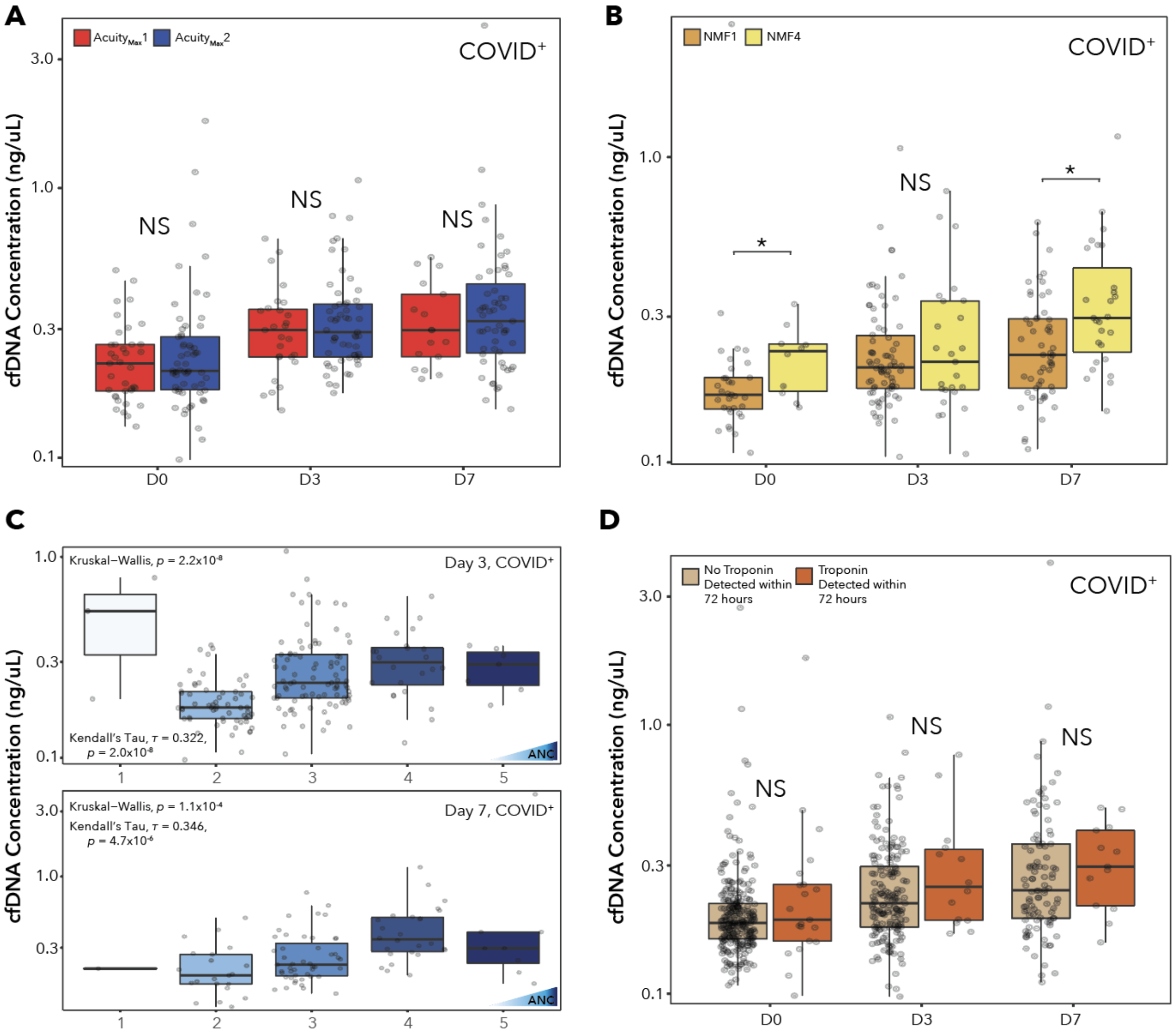
A
AcuityMax1
AcuityMax2
COVID+
cfDNA Concentration (ng/uL)
3.0
1.0
0.3
0.1
NS
NS
NS
D0
D3
D7
B
NMF1
NMF4
COVID+
cfDNA Concentration (ng/uL)
1.0
0.3
0.1
*
NS
*
D0
D3
D7
C
Kruskal–Wallis, p = 2.2x10-8
Day 3, COVID+
cfDNA Concentration (ng/uL)
1.0
0.3
0.1
Kendall's Tau, τ = 0.322,
p = 2.0x10-8
ANC
1
2
3
4
5
Kruskal–Wallis, p = 1.1x10-4
Day 7, COVID+
Kendall's Tau, τ = 0.346,
p = 4.7x10-6
3.0
1.0
0.3
ANC
1
2
3
4
5
D
No Troponin Detected within 72 hours
Troponin Detected within 72 hours
COVID+
cfDNA Concentration (ng/uL)
3.0
1.0
0.3
0.1
NS
NS
NS
D0
D3
D7

**Figure S15. Cell-free DNA in COVID-19, Related to Figure 4.**
**(A)** Box plots comparing cell-free DNA concentration in plasma from COVID-19-positive patients across time, separated by $Acuity_{Max}$1 versus $Acuity_{Max}$2. No significant differences were observed with the Wilcoxon rank-sum test.
**(B)** Box plots comparing cell-free DNA concentration in plasma from COVID-19-positive patients across time, separated by NMF1 versus NMF4. Single asterisk denotes Wilcoxon rank-sum test $p < 0.05$.
**(C)** Box plots comparing cell-free DNA concentration in plasma from COVID-19-positive patients across ANC categories for (top) Day 3 and (bottom) Day 7. The Kruskal-Wallis test was used to determine differences between groups, and ordinal correlation was measured with Kendall's tau.
**(D)** Box plots comparing cell-free DNA concentration in plasma from COVID-19-positive patients across time, separated by whether or not Troponin was detected in the plasma within 72 hours. No significant differences were observed with the Wilcoxon rank-sum test.

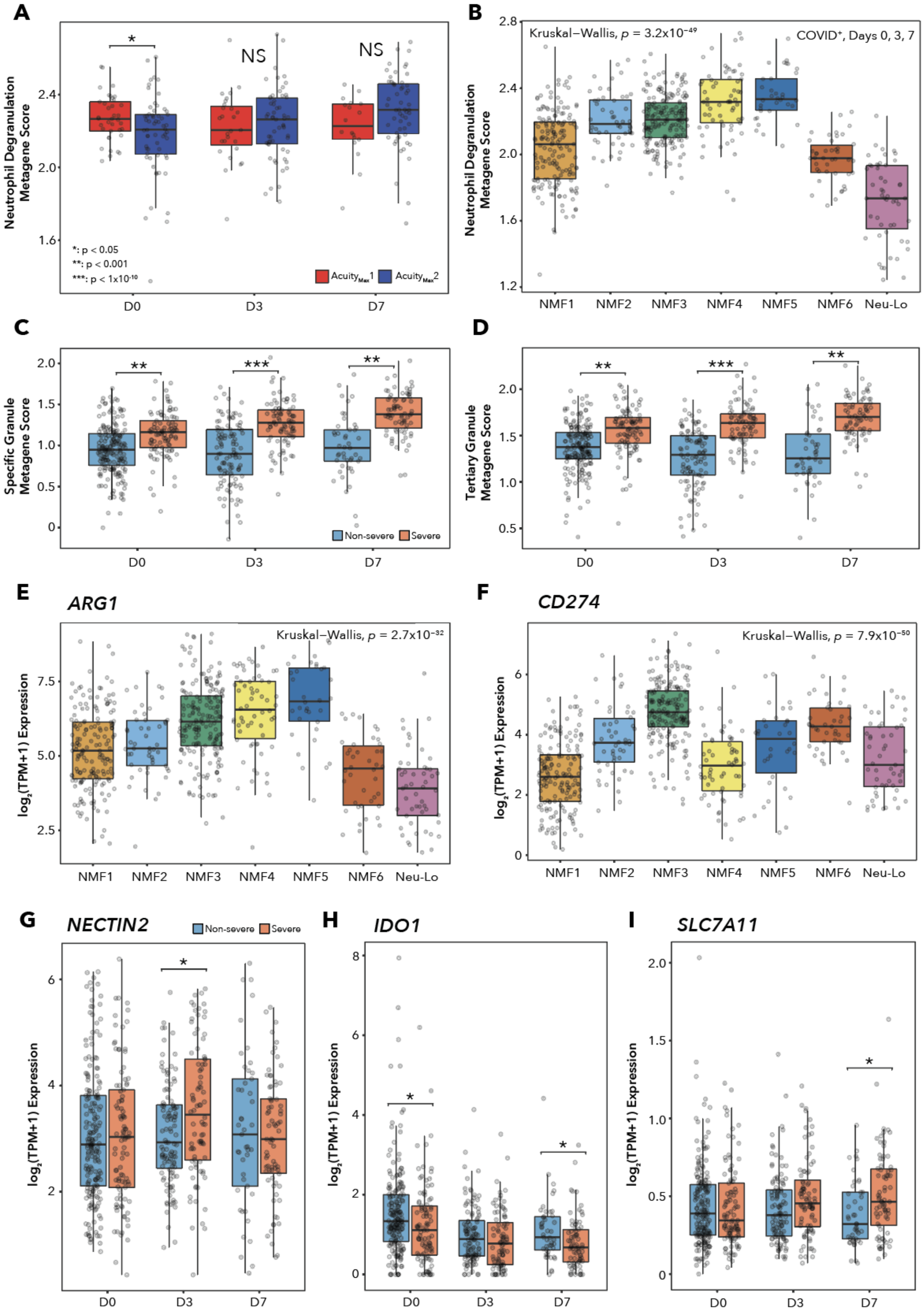
A
Neutrophil Degranulation Metagene Score
*
NS
NS
*: p < 0.05
**: p < 0.001
***: p < 1x10^-10
AcuityMax1
AcuityMax2
D0
D3
D7
B
Kruskal–Wallis, p = 3.2x10^-49
COVID+, Days 0, 3, 7
Neutrophil Degranulation Metagene Score
NMF1
NMF2
NMF3
NMF4
NMF5
NMF6
Neu-Lo
C
Specific Granule Metagene Score
**
***
**
Non-severe
Severe
D0
D3
D7
D
Tertiary Granule Metagene Score
**
***
**
D0
D3
D7
E
ARG1
Kruskal–Wallis, p = 2.7x10^-32
log2(TPM+1) Expression
NMF1
NMF2
NMF3
NMF4
NMF5
NMF6
Neu-Lo
F
CD274
Kruskal–Wallis, p = 7.9x10^-50
log2(TPM+1) Expression
NMF1
NMF2
NMF3
NMF4
NMF5
NMF6
Neu-Lo
G
NECTIN2
Non-severe
Severe
*
log2(TPM+1) Expression
D0
D3
D7
H
IDO1
*
*
log2(TPM+1) Expression
D0
D3
D7
I
SLC7A11
*
log2(TPM+1) Expression
D0
D3
D7

**Figure S16. Neutrophil Degranulation and T Cell Suppression Mechanisms in COVID-19, Related to Figure 4.**

**(A)-(B)** Box plots comparing the REACTOME_NEUTROPHIL_DEGRANULATION metagene score for COVID-19-positive patients across (A) time, separated by $Acuity_{Max}1$ versus $Acuity_{Max}2$, and (B) NMF clusters on Days 0, 3, and 7. P values refer to the Wilcoxon rank-sum test and Kruskal-Wallis test, respectively.

**(C)-(D)** Box plots comparing (C) GO_SPECIFIC_GRANULE metagene score and (D) GO_TERTIARY_GRANULE metagene score across time for COVID-19-positive patients, separated by $Severity_{Max}$. P values for Wilcoxon rank-sum test indicated according to the legend in (A).

**(E)-(F)** Box plots comparing $log_2$(TPM+1) expression of (E) *ARG1* and (F) *CD274* (gene for PD-L1) across NMF clusters for COVID-19-positive samples on Days 0, 3, and 7. Indicated p values for Kruskal-Wallis test.

**(G)-(I)** Box plots comparing $log_2$(TPM+1) expression of (G) *NECTIN2*, (H) *IDO1*, and (I) *SLC7A11* (gene for $x_c^-$) across time, separated by $Severity_{Max}$. Indicated p values for Wilcoxon rank-sum test.

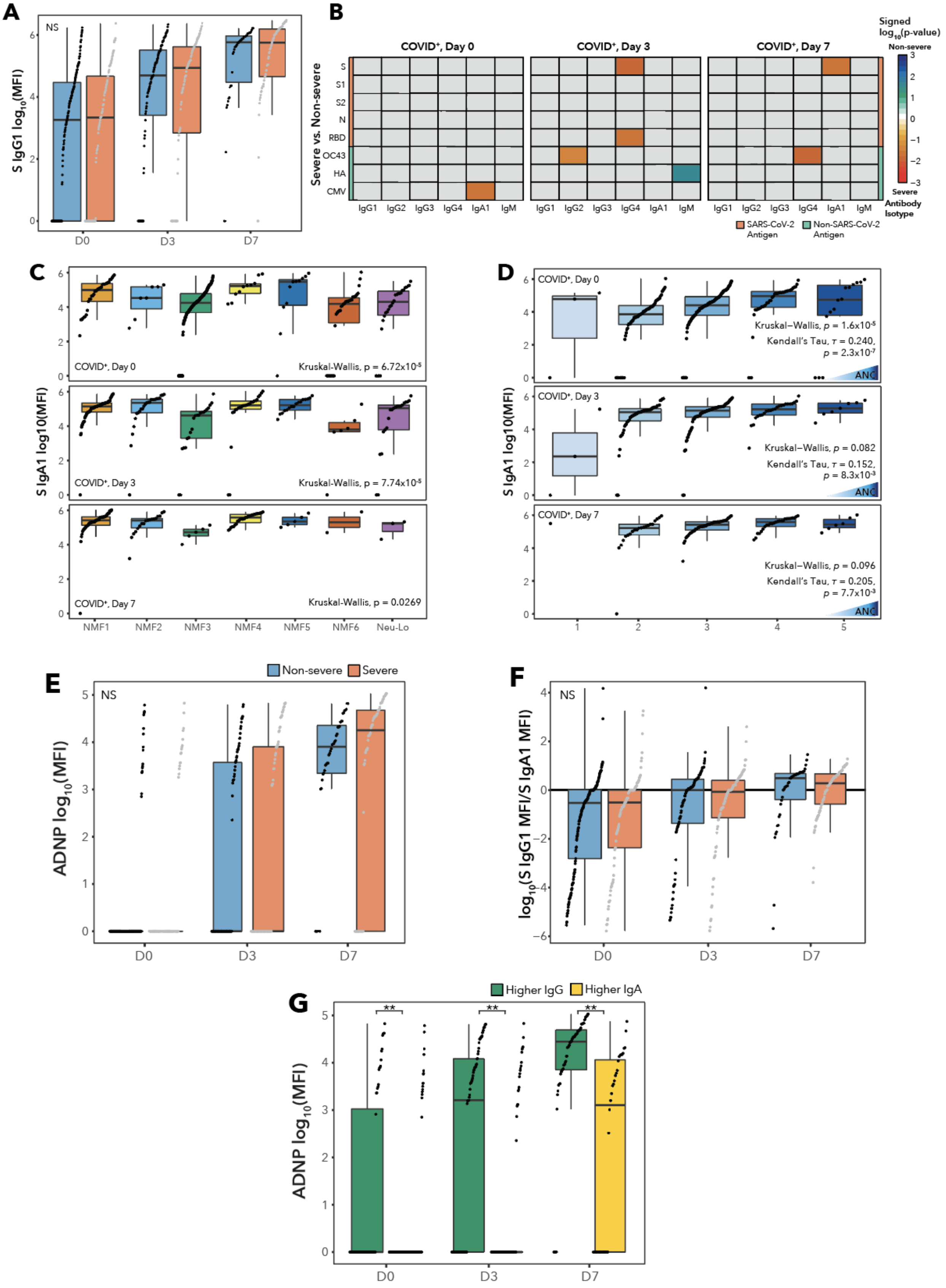
A
NS
S IgG1 $\log_{10}$(MFI)
D0
D3
D7
B
COVID⁺, Day 0
COVID⁺, Day 3
COVID⁺, Day 7
Severe vs. Non-severe
S
S1
S2
N
RBD
OC43
HA
CMV
IgG1
IgG2
IgG3
IgG4
IgA1
IgM
Signed $\log_{10}$(p-value)
Non-severe
Severe
Antibody Isotype
SARS-CoV-2 Antigen
Non-SARS-CoV-2 Antigen
C
S IgA1 log10(MFI)
COVID⁺, Day 0
Kruskal-Wallis, p = 6.72x10⁻⁵
COVID⁺, Day 3
Kruskal-Wallis, p = 7.74x10⁻⁵
COVID⁺, Day 7
Kruskal-Wallis, p = 0.0269
NMF1
NMF2
NMF3
NMF4
NMF5
NMF6
Neu-Lo
D
S IgA1 log10(MFI)
COVID⁺, Day 0
Kruskal–Wallis, p = 1.6x10⁻⁵
Kendall's Tau, τ = 0.240, p = 2.3x10⁻⁷
COVID⁺, Day 3
Kruskal–Wallis, p = 0.082
Kendall's Tau, τ = 0.152, p = 8.3x10⁻³
COVID⁺, Day 7
Kruskal–Wallis, p = 0.096
Kendall's Tau, τ = 0.205, p = 7.7x10⁻³
ANC
E
Non-severe
Severe
NS
ADNP $\log_{10}$(MFI)
D0
D3
D7
F
NS
$\log_{10}$(S IgG1 MFI/S IgA1 MFI)
D0
D3
D7
G
Higher IgG
Higher IgA
ADNP $\log_{10}$(MFI)
D0
D3
D7

**Figure S17. Immunoglobulin Levels in Plasma, and Antibody-Dependent Neutrophil Phagocytosis Assay, Related to Figure 5.**

**(A)** Box plots comparing spike protein-specific IgG1 $\log_{10}$(MFI) values in plasma across time, separated by $\text{Severity}_{\text{Max}}$. Indicated p values for Wilcoxon rank-sum test.

**(B)** Heatmaps displaying the signed (according to fold-change), log-transformed p values for the Wilcoxon rank-sum tests comparing levels of antigen-specific antibody isotypes across disease status for COVID-19-positive samples. Top row of heatmaps, comparisons between $\text{Acuity}_{\text{Max}}1$ and $\text{Acuity}_{\text{Max}}2$; bottom row of heatmaps, comparisons between Severe and Non-severe. The three main columns indicate Day. Within each heatmap, rows indicate viral antigens, color-coded by virus of origin: SARS-CoV-2 spike protein (S), SARS-CoV-2 spike protein S1 (S1), SARS-CoV-2 spike protein S2 (S2), SARS-CoV-2 nucleocapsid (N), SARS-CoV-2 receptor-binding domain (RBD), Human coronavirus OC43 (OC43), influenza hemagglutinin (HA), and cytomegalovirus (CMV). Within each heatmap, columns indicate antibody isotypes: IgG1, IgG2, IgG3, IgG4, IgA1, IgM.

**(C)** Box plots comparing S-specific IgA1 $\log_{10}$(MFI) values in plasma for COVID-19-positive samples, separated by matched NMF cluster and Day. P values are for the Kruskal-Wallis test.

**(D)** Box plots comparing S-specific IgA1 $\log_{10}$(MFI) values in plasma for COVID-19-positive samples, separated by ANC quintiles and Day. P values for global differences between quintiles are determined using the Kruskal-Wallis test, and correlations and their significance are determined using Kendall's tau.

**(E)** Box plots of background-corrected ADNP $\log_{10}$(MFI) values separated by $\text{Severity}_{\text{Max}}$ and Day. P values determined according to the Wilcoxon rank-sum test.

**(F)** Box plot depicting the log-transformed ratio of the spike (S) protein-specific IgG1 MFI to the S-specific IgA1 MFI, separated by $\text{Severity}_{\text{Max}}$ and Day. Positive values indicate a ratio in favor of IgG1, while negative values indicate a ratio in favor of IgA1. P values are for the Wilcoxon rank-sum test.

**(G)** Box plots of background-corrected ADNP $\log_{10}$(MFI) values for COVID-19-positive patients on Days 0, 3, and 7, separated by IgG to IgA ratios. P values are for the Wilcoxon rank-sum test.

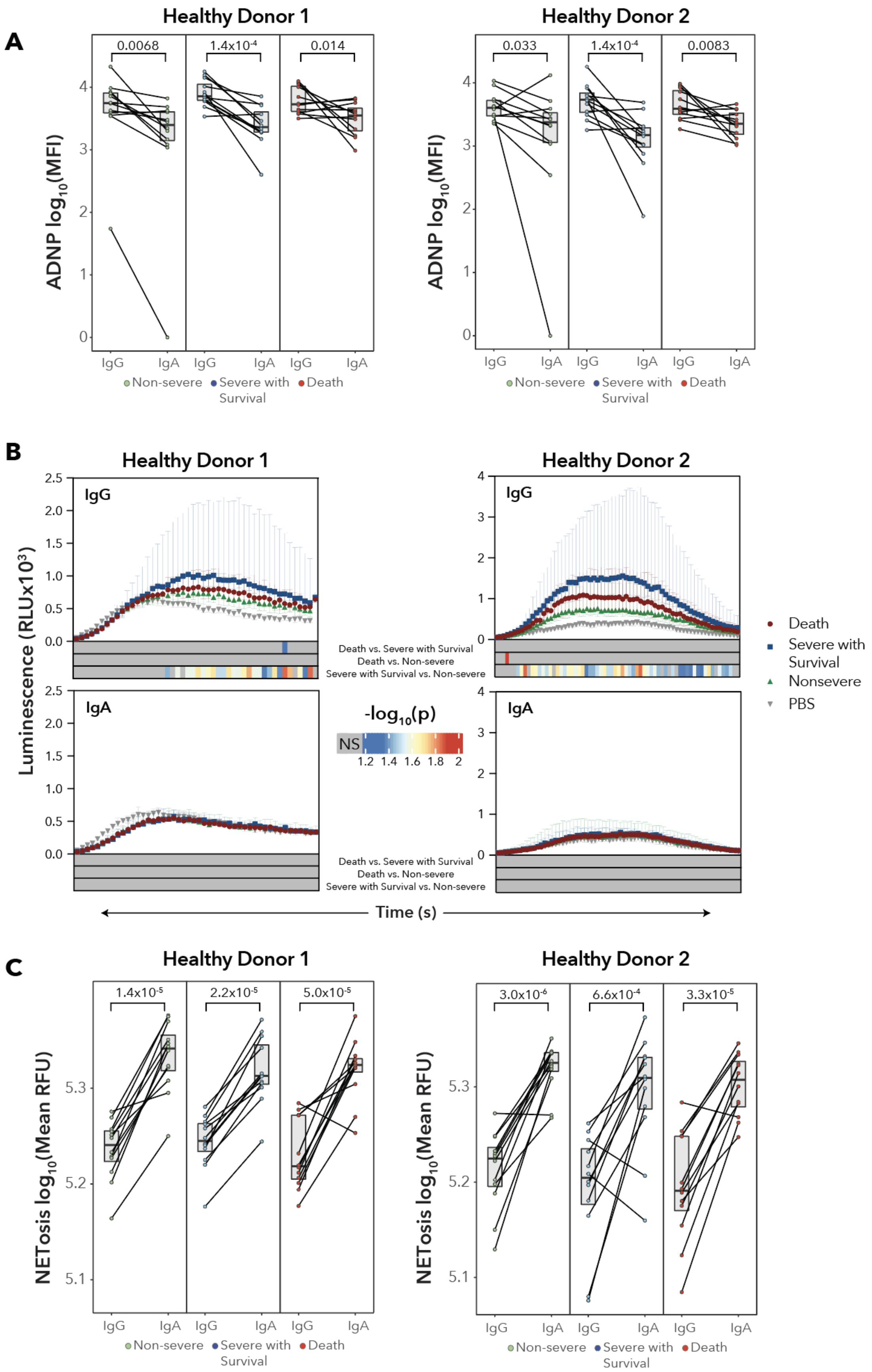
A
Healthy Donor 1
Healthy Donor 2
0.0068
1.4x10-4
0.014
0.033
1.4x10-4
0.0083
ADNP log10(MFI)
IgG
IgA
Non-severe
Severe with Survival
Death
B
Luminescence (RLUx10³)
Death vs. Severe with Survival
Death vs. Non-severe
Severe with Survival vs. Non-severe
-log10(p)
NS
1.2 1.4 1.6 1.8 2
Death
Severe with Survival
Nonsevere
PBS
Time (s)
C
1.4x10-5
2.2x10-5
5.0x10-5
3.0x10-6
6.6x10-4
3.3x10-5
NETosis log10(Mean RFU)

**Figure S18. Differential Effects of IgG versus IgA Antibodies on Neutrophil Effector Functions, Related to Figure 5.**

**(A)** Paired-line graphs of ADNP $\log_{10}$(MFI) values showing the effect of isolated SARS-CoV-2 S-specific IgG or IgA antibodies from serum of patients who died (n = 12), patients with severe disease who survived (n = 12), and patients with non-severe disease (n = 12). P values are for the Wilcoxon rank-sum test. Measurements using neutrophils from the two healthy donors are shown separately.

**(B)** Point-range plots showing the luminescence of the reactive oxygen species reagent, luminol, over time when neutrophils from two healthy donors are exposed to IgG:S or IgA:S immune complexes using the same purified IgG and IgA antibodies as (A) or PBS. Point ranges are plotted as median +/- interquartile range. Color bar beneath each plot displays the log-transformed P values for the Wilcoxon rank-sum test between (top) Death vs. Severe with survival, (middle) Death vs. Non-severe, and (bottom) Severe with survival vs. Non-severe values at each time point, with gray values indicating no significant difference.

**(C)** Paired-line graphs of mean SYTOX Green Nucleic Acid Stain $\log_{10}$(RFU) (quantification of NETosis) from neutrophils from two healthy donors when exposed to free antibodies from the same batch of IgG or IgA purification as in (A). P values are for the Wilcoxon rank-sum test. Measurements from the two healthy donors are shown separately.

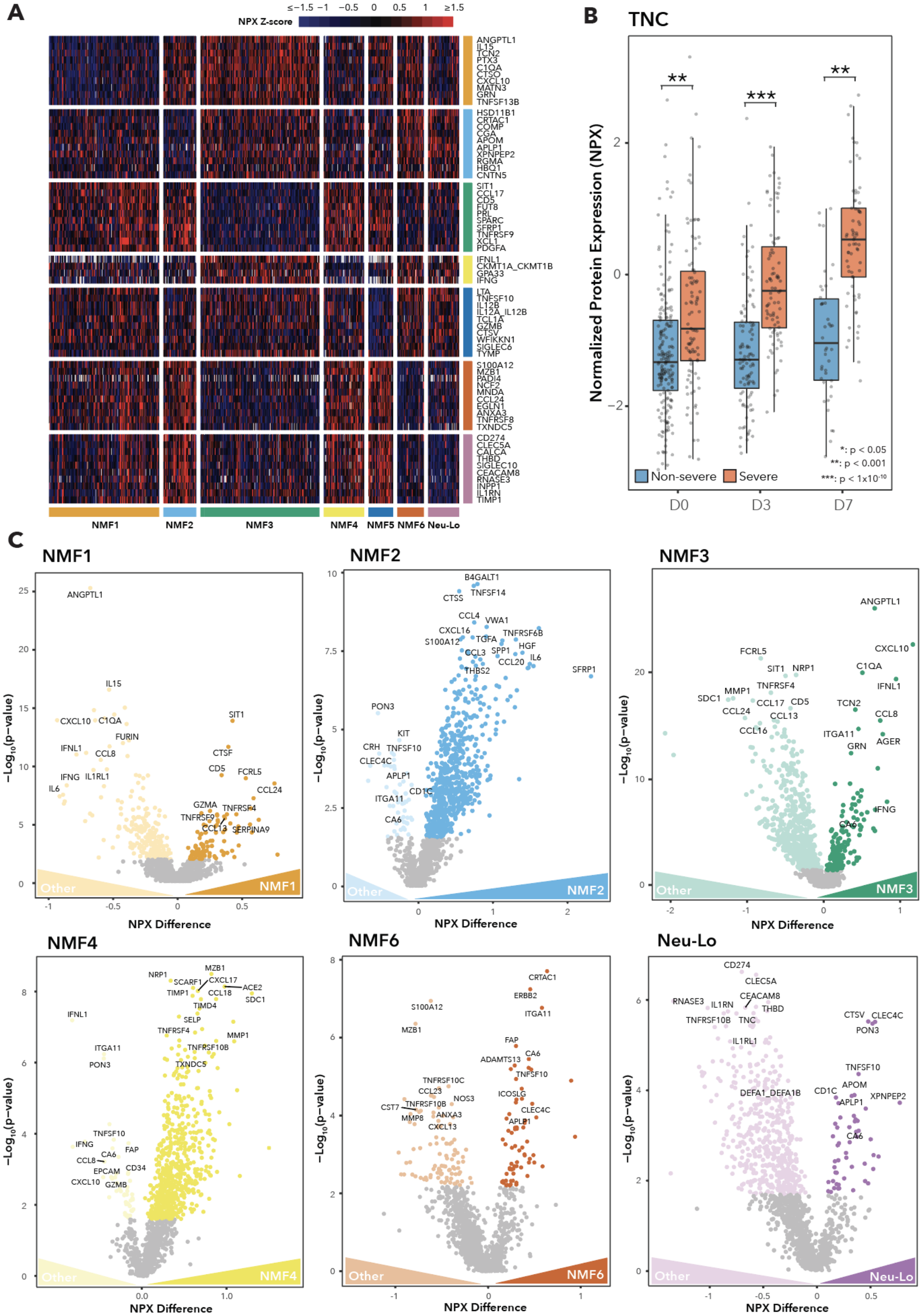

A
NPX Z-score
≤−1.5 −1 −0.5 0 0.5 1 ≥1.5
ANGPTL1
IL15
TCN2
PTX3
C1QA
CTSO
CXCL10
MATN3
GRN
TNFSF13B
HSD11B1
CRTAC1
COMP
CGA
APOM
APLP1
XPNPEP2
RGMA
HBQ1
CNTN5
SIT1
CCL17
CD5
FUT8
PRL
SPARC
SFRP1
TNFRSF9
XCL1
PDGFA
IFNL1
CKMT1A_CKMT1B
GPA33
IFNG
LTA
TNFSF10
IL12B
IL12A_IL12B
TCL1A
GZMB
CTSV
WFIKKN1
SIGLEC6
TYMP
S100A12
MZB1
PADI4
NCF2
MNDA
CCL24
EGLN1
ANXA3
TNFRSF8
TXNDC5
CD274
CLEC5A
CALCA
THBD
SIGLEC10
CEACAM8
RNASE3
INPP1
IL1RN
TIMP1
NMF1
NMF2
NMF3
NMF4
NMF5
NMF6
Neu-Lo
B
TNC
Normalized Protein Expression (NPX)
**
***
**
Non-severe
Severe
*: p < 0.05
**: p < 0.001
***: p < 1x10^-10
D0
D3
D7
C
NMF1
NMF2
NMF3
NMF4
NMF6
Neu-Lo
−Log10(p−value)
NPX Difference
Other

**Figure S19. Differentially Expressed Plasma Proteins Associated with Neutrophil States, Related to Figure 6.**
**(A)** Heatmap displaying scaled expression values for negative protein markers in matched plasma of neutrophil NMF clusters for all COVID-19-positive samples.
**(B)** Box plot comparing NPX values for the TNC protein, the strongest marker of NMF5, separated by Day and severity. P values determined by the Wilcoxon rank-sum test, with asterisks denoting significance levels in the key.
**(C)** Volcano plots showing differentially expressed proteins between a given NMF cluster and all other clusters for all COVID-19-positive samples. Color coded points are significant following FDR correction.

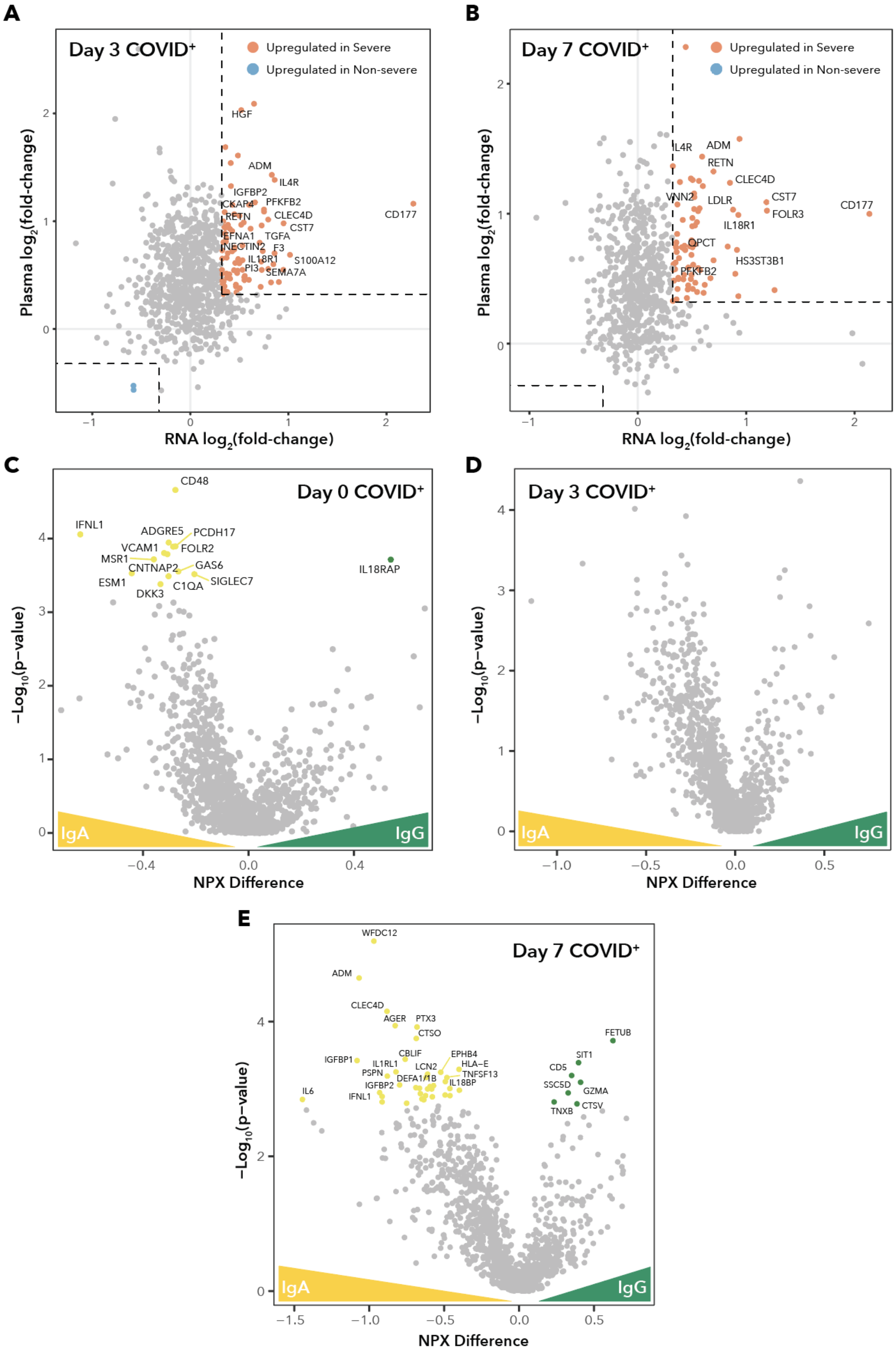
A
Day 3 COVID+
Upregulated in Severe
Upregulated in Non-severe
HGF
ADM
IL4R
IGFBP2
CKAP4
PFKFB2
RETN
CLEC4D
CD177
CST7
EFNA1
TGFA
NECTIN2
F3
IL18R1
S100A12
PI3
SEMA7A
Plasma log2(fold-change)
RNA log2(fold-change)
B
Day 7 COVID+
Upregulated in Severe
Upregulated in Non-severe
IL4R
ADM
RETN
CLEC4D
VNN2
LDLR
CST7
FOLR3
CD177
IL18R1
QPCT
HS3ST3B1
PFKFB2
Plasma log2(fold-change)
RNA log2(fold-change)
C
Day 0 COVID+
CD48
IFNL1
ADGRE5
PCDH17
VCAM1
FOLR2
MSR1
CNTNAP2
GAS6
ESM1
SIGLEC7
C1QA
DKK3
IL18RAP
IgA
IgG
−Log10(p−value)
NPX Difference
D
Day 3 COVID+
IgA
IgG
−Log10(p−value)
NPX Difference
E
Day 7 COVID+
WFDC12
ADM
CLEC4D
AGER
PTX3
CTSO
CBLIF
EPHB4
IGFBP1
IL1RL1
LCN2
HLA−E
PSPN
DEFA1/1B
TNFSF13
IGFBP2
IL18BP
IFNL1
IL6
FETUB
SIT1
CD5
SSC5D
GZMA
TNXB
CTSV
IgA
IgG
−Log10(p−value)
NPX Difference

**Figure S20. RNA and Protein Log-Fold-Change Comparisons for COVID-19 status and Severity Across Days, and Differentially-expressed Plasma Proteins Associated with Higher IgG or IgA Titers, Related to Figure 6.**
**(A)-(B)** Scatter plots comparing the $log_2$(fold-change) between severe and non-severe samples on (A) Day 3 and (B) Day 7 for neutrophil RNA-Seq with the $log_2$(fold-change) of the NPX difference of the corresponding protein in the plasma. Corrections were made for the clinical covariates of age, sex, ethnicity, heart disease, diabetes, hypertension, hyperlipidemia, pulmonary disease, kidney disease, immunocompromised status. Color-coded points indicate $log_2$ fold-change > 1.25 in both mRNA and protein.
**(C)-(E)** Volcano plots displaying differentially expressed genes between COVID-19-positive samples with either higher IgA1-to-IgG1 ratios or higher IgG1-to-IgA1 ratios on (C) Day 0, (D) Day 3, and (E) Day 7. Color-coded points have FDR q-values < 0.05.

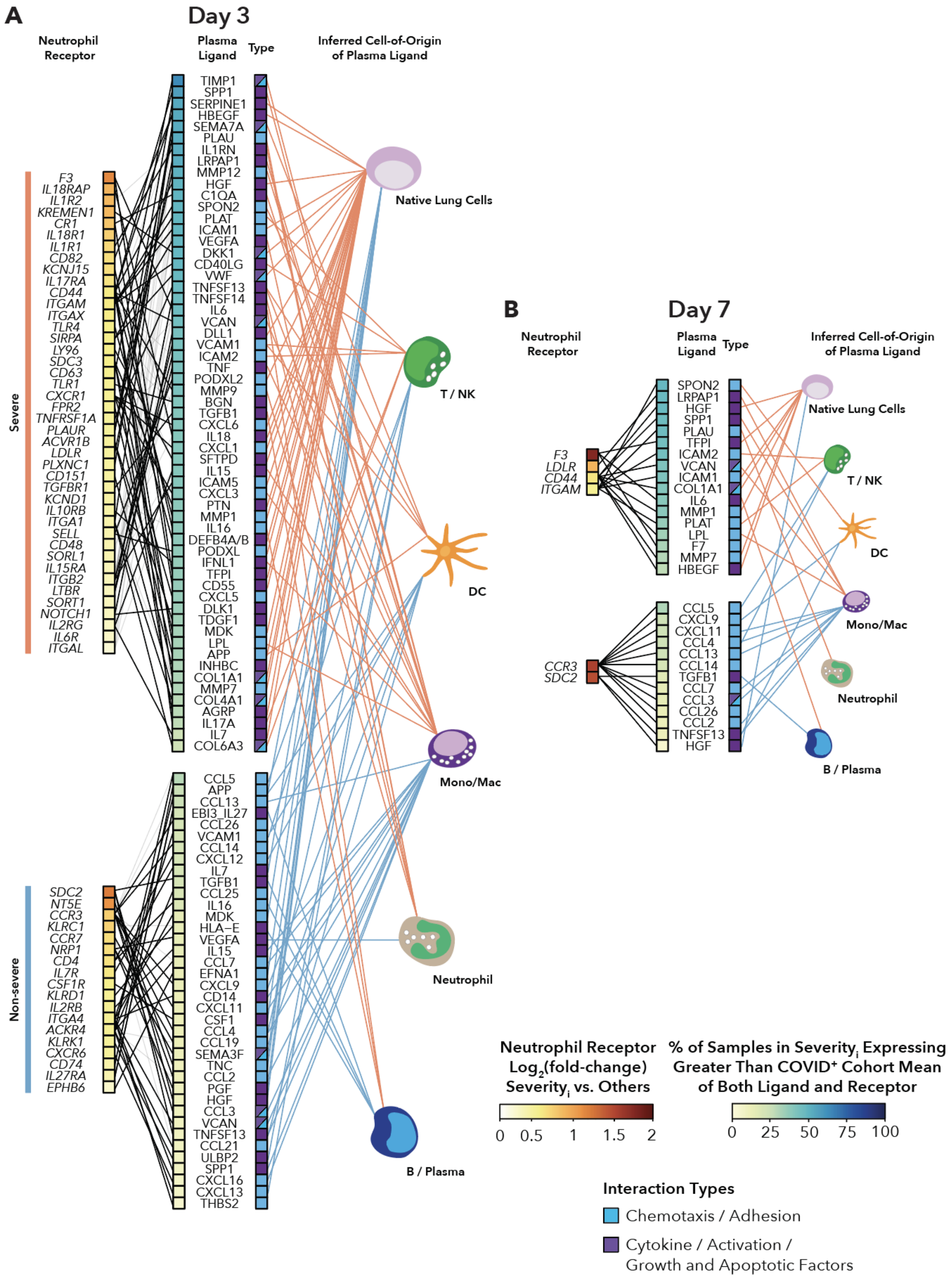
A
Day 3
Neutrophil Receptor
Plasma Ligand
Type
Inferred Cell-of-Origin of Plasma Ligand
Severe
F3
IL18RAP
IL1R2
KREMEN1
CR1
IL18R1
IL1R1
CD82
KCNJ15
IL17RA
CD44
ITGAM
ITGAX
TLR4
SIRPA
LY96
SDC3
CD63
TLR1
CXCR1
FPR2
TNFRSF1A
PLAUR
ACVR1B
LDLR
PLXNC1
CD151
TGFBR1
KCND1
IL10RB
ITGA1
SELL
CD48
SORL1
IL15RA
ITGB2
LTBR
SORT1
NOTCH1
IL2RG
IL6R
ITGAL
TIMP1
SPP1
SERPINE1
HBEGF
SEMA7A
PLAU
IL1RN
LRPAP1
MMP12
HGF
C1QA
SPON2
PLAT
ICAM1
VEGFA
DKK1
CD40LG
VWF
TNFSF13
TNFSF14
IL6
VCAN
DLL1
VCAM1
ICAM2
TNF
PODXL2
MMP9
BGN
TGFB1
CXCL6
IL18
CXCL1
SFTPD
IL15
ICAM5
CXCL3
PTN
MMP1
IL16
DEFB4A/B
PODXL
IFNL1
TFPI
CD55
CXCL5
DLK1
TDGF1
MDK
LPL
APP
INHBC
COL1A1
MMP7
COL4A1
AGRP
IL17A
IL7
COL6A3
Non-severe
SDC2
NT5E
CCR3
KLRC1
CCR7
NRP1
CD4
IL7R
CSF1R
KLRD1
IL2RB
ITGA4
ACKR4
KLRK1
CXCR6
CD74
IL27RA
EPHB6
CCL5
APP
CCL13
EBI3_IL27
CCL26
VCAM1
CCL14
CXCL12
IL7
TGFB1
CCL25
IL16
MDK
HLA-E
VEGFA
IL15
CCL7
EFNA1
CXCL9
CD14
CXCL11
CSF1
CCL4
CCL19
SEMA3F
TNC
CCL2
PGF
HGF
CCL3
VCAN
TNFSF13
CCL21
ULBP2
SPP1
CXCL16
CXCL13
THBS2
Native Lung Cells
T / NK
DC
Mono/Mac
Neutrophil
B / Plasma
B
Day 7
Neutrophil Receptor
Plasma Ligand
Type
Inferred Cell-of-Origin of Plasma Ligand
F3
LDLR
CD44
ITGAM
SPON2
LRPAP1
HGF
SPP1
PLAU
TFPI
ICAM2
VCAN
ICAM1
COL1A1
IL6
MMP1
PLAT
LPL
F7
MMP7
HBEGF
CCR3
SDC2
CCL5
CXCL9
CXCL11
CCL4
CCL13
CCL14
TGFB1
CCL7
CCL3
CCL26
CCL2
TNFSF13
HGF
Native Lung Cells
T / NK
DC
Mono/Mac
Neutrophil
B / Plasma
Neutrophil Receptor Log2(fold-change) Severityi vs. Others
0 0.5 1 1.5 2
% of Samples in Severityi Expressing Greater Than COVID+ Cohort Mean of Both Ligand and Receptor
0 25 50 75 100
Interaction Types
Chemotaxis / Adhesion
Cytokine / Activation / Growth and Apoptotic Factors

**Figure S21. Ligand-receptor Interaction Analysis for Severe versus Non-severe Patients, Related to Figure 7.**

Ligand-receptor analysis for differentially expressed ligands in plasma and receptors on neutrophils between COVID-19-positive severe and non-severe samples on (A) Day 3 and (B) Day 7. Ligands and receptors are color-coded by severity. Receptors are color-scaled according to the $\log_2$(fold-change) between severity groups. Ligands are color-scaled according to the percentage of samples within the severity group for which the ligand and receptor are both expressed above the overall mean expression.

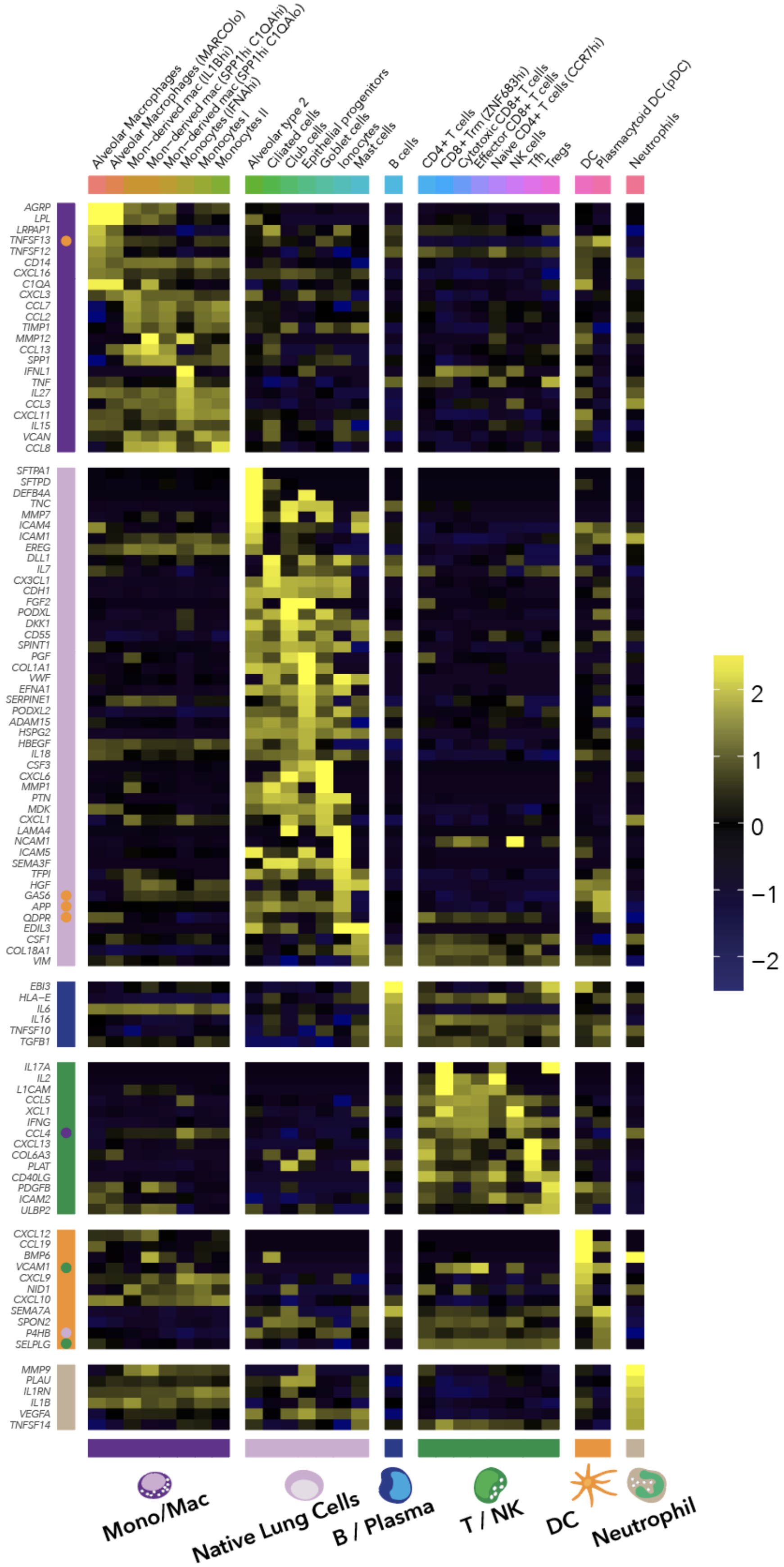
Alveolar Macrophages
Alveolar Macrophages (MARCOlo)
Mon−derived mac (IL1Bhi)
Mon−derived mac (SPP1hi C1QAhi)
Mon−derived mac (SPP1hi C1QAlo)
Monocytes (IFNAhi)
Monocytes I
Monocytes II
Alveolar type 2
Ciliated cells
Club cells
Epithelial progenitors
Goblet cells
Ionocytes
Mast cells
B cells
CD4+ T cells
CD8+ Trm (ZNF683hi)
Cytotoxic CD8+ T cells
Effector CD8+ T cells
Naive CD4+ T cells (CCR7hi)
NK cells
Tfh
Tregs
DC
Plasmacytoid DC (pDC)
Neutrophils
AGRP
LPL
LRPAP1
TNFSF13
TNFSF12
CD14
CXCL16
C1QA
CXCL3
CCL7
CCL2
TIMP1
MMP12
CCL13
SPP1
IFNL1
TNF
IL27
CCL3
CXCL11
IL15
VCAN
CCL8
SFTPA1
SFTPD
DEFB4A
TNC
MMP7
ICAM4
ICAM1
EREG
DLL1
IL7
CX3CL1
CDH1
FGF2
PODXL
DKK1
CD55
SPINT1
PGF
COL1A1
VWF
EFNA1
SERPINE1
PODXL2
ADAM15
HSPG2
HBEGF
IL18
CSF3
CXCL6
MMP1
PTN
MDK
CXCL1
LAMA4
NCAM1
ICAM5
SEMA3F
TFPI
HGF
GAS6
APP
QDPR
EDIL3
CSF1
COL18A1
VIM
EBI3
HLA−E
IL6
IL16
TNFSF10
TGFB1
IL17A
IL2
L1CAM
CCL5
XCL1
IFNG
CCL4
CXCL13
COL6A3
PLAT
CD40LG
PDGFB
ICAM2
ULBP2
CXCL12
CCL19
BMP6
VCAM1
CXCL9
NID1
CXCL10
SEMA7A
SPON2
P4HB
SELPLG
MMP9
PLAU
IL1RN
IL1B
VEGFA
TNFSF14
2
1
0
−1
−2
Mono/Mac
Native Lung Cells
B / Plasma
T / NK
DC
Neutrophil

**Figure S22. Inferring Cell-of-origin for Plasma Ligands Utilizing Single-cell RNA-seq Data, Related to Figure 7.**

Heatmap displaying single-cell RNA-sequencing (scRNA-seq) average scaled expression values per cell type for the genes encoding the protein ligands found to be differentially expressed in plasma between NMF clusters or severity groups for the ligand-receptor analysis in Figure 7 and Figure S21. scRNA-seq data is from bronchoalveolar lavage fluid (Bost et al. 2020). Column breaks indicate major cell lineages (Mono/Mac, Native lung cells, B/Plasma, T/NK, DC, Neutrophil). Row breaks indicate which genes have the highest average expression in a given major cell lineage. Color-coded dots indicate that the highest-to-second-highest difference in average scaled expression was less than 0.1, and thus the ligand was assigned to both lineages.
